# Allosteric regulation of glycogen phosphorylase solution phase structural dynamics at high spatial resolution

**DOI:** 10.1101/654665

**Authors:** Monika Kish, Victoria Smith, Sivaraman Subramanian, Frank Vollmer, Natasha Lethbridge, Lindsay Cole, Nicholas. J. Bond, Jonathan J. Phillips

## Abstract

Glycogen phosphorylase (GlyP) was the first allosteric enzyme to be described. Yet, the precise dynamic changes in solution phase structure and stability that underpin functional regulation have remained elusive. We have developed a new fully-automated and highly flexible implementation of hydrogen/deuterium-exchange mass spectrometry, operating in the millisecond regime. This enabled measurements of the solution phase local structural dynamics involved in allosteric regulation of GlyP. Here, we quantify GlyP structural dynamics in solution, describing correlated changes in structure in the activated (pSer14) and inhibited (glucose-6-phosphate bound) forms of the enzyme. The sensitivity of these measurements discerned that the 250s’ loop is natively disordered in the apo T-state, adopting a more ordered conformation in the active state. The quantitative change in stability of the 280s loop is identified, providing the first direct evidence of the entropic switch that sterically regulates substrate access to the active site.

**Significance Statement:** We have developed a new fully-automated and highly flexible implementation of hydrogen/deuterium-exchange mass spectrometry, operating in the millisecond regime. Measurements of glycogen phosphorylase quantify the solution phase stability of local structure at near-amino acid structural resolution and with no appreciable lower limit of stability. This uncovered the highly-resolved local alterations in stability which provides direct evidence of the entropic mechanism by which access to the active site is gated by the 280s loop.

**Footnotes:** Author contributions: M.K., V.S., S.S., N.L., F.V., N.B., L.C. and J.J.P. designed research; M.K., V.S., S.S., L.C. and J.J.P. performed research; M.K., V.S., S.S., L.C. and J.J.P. analyzed data; and M.K. and J.J.P. wrote the manuscript.

## Introduction

Almost 60 years ago, the term ‘allostery’ was coined by Monod and Jacob to describe structural and functional regulation of proteins by ‘non-steric’ means. Yet, the molecular mechanisms for this have remained unclear and hotly debated, with two hypotheses emerging: conformer selection (Monod, Wyman and Changeux) and induced fit (Koshland). Conformer selection is widely observed, where an effector acts by altering the equilibrium (stability) between the active (R) and inactive (T) states. The archetypal allosteric enzyme is glycogen phosphorylase (GlyP) and much insight has been gained from high-resolution structures of trapped R/T-states of GlyP, with seminal work by Barford and co-workers (1), but also from low-resolution studies of enzyme kinetics (2). It remains one of the most widely regulated proteins known, with multiple distinct binding sites for effectors, oligomerization and covalent modification (phosphorylation at Ser14) all contributing to functional modulation.

Now, one open question regarding functional control of GlyP is reflected across much of enzymology: What are the quantitative changes in local stability (ΔΔG) between R/T-states in solution that underpin allosteric conformer selection (3-5)? Although GlyP was the first allosteric enzyme to be identified, this is a pivotal and timely question as it is an important and proven therapeutic target for patients with type II diabetes (6), cancers (7) and neurodegenerative diseases (8). Structural biology efforts have revealed a great deal of detail of the alternative postures adopted by GlyP: significantly, some crystallographic models indicated missing density in the 280s loop in the R-state (9GPB.pdb), which condenses to a defined conformation in the T-state (1GPB.pdb) (9, 10). This has been interpreted qualitatively as the critical structural feature to gate catalytic activity – sterically regulating substrate access to the active site. It remains ambiguous the nature of such a key regulatory mechanism and here we sought to build a quantitative description of this from measurements of the GlyP dimer in solution.

It has been challenging to provide quantitative rationales for the structural switching that is observed in alternately regulated conformations of enzymes, particularly when the molecular size and complexity are large. One such method that can resolve local (near amino acid) perturbations in stability in arbitrarily large proteins is hydrogen/deuterium-exchange mass spectrometry (HDX-MS) (11-16). Moreover, these measurements are made in solution without the need to trap specific conformers, which has been used to answer questions of epitope mapping (17), protein-drug binding (18), protein-protein interactions (19), aggregation (20), effects of mutation (12), and allosteric regulation (21).

Current state-of-the-art HDX-MS systems reproducibly label protein with deuterium during labeling incubation times of approximately 30 seconds or longer, commonly using CTC PAL robotics for automation (22). Recently, there has been growing interest in measuring HDX at the sub-second time-scale to address our inability to measure the structural dynamics and stability of classes of molecule that are currently intractable (including peptide hormones, neurotransmitters and intrinsically disordered proteins/regions) (22, 23). A small number of academic laboratories have reported experimental systems and approaches to obtain millisecond HDX data (24, 25): these include microfluidic chips (26-31), a quench flow apparatus (16, 32) employed for analysis of disordered proteins, a completely online quench flow setup (11, 33) and a capillary mixer for characterizing biological reactions (34, 35).

Here, we describe the construction, validation and implementation of an online flow mixing and quenching system, termed ‘ms2min’ (to reflect a milliseconds to minutes deuterium-labeling capability). With this system we achieved reproducible and repeatable H/D exchange between 50 ms and five minutes. The capability of this instrument to measure accurate, fully-automated and flexible labeling HDX data was determined by studying the stability of synthetic peptides. This validated the approach which was then applied to build a quantitative map of GlyP stability under allosteric activation and inhibition, in solution. To the best of our knowledge, this shows for the first time the precise changes in local stability in response to allosteric modulation, notably in the order/disorder transition of the 280s loop that gates access to the active site.

## Results

### Fully-automated and flexible pulse-labeling hydrogen/deuterium-exchange mass spectrometry with millisecond precision

Our goal was to quantify the structural switch in solution, along with other coincident dynamic changes in structure between activated (pSer14) and inhibited (glucose-6-phosphate bound) forms of GlyP. To enable this, we developed a fully automated ‘bottom-up’ HDX-MS approach capable of accurate quantitative assessment of peptide stability. The newly developed ms2min system design (Figure 1A) offers certain significant advances over the other designs previously employed: fully-automated, software-selection of mixing times over six orders of magnitude (ms to minutes) with 1 ms time resolution; temperature control (0 - 25 C); quench temperature control (0 C); on-line connection for ‘bottom-up’ workflows; two-way communication for reciprocal control of chromatography; automated digestion column wash injection; intercalated blank injections and sample list scheduling of multiple runs.

**Figure 1:**
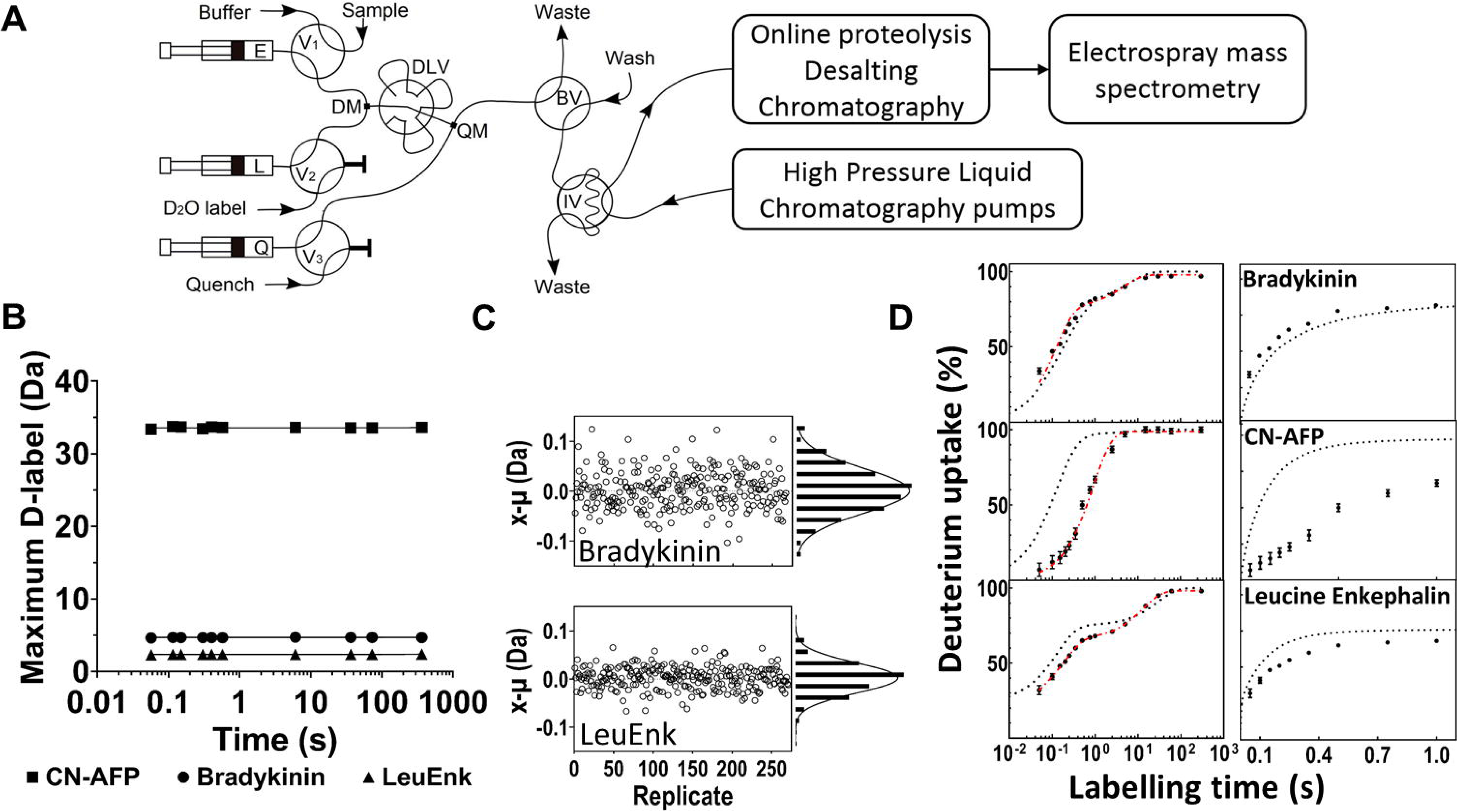
Characterisation of the fully automated ms2min system. (A) Instrument schematic. Three syringes (E, L and Q) store and deliver the buffers (protonated, deuterated and quenching buffers, respectively), with automated fill/delivery via valves V1-3. Unlabelled sample is introduced through V1 in the flow tube before the D_2_O mixer (DM). Labeling time is varied by matching fluid velocity and the choice of delay loop volume from six installed loops on the delay loop valve (DLV, four loops shown in the figure for clarity) prior to mixing with quench at the quench mixer (QM). During the labeling reaction the flow is directed to waste by a bypass valve (BV) to allow high flow-rates, in a subsequent step the quenched labeled sample is loaded onto the high pressure LC injection valve (IV) and introduced into the LC system enabling online LC-MS for ‘bottom-up’ workflows. (B) All labeling times tested exhibit identical back-exchange: the deuterium incorporation observed for fully-deuterated synthetic peptides is invariant with mixing time over more than five orders of magnitude. Linear regression fit with gradient <0.00012 in all cases. (C) Deviation from the mean measurement of deuterium label at 100 ms for Bradykinin and Leucine Enkephalin, n=270 replicate data points with dispersion of 3%. (D) Synthetic peptides exchange at different rates depending on their structure. Absolute hydrogen-exchange (expressed as % deuteration) shown for three peptides compared with theoretical maximum chemical exchange (black dotted curve), error bars denote ±1 r.s.d. from n=3. Stretched exponential fit to experimental data (Red dashed line).

### Precision of D-labeling

Measurement variability and labeling precision of the system were first determined for the quench-flow deuterium-labeling method. On three separate days we measured the D uptake of 50, 70 and 150 replicates of Bradykinin and Leucine enkephalin peptides at 100 ms mixing time. The short labeling period close to the limit of quantification was selected as these data points can be very sensitive to variability in the labeling reaction. Deviation from the mean value (*µ*) of each measurement and a histogram of the deviations are shown in Fig. 1C. Random, uniform and symmetrical distribution of the deviation from the mean was observed with <0.01 Da variability between days.The 95% confidence interval was ±0.01 Da (0.2% of the signal amplitude) for both Bradykinin and Leucine Enkephalin.

### Back exchange is independent of mixing time

The sample experiences different dwell times and flow velocities and characteristics, with respect to different mixing (D-labeling) times. This has the potential to confound attempts to correct for back-exchange, which is required to compare proteoforms and to calculate protection factors and free energy of stability values (ΔG_ex_). Therefore, we sought to determine whether there is back-exchange variation with mixing time. The highest measured dependence of back-exchange on mixing time is 0.012% from the linear regression of all five peptides, Fig. 1B. In all five peptides, with six different mixing loops that span labeling across four orders of magnitude, no significant correlation could be identified as indicated by the shallow gradient and poor linear correlation coefficient (R^2^ <=0.32). The back exchange shows a similar trend between the peptides analyzed by conventional automation, though the ms2min system preserves more of the deuterium label (Table S3). Therefore, we were able to correct for back exchange within an experiment by reference to a single fully-deuterated sample.

### Quantitation of peptide stability

Peptides often have weak protection factors against hydrogen-exchange (36, 37). This renders them intractable by conventional sampling (typically >10 s mixing) as much or even all labeling is complete within the dead-time of the experiment. We sought to determine the extent (i.e. lower limit) to which the ms2min system can quantify peptide stability in solution. We initially chose to measure the stability of synthetic peptides with the ms2min system as the simple workflow, small datasets and lyophilised samples allow us to validate our approach. Therefore, we collected data for a mixture of three peptides - a hormone (bradykinin), a neurotransmitter (leucine enkephalin), and a consensus sequence antifreeze protein (CN-AFP). The deuterium incorporation was measured at fifteen time points from 0.05 - 300 s and compared with their calculated maximum intrinsic exchange rates (i.e. rate corresponding to a fully unprotected species (36)). The data were corrected for the back-exchange of the system, permitting direct comparison between the measured and theoretical HDX rates. As the observed deuterium incorporation is the sum of a number (q - corresponding to the exchangeable amide groups in the peptide) of exponential exchange rates (*k*_*j*_), the ideal case would be to fit the data to q exponential phases. This becomes intractable for q>3, given the number of time points that would need to be sampled. Therefore, we fit the data to a stretched exponential function with the fewest phases required to generate a good fit and used this ensemble rate constant to calculate per peptide protection factors (Pf) and stability (G), as previously shown (38). In each case, it would be impossible to quantitatively estimate peptide stability under conventional labeling time regimes (>10 s). We were able to generate good fits to the HDX-MS data automatically and propose Pf for each peptide (Table 1). Interestingly, Bradykinin is observed to be natively disordered in solution, with no protection against maximum theoretical hydrogen exchange rates, Fig. 1D. All three peptides contained a single amide proton that exchanges even faster than the dead time of this experiment (50 ms), indicating that there is additional information to be gained from faster instrument performance in future.

Since the ms2min system is in-line with LC-MS for ‘bottom-up’ experiments, a similar quantitative assessment of peptide stability was then applied to study allosteric regulation of glycogen phosphorylase.

### Quantitative assessment of local stability in apo-GlyP

GlyP forms a dimer of a large (97 kDa) polypeptide chain and so it represents a major challenge to quantitatively link local changes in structure and stability in response to its many regulatory influences. Our approach, as established above, stands to provide this link, but only if high data density can be achieved, which results in high structural resolution and data confidence. For such a large dataset (three protein conditions each comprising hundreds of individual peptide fragments measured at ten deuterium labeling times, together with simulated uptake data), it is important that all data acquisition and nonlinear regression are automated. 273 unique peptides were identified, corresponding to 96.4% sequence coverage (Fig. 2A). In this way, a schedule of ms2min labeling experiments of GlyP - from 50 ms to five minutes - was acquired and analyzed automatically.

**Figure 2:**
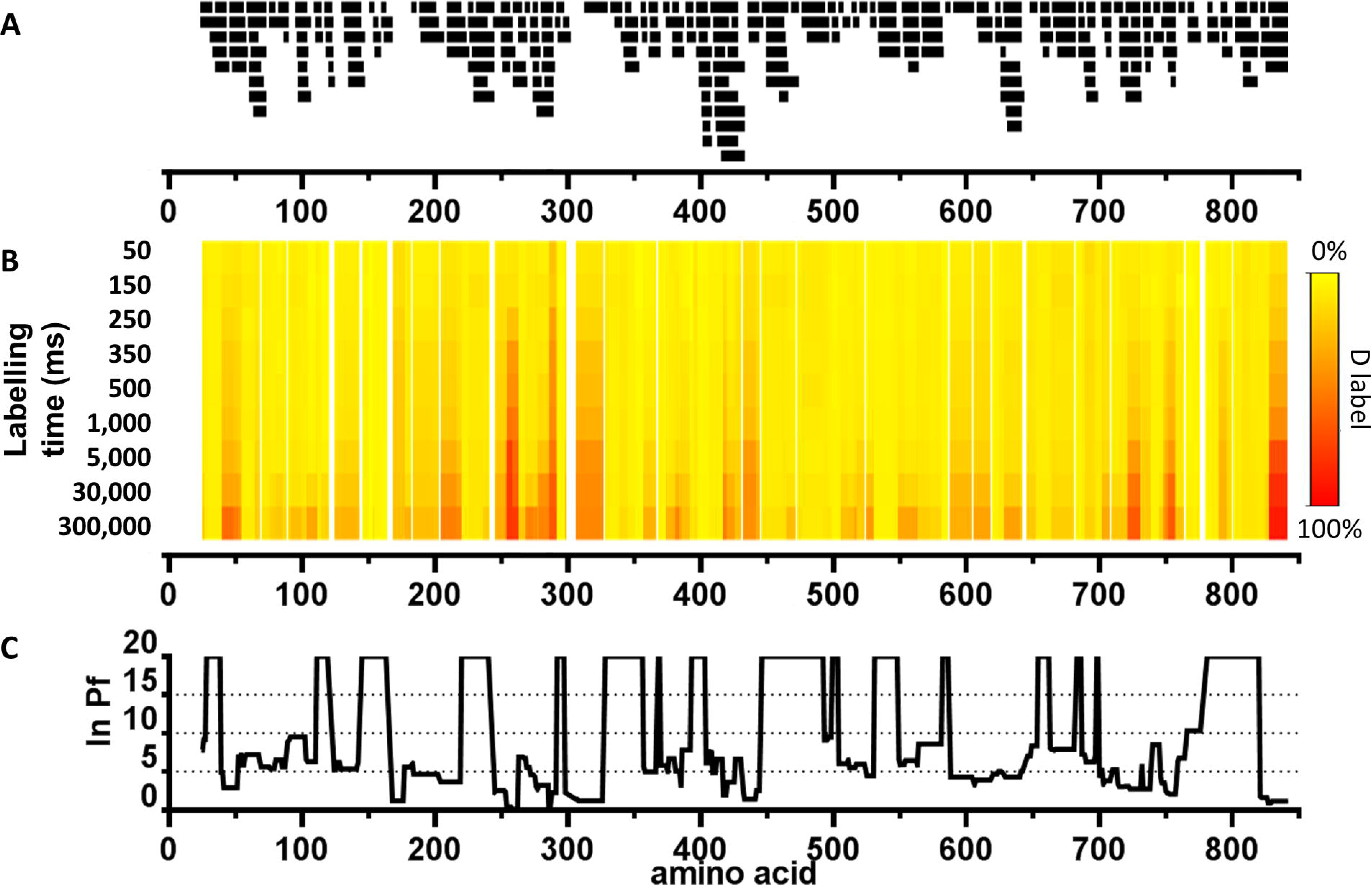
Solution phase local stability and structural perturbation in apo GlyPb dimer. (A) Coverage map of GlyP: Mass spectral assignment of GlyP peptides resulting from pepsin enzyme digest. 273 peptides were identified, corresponding to 95% protein coverage. (B) Heat map indicates the extent of the hydrogen-exchange at each labeling time (rows), given as relative fractional uptake of deuterium (%). Peptide level data has been averaged per amino acid with linear weighting. (C) Average hydrogen-exchange protection factors per amino acid. Up to two protection factors were determined for each peptide (black bars in panel A) and weighted by the fraction of the peptide exchanging under each regime. Gaps in the data were linearly interpolated. Upper limit of quantitation at ln(Pf) = 20.

We characterized the local stability of GlyP in weakly structured regions (i.e. not the folded core which has HDX half lives > hours) by measuring the structural dynamics in solution at physiologically relevant pH (7.0). Previously it was determined that the subunit interface of the dimer has two main contact regions on opposite sides of the enzyme (1): one between the cap region (residues 35-46) and the α1-α2 loop, β7 and the α2’ helix (47-78) of the opposite subunit. The other contact is antiparallel association of the two tower helixes, α7 and the immediately adjacent structural elements. The pattern of HDX protection factors matches well to the crystallographic dimer interface (Figure 2B-C) with no additional large protection factors that would indicate tetramer (Fig. S9,S13). The overall level of exchange in apo-GlyP T-state is low, consistent with a natively folded and rigid enzyme structure.

With a detailed map of local stability in the apo-state of GlyPb established, we then examined perturbations upon activation/inhibition. The entirely automatic approach to obtain protection factors and free energy of stability was applied to GlyP in two allosterically regulated states: active GlyPa phosphorylated at Ser14 (pSer14-GlyPa) in the R-state and inactive GlyPb bound to glucose-6-phosphate inhibitor (GlyP:G6P) in the T-state.

### Apo GlyP T-state closely resembles GlyP:G6P T-state

Binding of the allosteric inhibitor glucose-6-phosphate (G6P) to the nucleotide allosteric site resulted in only local changes in structure and stability, with some exceptions of long-range minor alterations to helices, α11 and α25 (Fig. S2-10). The G6P interface overlaps the AMP binding site (39), located within the nucleotide allosteric site at a subunit-subunit interface near the C-terminus (Fig. 3). Largest changes in stability were observed for the N-terminus of the α8 helix (−0.4 kcal/mol). Consequently, the C-terminus of α8 and the preceding 280s loop, destabilize by 0.1 kcal/mol. This may represent direct mediation of allostery from the nucleotide site via the rigid α8 helix to the active site, although it is unclear how the partial release of the 280s entropic gate can manifest as inhibitory. The footprint of bound G6P on the polypeptide chain is discernible from the HDX-MS data, yet it is considerably more diffuse than might be anticipated, based on the crystallographic contacts (Fig. S11). Notably, only one inter-chain (van der Waals) contact is identified from the co-crystal structure (between G6P sugar ring around O2 and Val40’ of the opposing subunit), yet there is extensive protection against hydrogen-exchange observed throughout the a1’-a2’ loop, stabilising those amino acids by −0.2 kcal/mol (Fig. 3). This indicates that the loop has considerable conformational flexibility in the apo-form that is significantly constrained by the contacts formed upon ligand binding, namely van der Waals contacts with G6P and with Ile68 and Trp69. Though perhaps expected, no amide hydrogen perturbation is observed in the beta sheet at the rear of the nucleotide site, or in α3, which both consist of extensive H-bonding networks.

**Figure 3:**
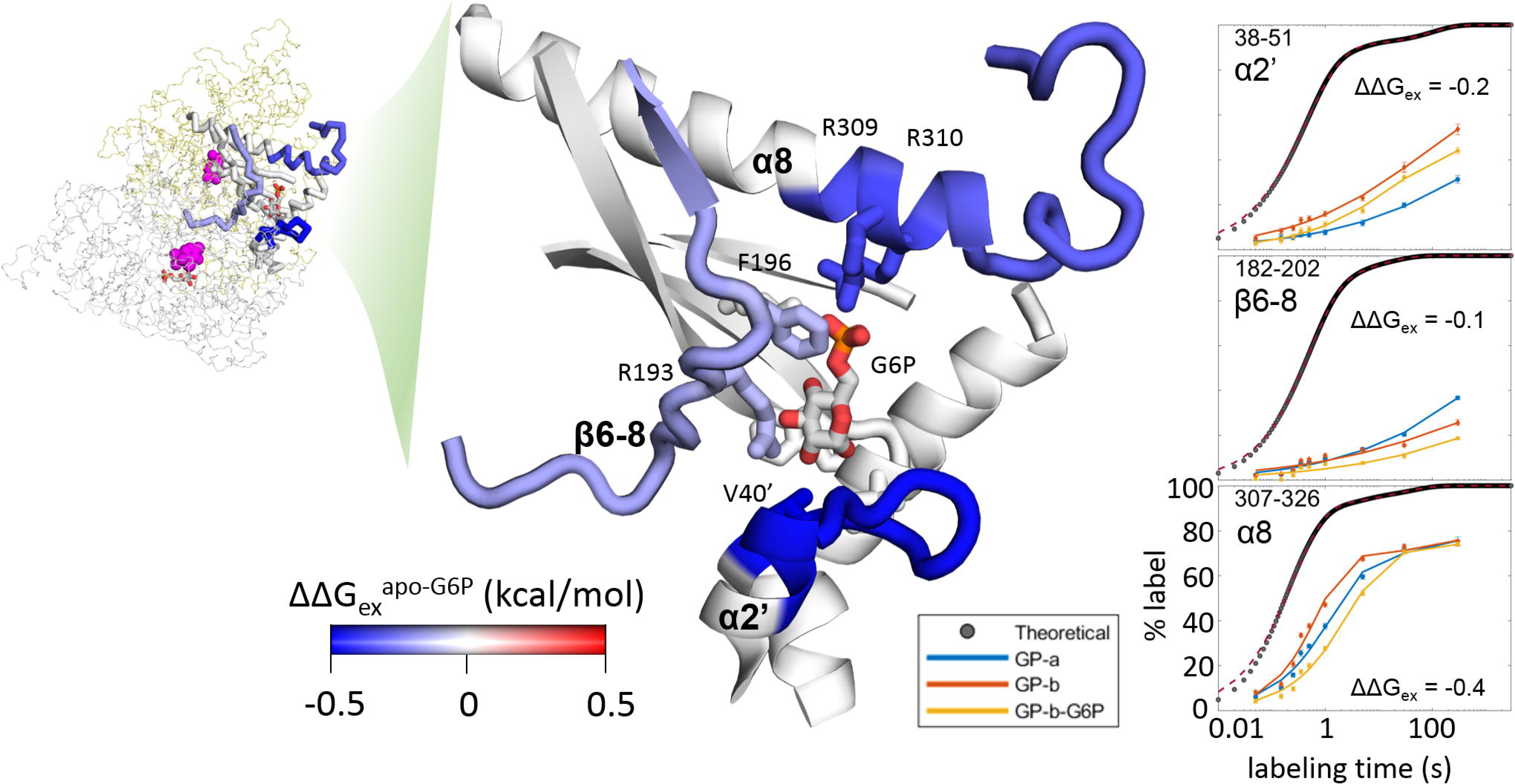
Direct local stabilization of GlyP by binding of glucose-6-phosphate (G6P) to the nucleotide site. The footprint on the polypeptide chain of bound G6P is discernible from the HDX-MS data, yet is more diffuse than might be anticipated. The α1’-α2’ loop is considerably stabilized by −0.2 kcal/mol, even though there is only one contact between G6P and the opposing monomer in the co-crystal structure (SI Fig. 11). All stability values in kcal/mol, estimated from HDX-MS rates (see SI). Plots: apo-GlyPb (red); pSer14-GlyPa (blue); GlyPb:G6P (yellow); theoretical maximum HDX rate for unstructured polypeptide (black traces).

### Allosteric activation alters local stability throughout the protein

Phosphorylation at the N-terminus (pSer14) induces the well-established flip in local conformation from T-state to R-state. The active and inactive states have notable structural differences, with the major changes known to occur in (i) the catalytic site, (ii) the nucleotide site and (iii) the tower helix (2, 10). We describe the solution phase stability changes in these regions below.

### Catalytic site

pSer14-GlyP (GlyPa) exhibits large destabilization extensively throughout the catalytic site (Fig. 4). Largest differences were observed in α6 (+0.3 kcal/mol), β13 (+0.3 kcal/mol) and the β22-α21 loop (+0.2 kcal/mol). Several of these affected amino acids make direct contacts with the pyridoxal phosphate (PLP) cofactor and may contribute to reorientation for productive catalysis. This allosteric effect is long range: 57 Å from the phosphorylated N-terminus to beyond the PLP. Several of the observed sites within these PLP-contacting loops are shielded from solvent in the crystal model, indicating that a considerable dynamic rearrangement of the PLP-polypeptide interactions must occur to result in hydrogen exchange.

**Figure 4:**
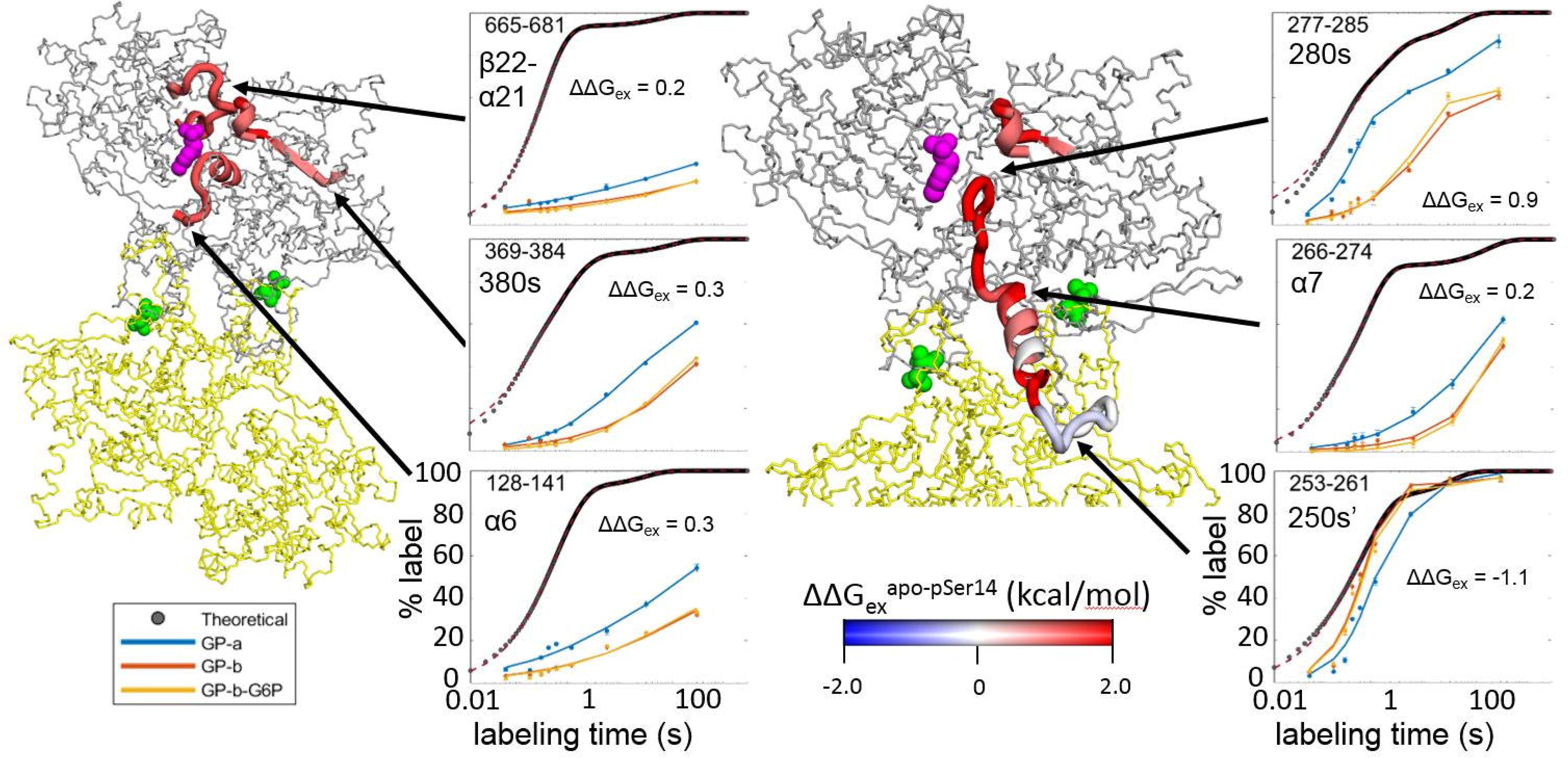
Phosphorylation at Ser14 induces the well-established switch in conformation from T- to R-state. The catalytic site is extensively remodelled (left panel) upon transition to the R-state by phosphorylation at Ser14 (green). Perturbations in HDX-MS measured rates were detected in several of the loops that contribute to catalytic activity and bind PLP (purple). The tower helix region (right panel) appears to behave as an entropic lever with activation inducing stabilization in the 250s’ loop (bottom), slight destabilization in the tower helix (center) and large destabilization in the 280s loop at the entrance to the catalytic site (top). All stability values in kcal/mol, estimated from HDX-MS rates (see SI). Plots: apo-GlyPb (red); pSer14-GlyPa (blue); GlyPb:G6P (yellow); theoretical maximum HDX rate for unstructured polypeptide (black traces).

### Tower helix

Upon activation, the tower helix α7 (residues 262-276) alters its angle upwards 10° relative to the long axis of the dimer – increasing solvent accessibility - and unfolds a partial turn into the 250s’ loop. This necessarily breaks certain stable contacts with the tower helix’ of the other subunit (40). Stability changes are observed in solution consistent with this: the tower helix destabilizes by ΔΔG_ex_ of 0.2 kcal/mol at the N-terminal end (Arg269-Leu271) up to 0.9 kcal/mol at the C-terminal end (Ala272-Ile275). The 250s’ loop preceding α7 is found to be natively disordered with a positive (unstable) ΔG_ex_ of +0.8 kcal/mol in the apo-form, but is significantly stabilized in GlyPa by 1.1 kcal/mol. The inverse relationship is seen at the other end of the tower helix, in the 280s loop. It was correctly assumed by Johnson and co-workers that “these residues…become mobile” upon transition to the phosphorylated R-state. The relatively stable loop in the apo T-state (−0.8 kcal/mol) is destabilized by 0.9 kcal/mol (Fig. S9-10) (41). This provides direct solution phase evidence for the entropic switch mechanism and the role of the 280s loop in gating access to the catalytic site. Moreover, it quantifies the relative changes that occur upon enzyme activation. The reciprocal relationship between the stability of the 280s and 250s’ loops supports an extension to the hypothesis whereby these loops, connected by the rigid yet mobile tower helix, may act together as an entropic lever (42).

### Allosteric sites

– The cap’ (residues 35-46) and α2’ (residues 47-78) interface showed smaller changes than elsewhere in the allosteric transition between GlyPb and GlyPa, however this interface is somewhat stabilized in the active conformation, by an average of −0.2 kcal/mol. The cap’ and α2 have been observed to come closer together (40) and, in line with that, we see the solution phase HDX-MS protection is increased in those regions Fig. S9-10.

## Discussion

Structural switching associated with allosteric regulation of enzymes is often predicated on alterations in local stability. This is epitomised in the archetypal allosteric enzyme, glycogen phosphorylase which has five known sites of allosteric regulation. Access to the GlyP active site is hypothesised to be gated by an order/disorder transition in the 280s loop, constituting an entropic switch. We sought to measure the changes in local structure and stability in GlyP upon allosteric regulation to test this hypothesis and to provide a quantitative basis for the perturbations in solution phase structure.

### An automated millisecond hydrogen/deuterium-exchange instrument

Hydrogen exchange rates, under the EX2 regime, can be considered a surrogate of free energy of stability (ΔG_ex_). However, reliable quantitation of exchange rates requires accurate measurement with deuterium labeling times spanning orders of magnitude – from milliseconds to minutes. We created a fully-automated, on-line quench flow instrument (“ms2min”) capable of making these measurements in GlyP with flexible labeling with short dead-times. Initial tests on synthetic peptide hormone (bradykinin), neurotransmitter (leucine enkephalin) and consensus anti-freeze protein (CN-AFP) validated the approach by permitting calculation of protection factors, even in natively unstructured regions. The new ms2min system developed here has significant merit from a technical standpoint: considerable potential for HDX-MS resolved at the millisecond level has been demonstrated previously, for example applied to the study of protein folding (11) and to TEM lactamase catalysis (15). The ms2min system stands to be particularly impactful in the future study of time-resolved (i.e. non-equilibrium) protein conformational dynamics, given its flexible and high temporal resolution, with software control over single millisecond differences in labeling time, and online full automation.

### Apo GlyPb in solution

Apo form GlyPb could be seen as a dimer in solution with HDX-MS protection patterns in keeping with the known crystallographic dimer interface. Few changes upon binding of inhibitor, glucose-6-phosphate (G6P) were observed outside of the nucleotide site, confirming that the apo GlyPb resides in a T-state. Just one predicted interaction between G6P and the α1’-α2’ loop, at Val40’, is sufficient to induce large stabilization in that part of the opposing monomer and gives rise to a diffuse footprint upon inhibitor binding. More direct impact on catalysis was observed as the stability of the 280s loop was reduced, likely mediated via the rigid body helix α8 that experiences the largest stabilizing effect by directly binding to G6P in the nucleotide site. The impact of this change in 280s stability is not immediately clear and warrants further study, such as comparison with the effects of AMP, but it is attractive to consider it may be a necessary conflation of the dual functions of the nucleotide site which binds to activating and inhibiting allosteric regulators.

### Remodelling of the active site in R-state transition

Activation of GlyP by phosphorylation at Ser14 results in large perturbations in the structural ensemble throughout the protein. Although the site of phosphorylation itself, at the N-terminus, was absent in the peptide coverage, the allosteric effects spanned almost 6 nm to the far side of the catalytic site. Multiple loops that contact the pyridoxal phosphate (PLP) cofactor exhibited increased hydrogen-exchange. Much of the solvent accessibility of these loops is via the deep catalytic site itself, which makes it particularly attractive to consider that these dynamic changes result in a reorganisation of PLP orientation in the pocket, with the implication that PLP is held in place somewhat loosely.

### Direct evidence for an entropic switch gating catalytic site access

In the tower helix region – known to traverse the largest geometric changes in the T/R-state transition – the 250s’ loop is found to be natively disordered in the T-state. Transition to the R-state reciprocally stabilizes the 250s’, together with the unfolding of the N-terminal turn of the adjacent tower helix (α7). The compensation in stability is even more pronounced at the far (C-terminal) end of the tower helix, where the 280s loop is highly stabilized. Importantly, the 280s loop is absent in some crystal models of the R-state. This has given rise to the hypothesis that it acts as an entropic switch to gate access to the active site: ordered/structured in T-state which blocks glycogen entry; disordered/unstructured in R-state which permits glycogen to access the deep catalytic pocket. Here, we quantitatively confirm this hypothesis by calculating the relative local changes in Gibbs free energy of stability from hydrogen/deuterium-exchange measurements of glycogen phosphorylase dimer in solution. Moreover, we observed data consistent with an extension to this hypothesis, proposing that the tower helix acts as an entropic lever to couple the order/disorder of the 280s loop with disorder/order in the 250s’ loop.

## Materials and Methods

### Materials

Glycogen phosphorylase b from rabbit muscle and Glycogen phosphorylase a from rabbit muscle were purchased from Sigma. Chemicals were purchased as follows: potassium phosphate dibasic (99.9%), potassium phosphate monobasic (99.9%), TRIS hydrochloride, Tris(2-carboxyethyl)phosphine hydrochloride (TCEP) and dimethyl sulfoxide-d6 (99.96%) from Sigma; dimethyl sulfoxide (Fisher Bioreagents). Deuterium oxide (99.9% D) was purchased from Goss Scientific. Water, acetonitrile and formic acid (99.5 %) Optima™ LC/MS Grade were from Fisher Scientific. Five peptides were used to prepare the peptide mixture, including Bradykinin (RPPGFSPFR) obtained from Sigma, Leucine enkephalin (YGGFL) from Waters and CN-AFP (DTASDAAAAAALTAANAAAAAEKTAADAAAAAAATAA) from Peptide Synthetics. **ms2min system design:** The ms2min system was designed to allow fully automated HDX labeling from the low millisecond to minutes time-scale with software control of labeling time, temperature control of labeling and quenching and on-line connection to a 2D-chromatography system for conventional ‘bottom-up’ workflows. **Deuterium-labeling of peptides:** All H/D experiments were performed in triplicate at 23°C. The fast mixing system was directly connected to the digestion/separation chamber, however the pepsin column was replaced with a narrow-bore Ti union when the peptides (not protein) were analyzed. Peptides were solubilized in 20 mM potassium phosphate buffer, pH 7.40 in H_2_O and labelled by rapid 20-fold dilution with 20 mM potassium phosphate, pD 7.46 in D_2_O. This mixture passed through six different loops at different velocities, depending on the labeling time (0.05, 0.1, 0.15, 0.20, 0.25, 0.35, 0.5, 0.75, 1, 2.5, 5, 15, 30, 60 and 300 s). The labeling reaction was then rapidly quenched by 2-fold dilution with quench buffer (100 mM potassium phosphate, pH 2.50 in H_2_O) at 0°C. **Deuterium-labeling of GlyP:** Protein samples were analyzed in triplicate in a completely randomized manner at 9 time points (0.05, 0.15, 0.25, 0.35, 0.5, 1, 5, 30, and 300 s), essentially as described above. Digestion was performed online with a pepsin column (Waters). Protein samples were prepared in 40 mM TRIS hydrochloride, 1 mM TCEP, pH 7.00 in H_2_O and labelled by 20-fold dilution in 40 mM TRIS hydrochloride and 1 mM TCEP at pD 7.00 in D_2_0. The labeling reaction was then rapidly quenched by 2-fold dilution with quench buffer (100 mM potassium phosphate, pH 2.50 in H_2_O) at 0 °C. **Kinetic analysis:** Mass spectrometry data were corrected for back exchange with measurements from lyophilized sample resuspended in deuterated quench buffer and equilibrated overnight at 37 °C. Measurement of labeling was essentially as previously described (43). An in-house program was developed (MatLab; Mathworks) for automatic fitting to stretched exponential kinetics and for calculation of protection factor (Pf) in experimental data versus published values for unstructured amino acids. This involves four steps: (i) simulation of intrinsic uptake curves (ii) F-test to select appropriate number of components for fitting of intrinsic exchange simulation to a stretched exponential model, (iii) fitting experimental data to the chosen model and (iv) plotting.

## Acknowledgements

This work was supported by InnovateUK grant 102612. Applied Photophysics Ltd holds patents relating to design of the ‘ms2min’ instrument.

## Notes

#### Summary of Updates

Reinterpretation of results, edited for clarity.

