## Supplementary Information for "Allosteric regulation of glycogen phosphorylase solution phase structural dynamics at high spatial resolution"

### Supporting information

### SI Materials and Methods

**Materials.** Chemicals were purchased as follows: potassium phosphate dibasic (99.9%), potassium phosphate monobasic (99.9%), TRIS hydrochloride, Tris(2-carboxyethyl)phosphine hydrochloride (TCEP) and dimethyl sulfoxide-d6 (99.96%) from Sigma; dimethyl sulfoxide (Fisher Bioreagents). Deuterium oxide (99.9% D) was purchased from Goss Scientific. Water, acetonitrile and formic acid (99.5 %) Optima™ LC/MS Grade were from Fisher Scientific. All other ultrapure water used was purified on a Milli-Q Advantage A10 system (Merck).

Five peptides were used to prepare the peptide mixture, including Bradykinin (RPPGFSPFR) obtained from Sigma, Leucine enkephalin (YGGFL) from Waters, CN-AFP (DTASDAAAAAALTAANAAAAAEKTAADAAAAAAATAA) from Peptide Synthetics and cTPRH1 (AEAWYNLGNAYYK) and cTPRS (AEAKQNLGNAKQK) were synthesized in-house on a Biotage parallel synthesizer (SYRO II) on wang resin with purification by reverse phase against a C8 column (Polaris). All of the peptides were dissolved in DMSO, or d6-DMSO for determination of the maximum deuteration level and back exchange of the system. The peptide mixture was prepared with each peptide at ≈5 µM concentration in 20 mM Potassium phosphate buffer at pH 7.40.

Glycogen phosphorylase a and b from rabbit muscle were purchased from Sigma. The protein was dissolved and diluted in 40 mM TRIS hydrochloride and 1 mM TCEP at pH 7.00 to a final concentration of 10 µM. For the glycogen phosphorylase b and inhibitor (glucose-6 phosphate) equilibrium experiments same buffers and concentrations were used, with glycogen phosphorylase b to glucose-6-phosphate ratio of 1:200.

**msHDX system design.** The msHDX system was designed to allow fully automated HDX labelling from the low millisecond to hours time-scale, with high temporal resolution, temperature control and on-line connection to a 2D-chromatography system for conventional 'bottom-up' workflows. A schematic representation of the system is shown in Figure 1.

**Calibration of ms2min labelling times.** Time point accuracy is determined by relating the applied velocities of the labelling and sample syringes (stepper motors) with the accurate calibration of the volume of each of six delay loops between the labelling $D_2O$ mixer (DM) and quench mixer (QM). Each syringe stepper-motor step to volume was determined gravimetrically by measuring the volume of $H_2O$ flushed through the system, with flow-rate accuracy determined to be within 1% for all syringes. The delay loops were then calibrated using 24 mg/mL nicotinic acid (Sigma Aldrich) in 0.1 M HCl loaded into the sample syringe; the quench and label syringes were loaded with 0.1 M HCl. 1 mL of nicotinic acid was delivered from the sample syringe and 1 mL loading the delay loop. Next 1 mL of HCl was delivered from the quench syringe, flushing the final tubing section. Finally 1 mL of HCl from the label syringe was collected and the loop volume calculated from the equation below:

$$Volume\ of\ loop\ (uL) = \frac{A261nm\ collected\ sample}{A261nm\ undiluted\ standard} \times\ Delivered\ volume\ (uL) \qquad \text{Equation S1}$$

Where delivered volume = $1000\ uL$.

Timing accuracy was determined using the base catalyzed hydrolysis of 2,4-dinitrophenyl acetate (DNPA , Acros Organics) to 2,4-dinitrophenol (DNP) [2]. A solution of 2 mM DNPA in 2% (v/v) DMSO, 0.1 M HCl (sample syringe) was mixed 1:10 with 0.5 M NaOH (labeling syringe), and quenched 1:1 with 1 M HCl (quench syringe). DNPA and DNP in the acid quenched reaction solutions were separated on a C18 reverse phase column (Telos) with isocratic 50% acetonitrile/water, 0.1% trifluoroacetic acid, and detected by UV absorbance (254 nm). Timepoints between 40 ms and 500 ms were collected in triplicate, as well as 0 ms time points unreacted solution where acid was added by hand before the NaOH was added. For direct comparison, the same reactions solutions were analysed using UV absorbance at 360 nm (1 ms to 500 ms) in a SX20 stopped-flow spectrophotometer (Applied Photophysics Ltd).

Both ms2min and stopped-flow data sets were fitted with a single exponential decay model using the fitting method of Kemmer and Keller [3].

**Deuterium-labelling with the msHDX system.** All H/D experiments were performed at 23°C labeling temperature in triplicate. The fast mixing system was directly connected to the digestion/separation chamber, however the pepsin column was replaced with a narrow-bore union when the peptides were analyzed. Labeling was initiated by injecting 20 µL of the peptide mixture or protein into the system by a titrator.

During the peptide mixture experiments the carrier syringe X contained 20 mM potassium phosphate buffer, pH 7.40 in $H_2O$. The labelling buffer, 20 mM potassium phosphate buffer at pH 7.06 in $D_2O$, was placed in the second syringe Y. The mixing ratio in the first mixer was 1:20. Depending on the labeling time, this mixture passed through six different loops at different velocities. The labeling reaction was then rapidly quenched by delivering the quench buffer, 100 mM Potassium phosphate buffer at pH 2.50 in $H_2O$ at 0°C degrees into the second mixer, which constituted an approximate 2-fold dilution. The peptide mixture was quenched at 15 time points: 0.05, 0.1, 0.15, 0.20, 0.25, 0.35, 0.5, 0.75, 1, 2.5, 5, 15, 30, 60 and 300 s. The quench buffer was kept in the quench chamber at 0°C degrees in the third syringe Z of the system.

Protein samples were analyzed in triplicates in a completely randomized manner at 9 time points including 0.05, 0.15, 0.25, 0.35, 0.5, 1, 5, 30, and 300 s, with the msHDX instrument as described above. Digestion was performed online with a pepsin column. The buffers used while analyzing the protein samples were 40 mM TRIS hydrochloride and 1 mM TCEP at pH 7.00 in $H_2O$, 40 mM TRIS hydrochloride and 1 mM TCEP at pH 6.60 in $D_20$ and 100 mM potassium phosphate buffer at pH 2.50 in $H_2O$, as carrier, labeling and quenching buffers respectively.

**Back-exchange correction.** Maximum deuterated reference samples (peptide mixture) were analyzed separately, however both syringes X and Y contained 20 mM Potassium phosphate buffer at pH 6.60 in $D_2O$. The peptides were dissolved in d6-DMSO, and all subsequent dilutions prepared in the D2O buffer described above. The same procedure as for the labeling experiments was followed, and the labeling was quenched at the same time points as the labeled samples.

For back-exchange correction of the protein samples, glycogen phosphorylase b was predigested offline in a fully deuterated control buffer 20 mM potassium phosphate buffer at pH 2.55 in $D_20$ with pepsin, for 5, 10 and 30 min at RT. The digested fully deuterated peptides were then manually injected, processed and analyzed using the HDX-MS workflow described above, with the only exception that a narrow-bore union was placed instead of the pepsin column in order to avoid double digestion.

**LC-MS.** Online digestion, desalting and separation of the peptides was done on a Waters HDX Manager with an immobilized pepsin column (Enzymate BEH Pepsin Column 2.1 × 30 mm, 5 µm), C18 trapping column (VanGuard ACQUITY BEH 1.7-µm, 2.1 × 5 mm; Waters), and analytical C18 column (1.7-µm, 1.0 × 100 mm ACUITY BEH; Waters). Mobile phases were 0.1% formic acid in H2O (A) and 0.1% formic acid in ACN (B), such that their pH was 2.50. Peptides were trapped for 4 min at a flow rate of 100 µL/min. Approximately 2.5 pmol of each peptide, or 10 pmol of the protein, were delivered from the fast mixing system. The peptides were then loaded onto the analytical column and eluted using a linear gradient (15-40% over 4 min at a flow rate of 40 µL/min). During the protein analysis the separation of the digested peptides was with a linear gradient (5-40% over 7 min at a flow rate of 40 µL/min).

Eluates after separation, were directed into a quadrupole time of flight mass analyzer (Synapt G2-Si HDMS QTOF, Waters) with positive ion electrospray ionization tuned for collision induced dissociation (CID) and lock-mass correction (using Leucine enkephalin peptide, 556.2771 m/z). Mass spectra were obtained in Waters $HDMS^E$ mode (from 50 *m/z* to 2000 *m/z)* for 3D (LC, IM, m/z) peptide separation. The instrument configuration was the following: capillary voltage was 3.0 kV; cone voltage at 50 V; trap collision energy of 4 V, travelling wave ion mobility separation was done with 575 m/s, 36.5 V wave amplitude and 2.75 mbar $N_2$. Transfer collision energy 4 V was used for low energy scans and four separate ramps between 15-55V for high energy scans.

**Data analysis.** The identity of each peptide was assigned from $HDMS^E$ fragment data with ProteinLynx Global Server 3.02 (PLGS) (Waters). Deuterium incorporation was determined with DynamX 3.0™ (Waters), all assigned ion spectra were manually reviewed. The observed uptake was corrected for the observed peptide specific back exchange, to attain the absolute uptake for each detected peptide. Afterwards, the absolute uptake was normalized to the theoretical maximum

exchangeable backbone amides (excluding N-terminus and proline) and accounting for approximately 95% $D_2O$ for each labelling reaction. The deuterium uptake curves were generated by plotting the absolute deuterium uptake (%) against the labeling time.
Subsequent data filtering, fitting, normalization and protection factors calculation was done using in-house programs in MatLab (MathWorks). Structures were modelled in PyMol (Schrödinger). Statistical tests were performed with Prism v5.0 (Graphpad).

**Kinetic analysis.** An in-house Matlab code was developed for automatic calculation of the segment averaged protection factors (Pf) as a measure of the reduced exchange brought by the structure of the protein. The code involves three steps: generating intrinsic uptake curves, fitting them into one- or two stretched exponentials, and plotting and fitting the experimental uptake curves in the same manner. Firstly, the intrinsic chemical amide exchange rates were calculated and simulated for each peptide according to equation 1 as demonstrated previously by Bai et all and adapted from the excel sheet provided by Englander lab (available online here http://hx2.med.upenn.edu/).

$$D(t) = \sum_{i=2}^{n} (1 - e^{-k_{int}^{(i)} t})$$

*Equation S2*

Where n is the number of residues in each peptide, $k_{int}$ is the intrinsic rate constant of chemical exchange for each residue and t is the labeling time. This equation uses the sum of exponentials for each amide in a peptide to provide the degree of deuterium incorporation as a function of a labeling time. At the N-terminus of the peptide the first amide becomes a primary amine after proteolysis, thus the first residue back-exchanges quickly during the LC-MS analysis. As a result, index n starts from the second residue onwards. Proline does not contain an amide hydrogen, and so its rate constant will always be zero. We adapted all of the calculations from the excel spreadsheet available on W. Englander's website. From these we generated and simulated the uptake curves for each peptide, at the experimental pH and temperature.
The exchange kinetics were quantified as it was described previously, where the theoretical and experimental deuterium uptake were fit to a single- or double-stretched exponential function.

$$D(t) = Q[1 - e^{-((k)t)^{\beta}}]$$

*Equation S3*

$$D(t) = Q_1 \left[1 - e^{-((k_1)t)^{\beta}}\right] + Q_2 \left[1 - e^{-((k_2)t)^{\beta}}\right]$$

*Equation S4*

Where *D(t)* is the deuterium uptake as a function of the labelling time t, Q is the number of exchangeable amides, k represents the segment-averaged exchange constant, and β is an exponential stretching factor that accounts for the distribution of the exchange rates of the individual amides. The stretched exponential function is used when fitting the H/D kinetics, as it requires less adjustable parameters than the commonly used multi-exponential one. To confirm the suitability and aptness of the stretched exponential models, we performed an F-test on the one- and two-stretched exponential models. If the more complicated (two-stretched exponential) model is correct, then the relative increase in the sum-of-squares would be greater than the relative degrees of freedom ($F > 1$, $p < .0005$). Thus, only when it is statistically necessary the two-stretched exponential model will be selected.
The experimental uptake curves were then fitted in the same manner as the intrinsic curves, where the starting points for the amplitudes Q1 and Q2 were restricted to be half of the previously determined Q1 and Q2 from the intrinsic fits. Also, it was necessary to add arbitrary maximum uptake data at extremely long time points to aid the fits. This facilitated the correct fit particularly for the protected regions of the protein that are exchanging predominantly slow.
The segment averaged protection factors can be then estimated by using the ratio of the intrinsic

exchange rate constant ($k_{int}$) to the measured (experimental) rate constant ($k_{exp}$).

$$Pf = \frac{k_{int}}{k_{exp}}$$

*Equation S5*

Where Pf = protection factor against hydrogen exchange, $k_{int}$ = intrinsic amide hydrogen exchange rate constant from published values [1], $k_{exp}$ = fitted exchange rate constant for back-exchange corrected experimental data using Equation S3 or Equation S4.

Gibbs free energy estimates from HDX-MS data were calculated from the established relationship:

$$\Delta G_{ex}(HDX) = -RT \ln \frac{1}{Pf}$$

*Equation S6*

Where R = gas constant (K/kcal/mol), T = temperature (K) and Pf = protection factor from Equation S5.

### SI Results

*Table S1 Parameters obtained from manual fitting of theoretical and experimental data acquired for Bradykinin, CN-AFP and Leucine Enkephalin.*

| Peptide | Measured | | | | | | | Intrinsic | | | | | | |
|---|---|---|---|---|---|---|---|---|---|---|---|---|---|---|
| | $N_1$ | $N_2$ | $N_3$ | $k_1$ ($s^{-1}$) | $k_2$ ($s^{-1}$) | $k_3$ ($s^{-1}$) | β | $N_1$ | $N_2$ | $N_3$ | $k_1$ ($s^{-1}$) | $k_2$ ($s^{-1}$) | $k_3$ ($s^{-1}$) | β |
| Bradykinin | 1.1 | 3.8 | | 0.2 | 8.4 | | 0.7 | 1.2 | 3.8 | | 0.2 | 5.5 | | 0.8 |
| CN-AFP | 35.5 | | | 1.1 | | | 0.9 | 35.6 | | | 7.2 | | | 0.8 |
| Leucine Enkephalin | 1.0 | 1.0 | 1.9 | 50.4 | 0.0 | 3.2 | 1 | 1.0 | 1.0 | 1.9 | 0.1 | 302 | 7.6 | 0.9 |

*Table S2 Calculated segment averaged protection factors.*

| Peptide | ln(*Pf*) | | |
|---|---|---|---|
| | Pf1 | Pf2 | Pf3 |
| Bradykinin | -0.14 | -0.42 | |
| CN-AFP | 1.83 | | |
| Leucine Enkephalin | 1.79 | 0.86 | -0.15 |

*Table S3 Comparison of the back exchange observed when using the CTC-PAL automation and the ms2min system. The back exchange shows a similar trend between the peptides analyzed by CTC-PAL automation, though the ms2min system preserves more of the deuterium label.*

| | Maximum number of deuterons | Back-exchange CTC-PAL (LEAP) | Back-exchange ms2min |
|---|---|---|---|
| Bradykinin | 5 | 15% | 6% |
| CN-AFP | 36 | 13% | 7% |
| LeuEnk | 4 | 49% | 41% |
| cTPRH1 | 12 | 14% | 15% |
| cTPRS | 12 | 29% | 26% |

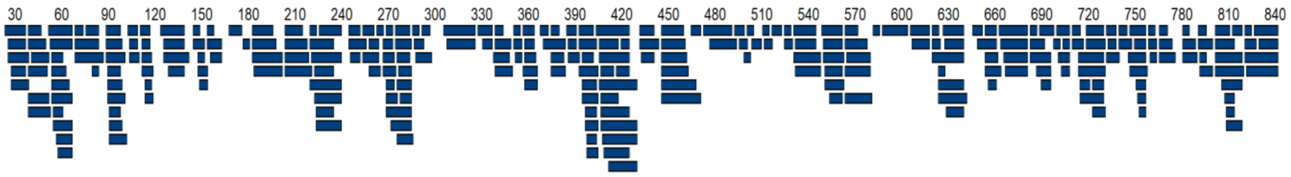


*Figure S 1. Sequence coverage of glycogen phosphorylase from rabbit muscle. The peptides obtained via peptic digestion and LC-IMS/MS analysis are shown as blue bars along the sequence numbers. Each bar under the sequence number annotation indicates an identified peptic peptide that was monitored during all HDX-MS experiments. These 273 peptides cover up to 800 amino acids of the total amino acid residues in the proteins, yielding a linear sequence coverage of up to 94.9%, with 4.04 redundancy.*

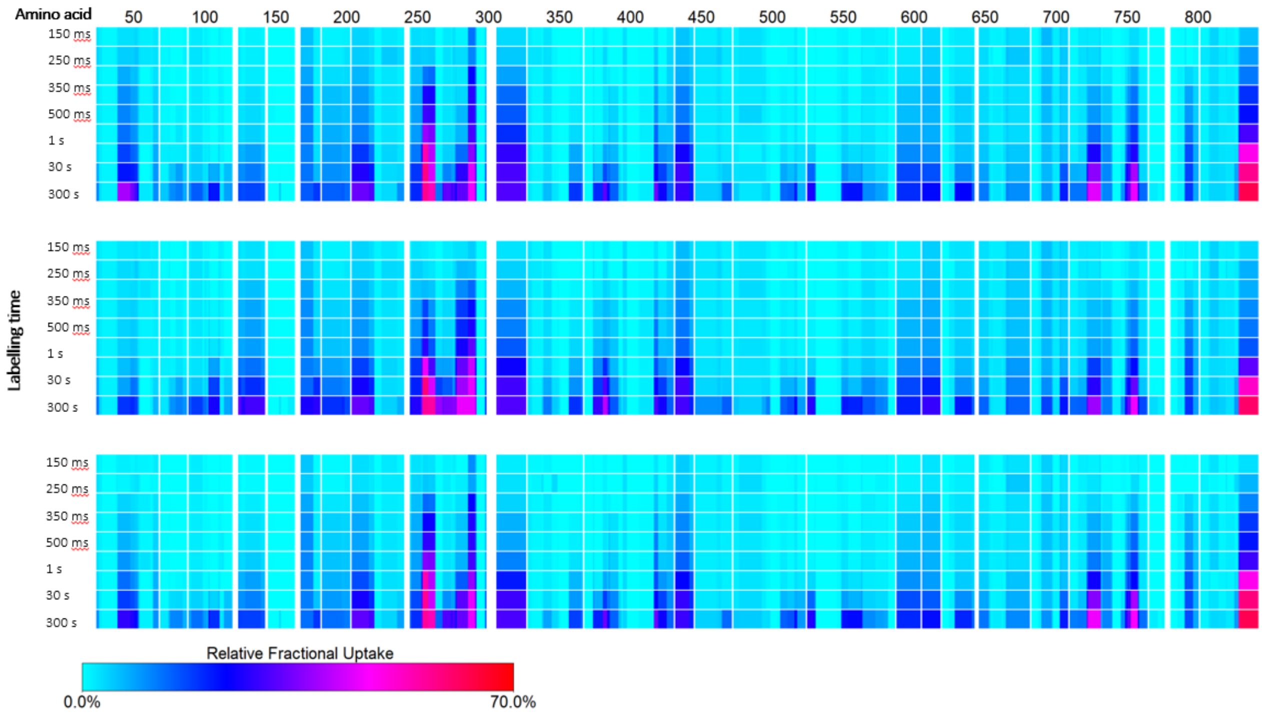


*Figure S 2. Heat maps of differential deuterium uptake for GlyP in active R-state (top), apo T-state (middle) and GlyP:G-6-P complex in T-state (bottom). No back-exchange correction and statistical filtering were performed for the heat map generation, made with DynamX (Waters). The HDX uptake is resolved at amino acid level, at nine time points including 0.05, 0.15, 0.25, 0.35, 0.5, 1, 5, 30, and 300 s from top to bottom. The deuteration level at each time point for each amino acid is color coded shown on left.*

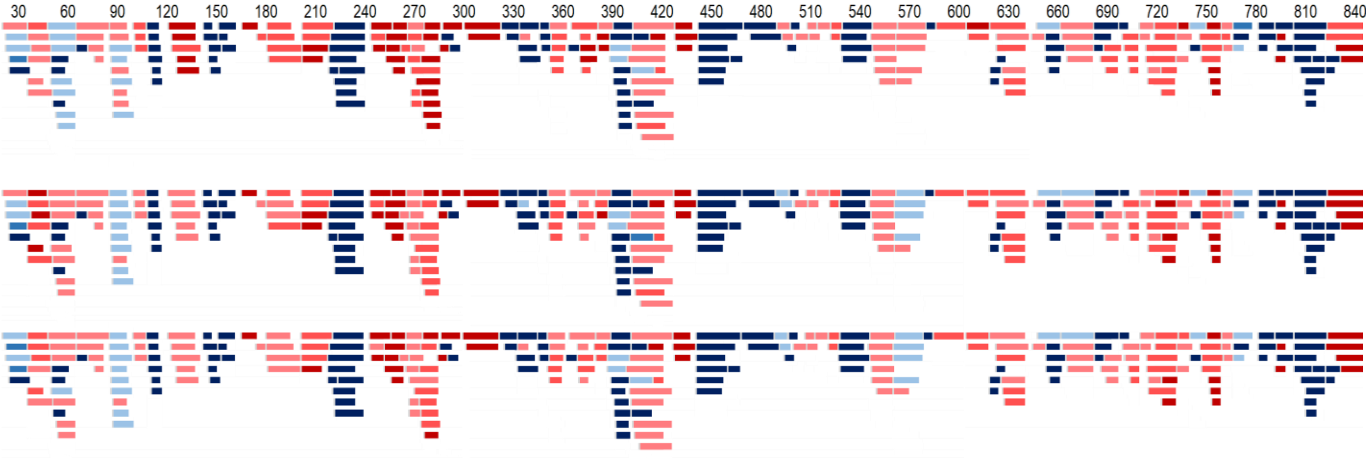


*Figure S 3. Protection coverage maps of plotted log (Pf) for GlyP in active R-state (top), apo T-state (middle) and GlyP:G-6-P complex in T-state (bottom). Back-exchange correction and statistical filtering were performed for the coverage map generation, made with in-house developed Matlab code for automatic calculation of protection factors.*

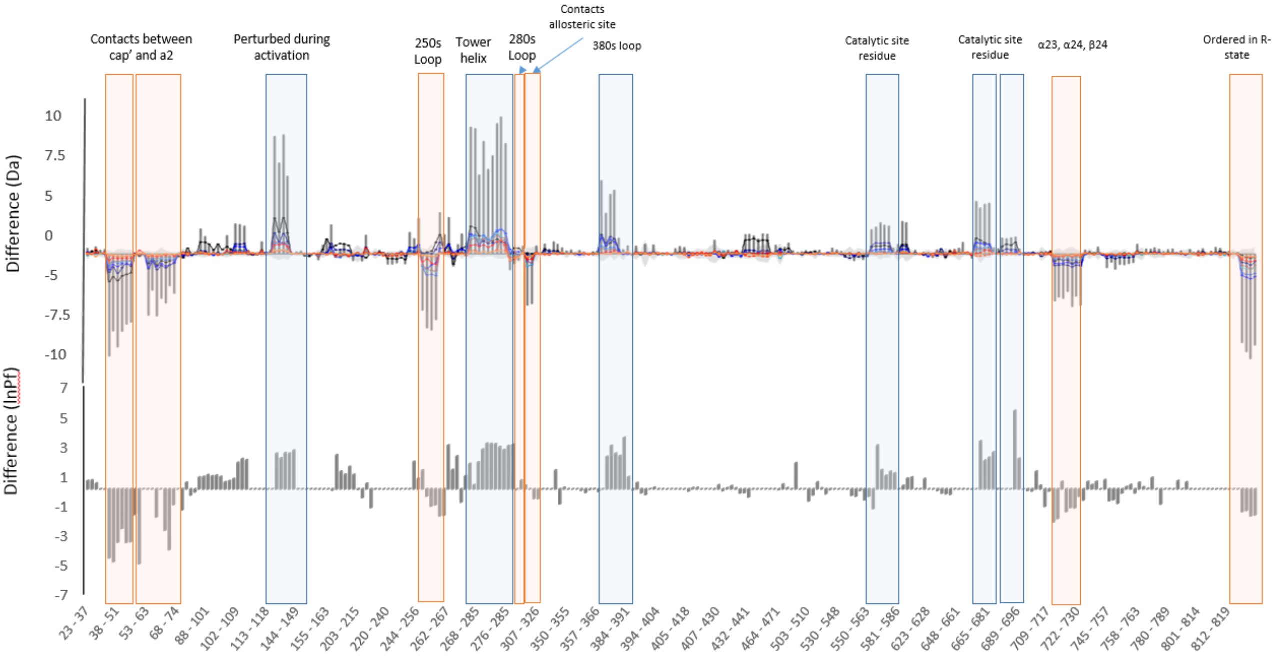


*Figure S 4. State difference maps. Difference of the observed deuterium uptake (top) and difference of the calculated ln(Pf) (bottom) among GlyP in active R-state and apo T-state. The deuterium uptake data from the apo T-state sample was subtracted from the data for GlyP in active R-state sample in order to create the deuterium uptake difference plot. Relative protection leads to a more negative value (bar on bottom side); deprotection (e.g. from an exposed domain interface) results in a more positive value (bar on upper side). The calculated protection factor data of GlyP in active R-state was subtracted from the data for apo T-state sample in order to create the protection factor difference plot. Relative protection leads to a more negative value (bar on bottom side); deprotection (e.g. from an exposed domain interface) results in a more positive value (bar onupper side ). Each horizontal bar represents a single peptide.*

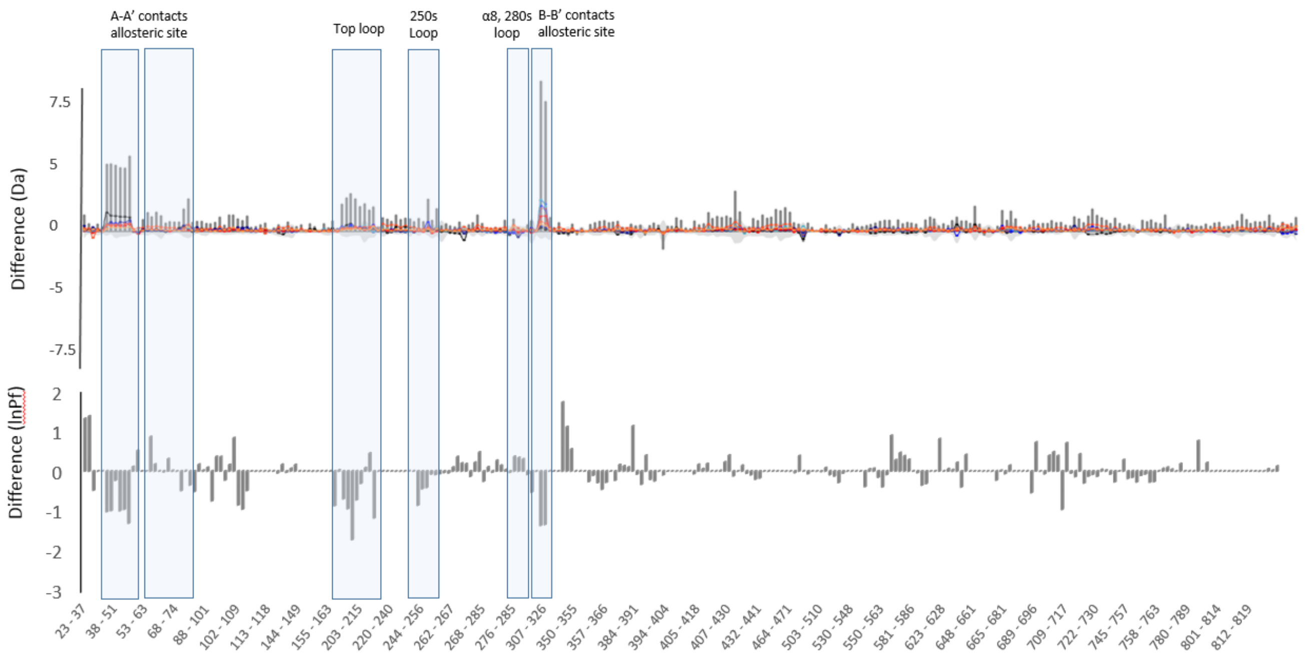


*Figure S 5. State difference maps. Difference of the observed deuterium uptake (top) and difference of the calculated ln(Pf) (bottom) among apo T-state and GlyP:G-6-P complex in T-state. The deuterium uptake data from the apo T-state sample was subtracted from the data for GlyP:G-6-P complex in T-state sample in order to create the deuterium uptake difference plot. Relative protection leads to a more positive value (bar on upper side); deprotection (e.g. from an exposed domain interface) results in a more negative value (bar on bottom side). The calculated protection factor data of GlyP:G-6-P complex in T-state sample was subtracted from the data for apo T-state in order to create the protection factor difference plot. Relative protection leads to a more positive value (bar on upper side); deprotection (e.g. from an exposed domain interface) results in a more negative value (bar on bottom side). Each horizontal bar represents a single peptide. Vertical scale is non-linear, but approximate values are shown from start residue (left) to end (right).*

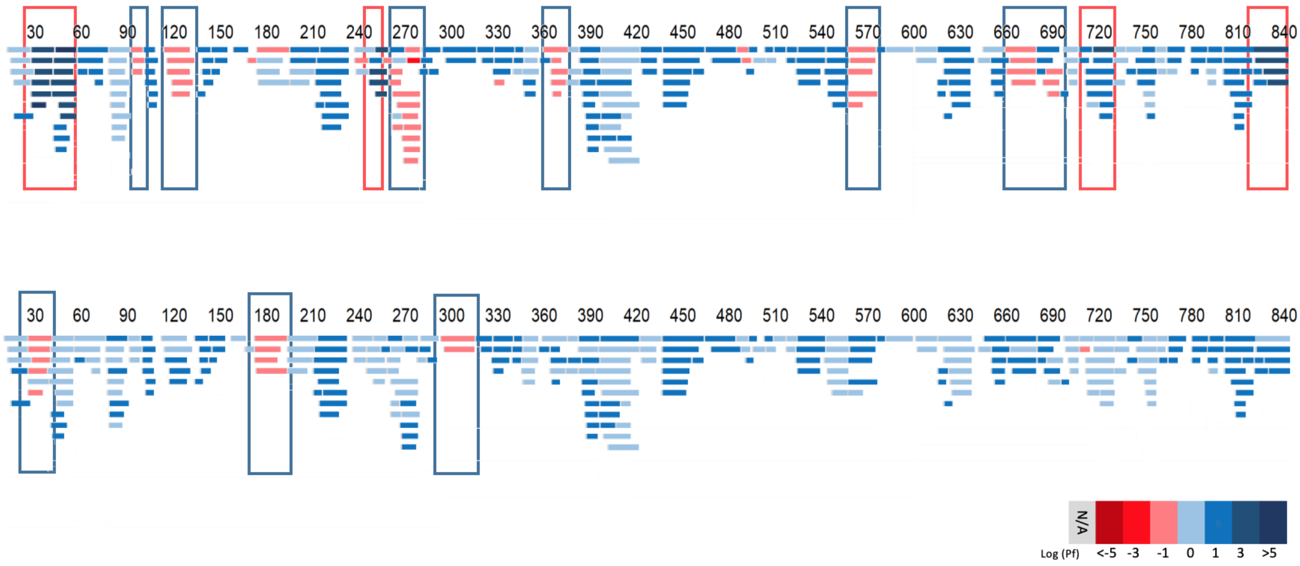


*Figure S 6. Difference in protection factors between GlyP in active R-state and apo T-state (top) and between inactive inhibitor-bound state and apo T-state (bottom). Each bar under the sequence number represents a peptide fragment monitored during the labeling experiments. Peptides colored according to the measured protection factors, where more positive value indicates more protection against hydrogen-exchange in apo GlyP. Regions of coherent difference highlighted in boxes.*

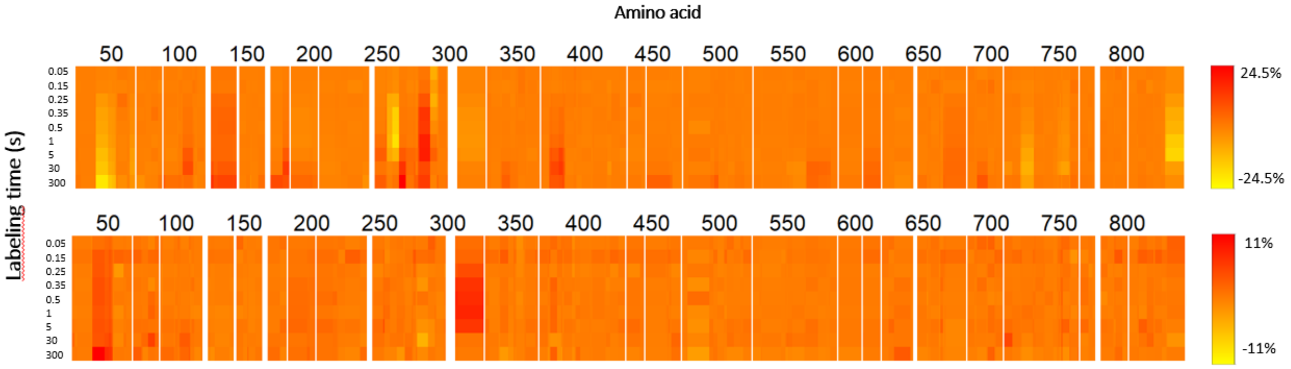


*Figure S 7. Heat maps of the difference in HDX labeling between different states: active pSer14-GlyPa minus apo (top) and GlyP:G6P minus apo (bottom). Several changes in the structural dynamics are evident only during a subset of the HDX time-course: for example the unstructured loop at the tip of the tower helix shows observable differences between 250 ms – 30 s deuterium labelling and only in active GlyP. The HDX uptake is resolved at amino acid level by linearly averaging data for overlapping peptides and is shown at nine D-labelling time points (0.05, 0.15, 0.25, 0.35, 0.5, 1, 5, 30, and 300 s). The relative deuteration level at each time point at each amino acid is color coded shown on the scale below; note different normalisation.*

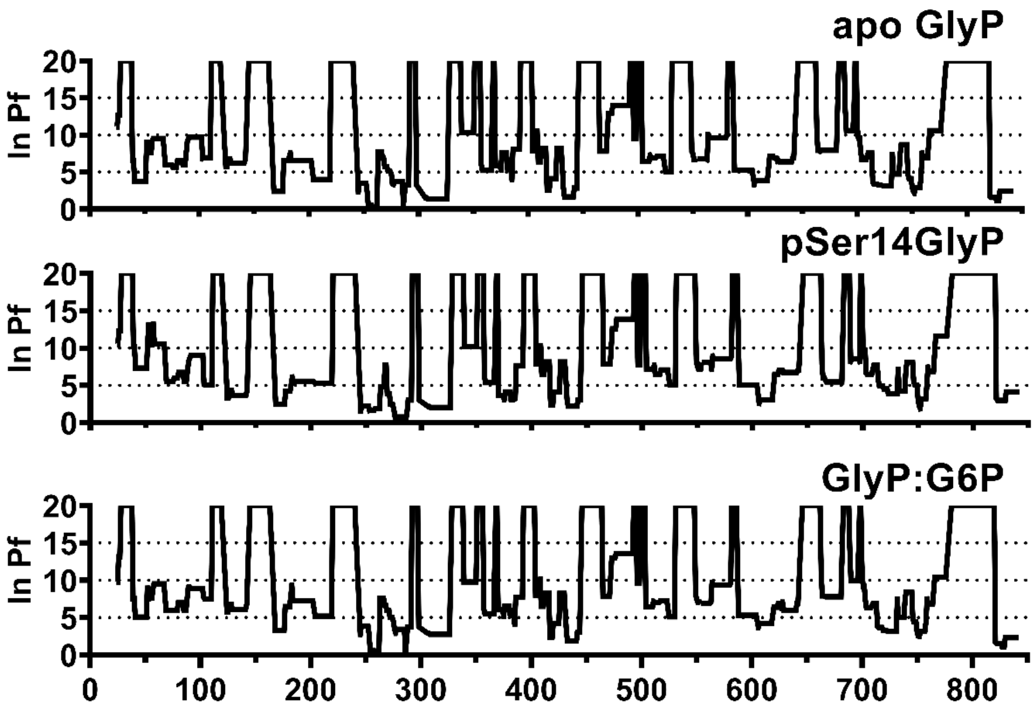


*Figure S 8. Protection factors per amino acid for GlyP in three states: apo GlyPb (top), pSer14 GlyPa (middle) and GlyP:G6P inactive complex (bottom). Values calculated from fitted exchange rate constants using Equation S5. (1)*

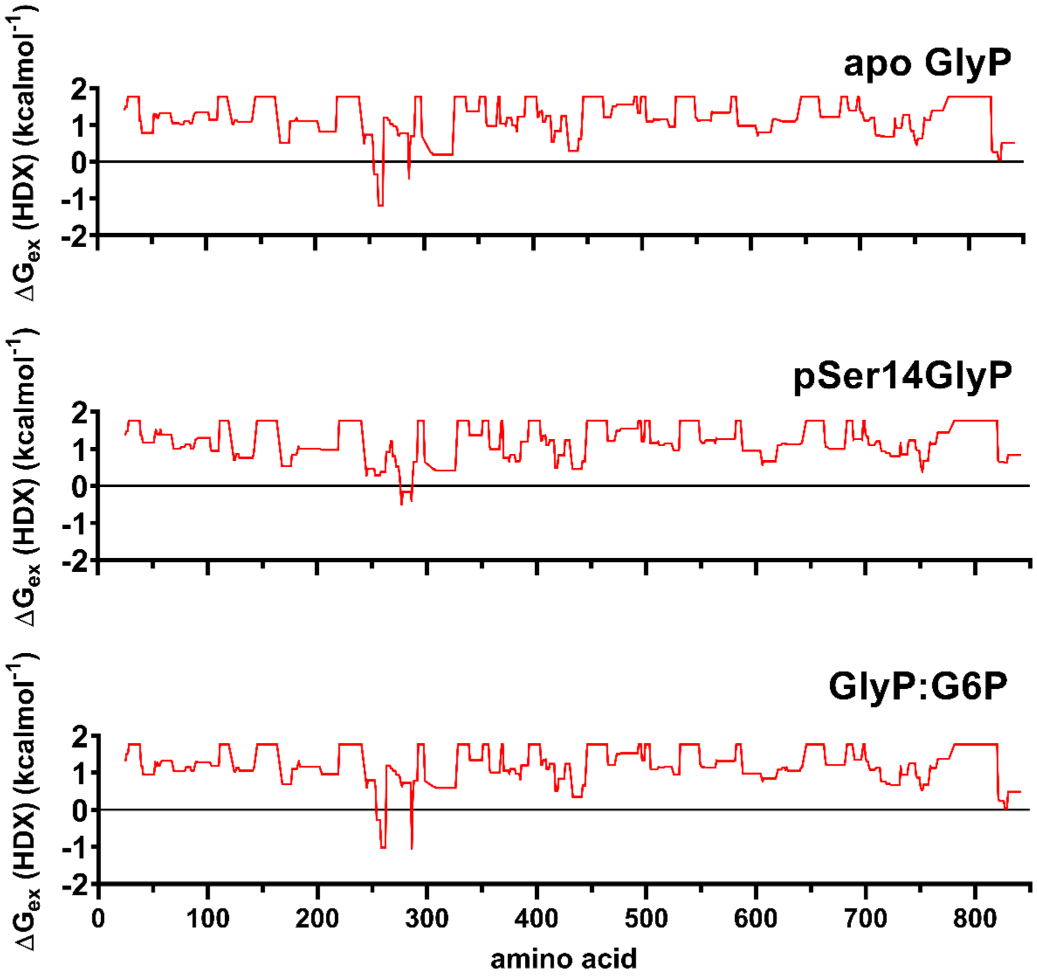


*Figure S 9. Estimates of the Gibbs free energy of stability $\Delta G_{ex}$(HDX) per amino acid in GlyP in three states: apo GlyPb (top), pSer14 GlyPa (middle) and GlyP:G6P inactive complex (bottom). Values calculated from fitted exchange rate constants using Equation S6. 2 kcal/mol is the upper limit of quantitation, given the very slow HDX rates of strongly protected amide protons. (1)*

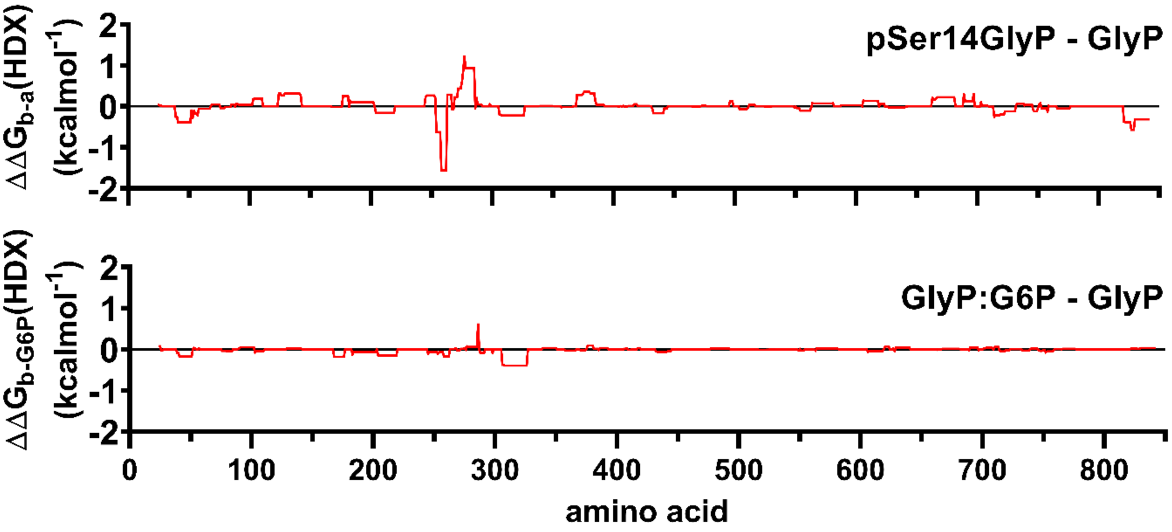


*Figure S 10. Estimates of the change in free energy of stability $\Delta\Delta G_{ex}$(HDX) per amino acid in GlyP upon activation by phosphorylation (top) and inhibition by G6P (bottom).*

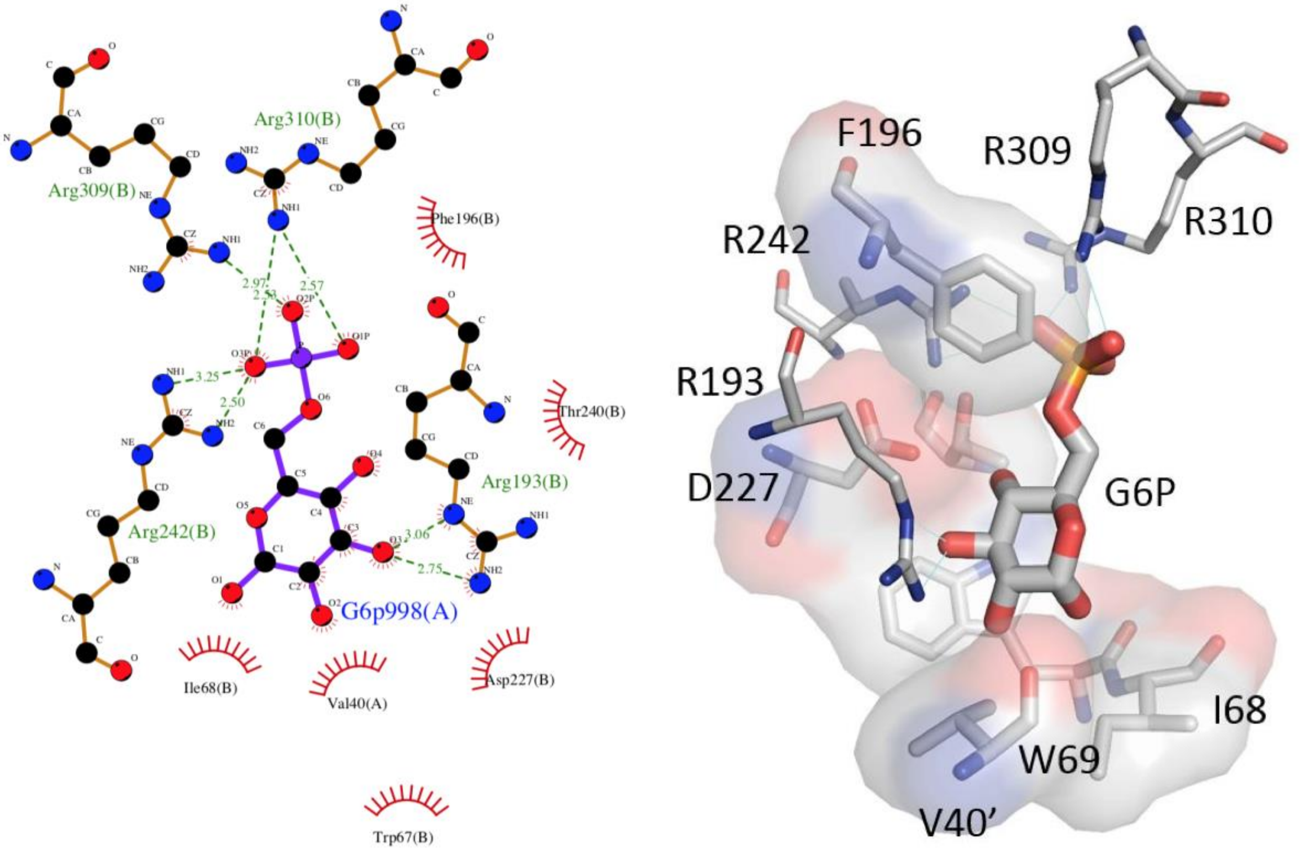


*Figure S 11. Interactions of G6P inhibitor with GlyP, from 1GPY.pdb. Created with Ligplot+ v2.1 (left) and Pymol (right - Schrodinger). The majority of interactions between G6P and GlyP are with the monomer that forms the nucleotide site and a single van der Waals interaction is identified between the G6P and the opposing monomer at Val40’.*

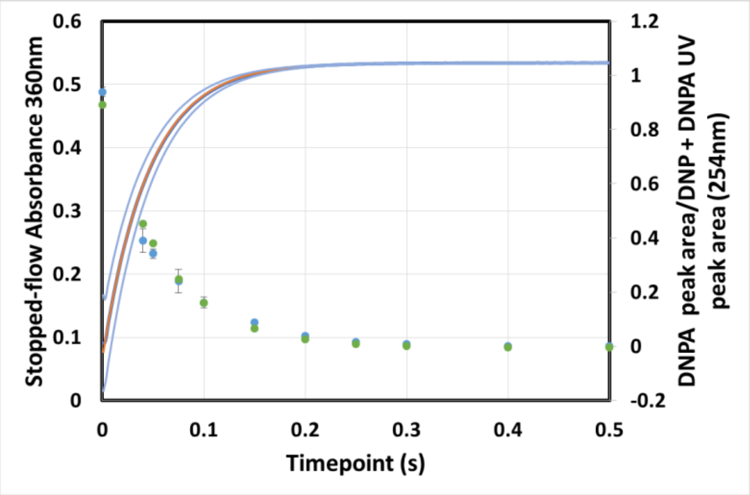


*Figure S 12. Kinetic data for the hydrolysis of DNPA to DNP in the ms2min prototype (blue dots, error bars are ±2 s.d. from n=3), and SX stopped-flow spectrometer (dark blue line, light blue lines are ±2 s.d. from n=9). The data was fitted to a single exponential to provide a first order rate constant for the ms2min (green dots) of 20.1 $s^{-1}$ (95% confidence interval range 16.9 to 23.4 $s^{-1}$) and SX20 (orange line) of 21.85 $s^{-1}$ (95% confidence interval range of 21.62 to 22.09 $s^{-1}$ ).*

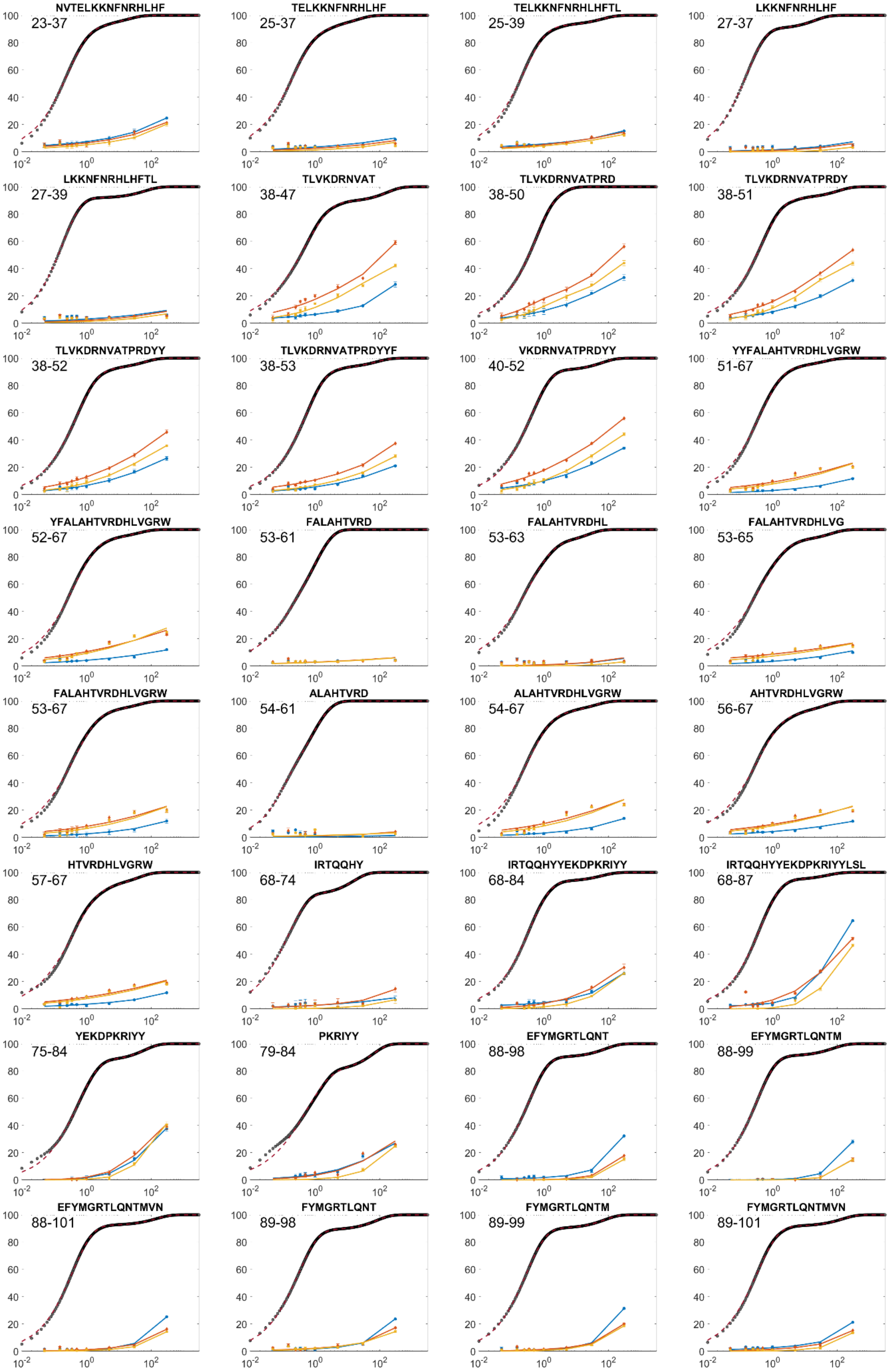
NVTELKKNFNRHLHF
23-37
TELKKNFNRHLHF
25-37
TELKKNFNRHLHFTL
25-39
LKKNFNRHLHF
27-37
LKKNFNRHLHFTL
27-39
TLVKDRNVAT
38-47
TLVKDRNVATPRD
38-50
TLVKDRNVATPRDY
38-51
TLVKDRNVATPRDYY
38-52
TLVKDRNVATPRDYYF
38-53
VKDRNVATPRDYY
40-52
YYFALAHTVRDHLVGRW
51-67
YFALAHTVRDHLVGRW
52-67
FALAHTVRD
53-61
FALAHTVRDHL
53-63
FALAHTVRDHLVG
53-65
FALAHTVRDHLVGRW
53-67
ALAHTVRD
54-61
ALAHTVRDHLVGRW
54-67
AHTVRDHLVGRW
56-67
HTVRDHLVGRW
57-67
IRTQQHY
68-74
IRTQQHYYEKDPKRIYY
68-84
IRTQQHYYEKDPKRIYYLSL
68-87
YEKDPKRIYY
75-84
PKRIYY
79-84
EFYMGRTLQNT
88-98
EFYMGRTLQNTM
88-99
EFYMGRTLQNTMVN
88-101
FYMGRTLQNT
89-98
FYMGRTLQNTM
89-99
FYMGRTLQNTMVN
89-101

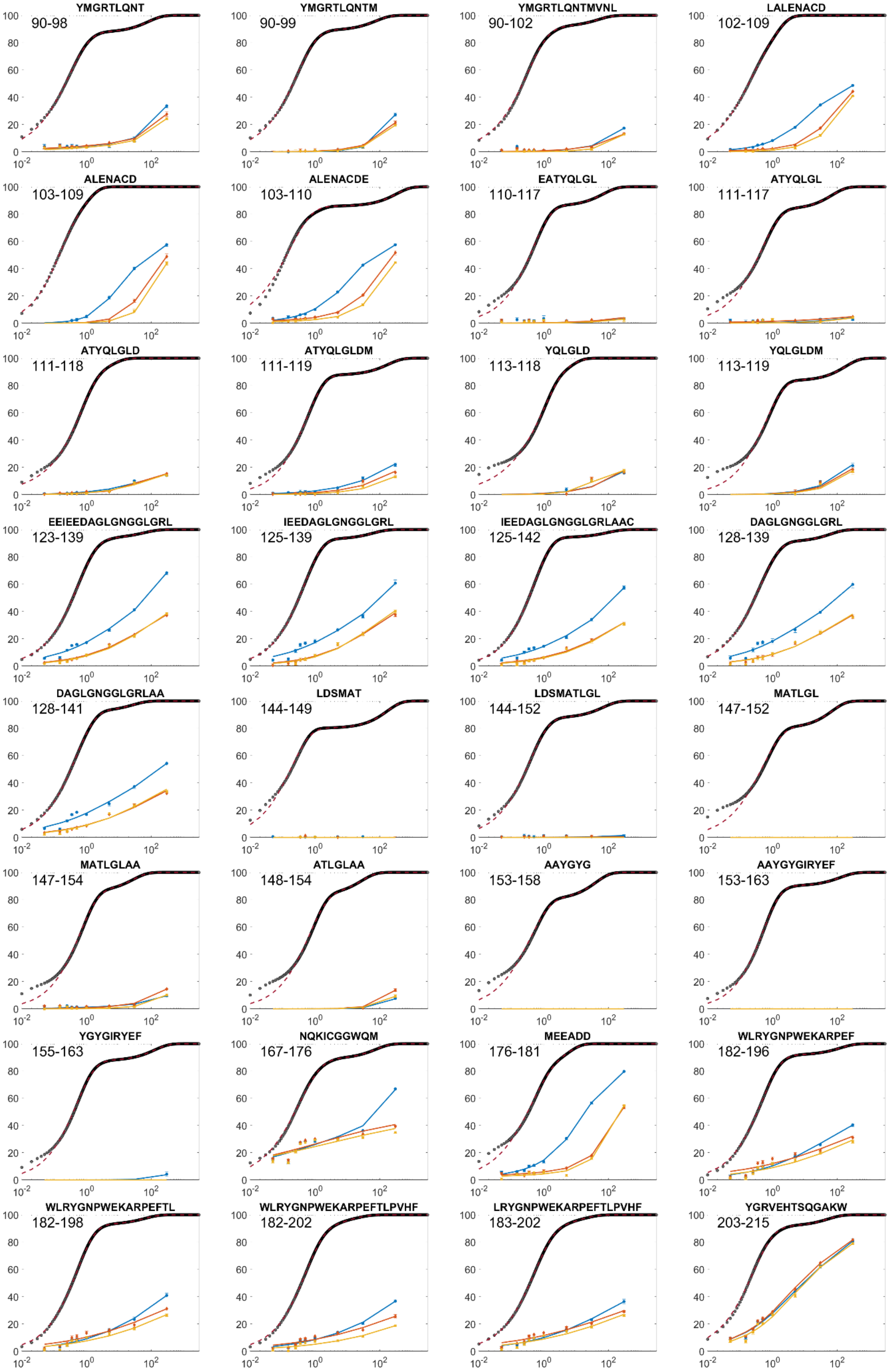
YMGRTLQNT
90-98
YMGRTLQNTM
90-99
YMGRTLQNTMVNL
90-102
LALENACD
102-109
ALENACD
103-109
ALENACDE
103-110
EATYQLGL
110-117
ATYQLGL
111-117
ATYQLGLD
111-118
ATYQLGLDM
111-119
YQLGLD
113-118
YQLGLDM
113-119
EEIEEDAGLGNGGLGRL
123-139
IEEDAGLGNGGLGRL
125-139
IEEDAGLGNGGLGRLAAC
125-142
DAGLGNGGLGRL
128-139
DAGLGNGGLGRLAA
128-141
LDSMAT
144-149
LDSMATLGL
144-152
MATLGL
147-152
MATLGLAA
147-154
ATLGLAA
148-154
AAYGYG
153-158
AAYGYGIRYEF
153-163
YGYGIRYEF
155-163
NQKICGGWQM
167-176
MEEADD
176-181
WLRYGNPWEKARPEF
182-196
WLRYGNPWEKARPEFTL
182-198
WLRYGNPWEKARPEFTLPVHF
182-202
LRYGNPWEKARPEFTLPVHF
183-202
YGRVEHTSQGAKW
203-215

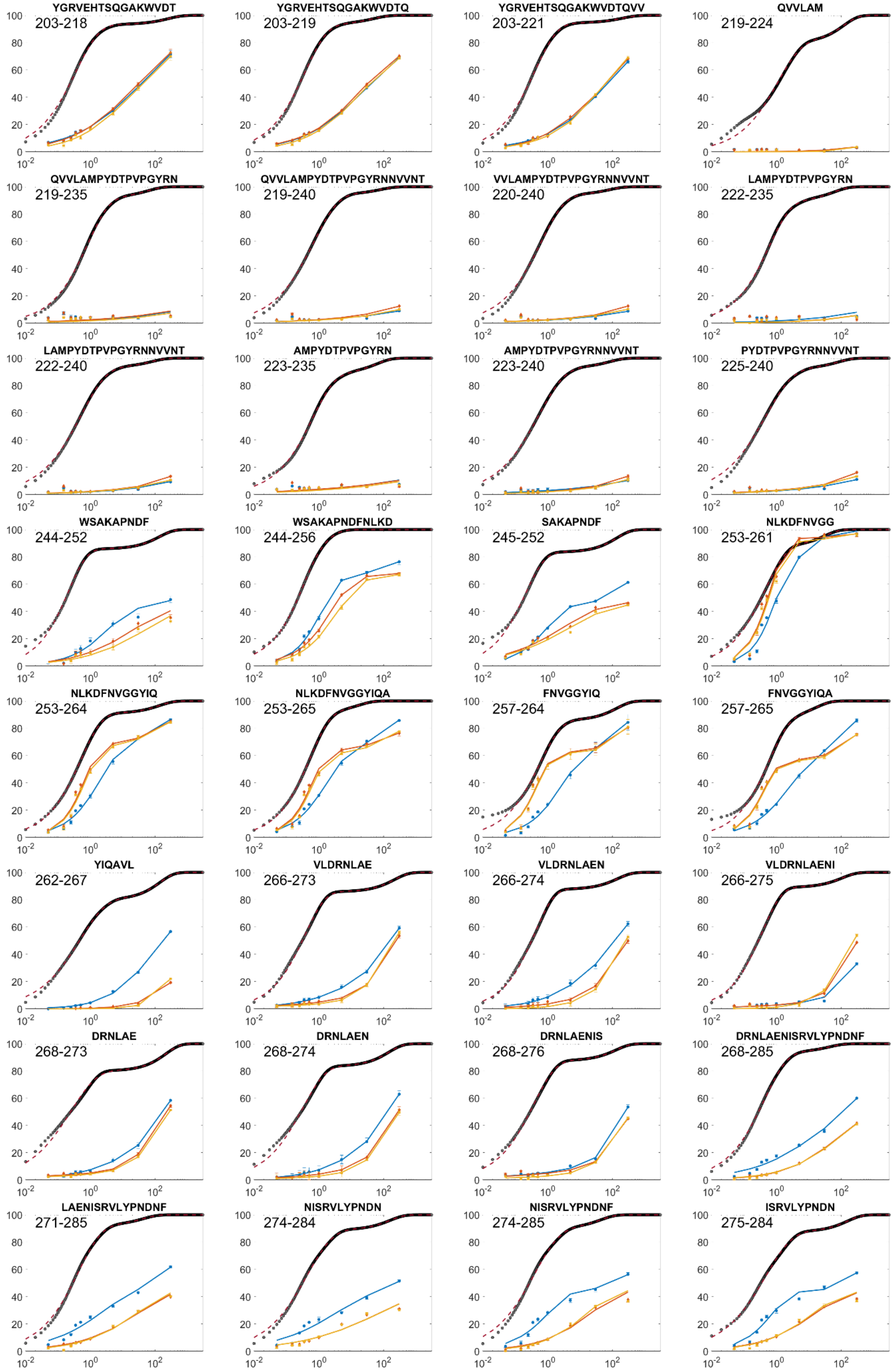

YGRVEHTSQGAKWVDT
203-218
YGRVEHTSQGAKWVDTQ
203-219
YGRVEHTSQGAKWVDTQVV
203-221
QVVLAM
219-224
QVVLAMPYDTPVPGYRN
219-235
QVVLAMPYDTPVPGYRNNVVNT
219-240
VVLAMPYDTPVPGYRNNVVNT
220-240
LAMPYDTPVPGYRN
222-235
LAMPYDTPVPGYRNNVVNT
222-240
AMPYDTPVPGYRN
223-235
AMPYDTPVPGYRNNVVNT
223-240
PYDTPVPGYRNNVVNT
225-240
WSAKAPNDF
244-252
WSAKAPNDFNLKD
244-256
SAKAPNDF
245-252
NLKDFNVGG
253-261
NLKDFNVGGYIQ
253-264
NLKDFNVGGYIQA
253-265
FNVGGYIQ
257-264
FNVGGYIQA
257-265
YIQAVL
262-267
VLDRNLAE
266-273
VLDRNLAEN
266-274
VLDRNLAENI
266-275
DRNLAE
268-273
DRNLAEN
268-274
DRNLAENIS
268-276
DRNLAENISRVLYPNDNF
268-285
LAENISRVLYPNDNF
271-285
NISRVLYPNDN
274-284
NISRVLYPNDNF
274-285
ISRVLYPNDN
275-284

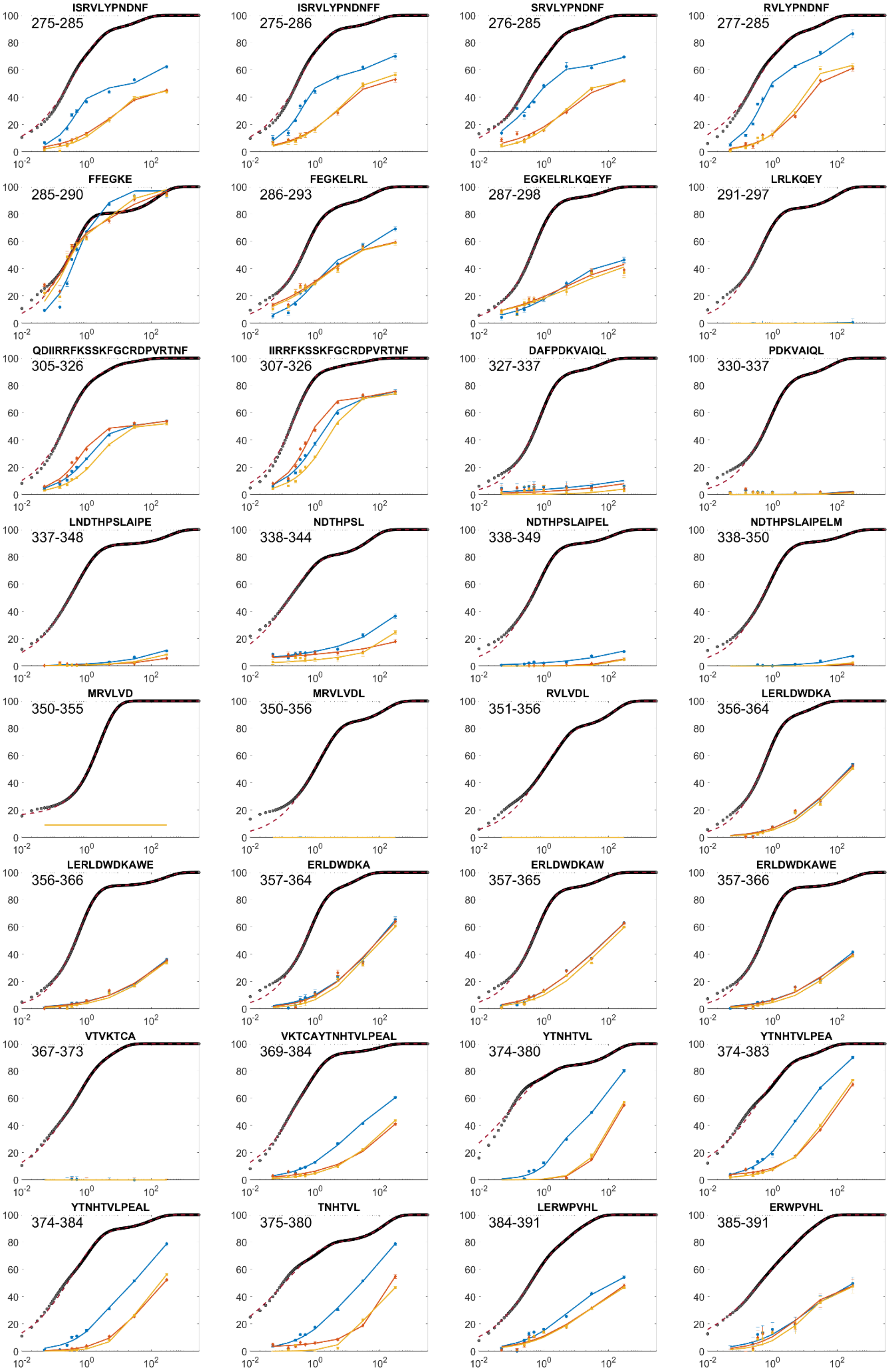
ISRVLYPNDNF
275-285
ISRVLYPNDNFF
275-286
SRVLYPNDNF
276-285
RVLYPNDNF
277-285
FFEGKE
285-290
FEGKELRL
286-293
EGKELRLKQEYF
287-298
LRLKQEY
291-297
QDIIRRFKSSKFGCRDPVRTNF
305-326
IIRRFKSSKFGCRDPVRTNF
307-326
DAFPDKVAIQL
327-337
PDKVAIQL
330-337
LNDTHPSLAIPE
337-348
NDTHPSL
338-344
NDTHPSLAIPEL
338-349
NDTHPSLAIPELM
338-350
MRVLVD
350-355
MRVLVDL
350-356
RVLVDL
351-356
LERLDWDKA
356-364
LERLDWDKAWE
356-366
ERLDWDKA
357-364
ERLDWDKAW
357-365
ERLDWDKAWE
357-366
VTVKTCA
367-373
VKTCAYTNHTVLPEAL
369-384
YTNHTVL
374-380
YTNHTVLPEA
374-383
YTNHTVLPEAL
374-384
TNHTVL
375-380
LERWPVHL
384-391
ERWPVHL
385-391
0
20
40
60
80
100
$10^{-2}$
$10^{0}$
$10^{2}$

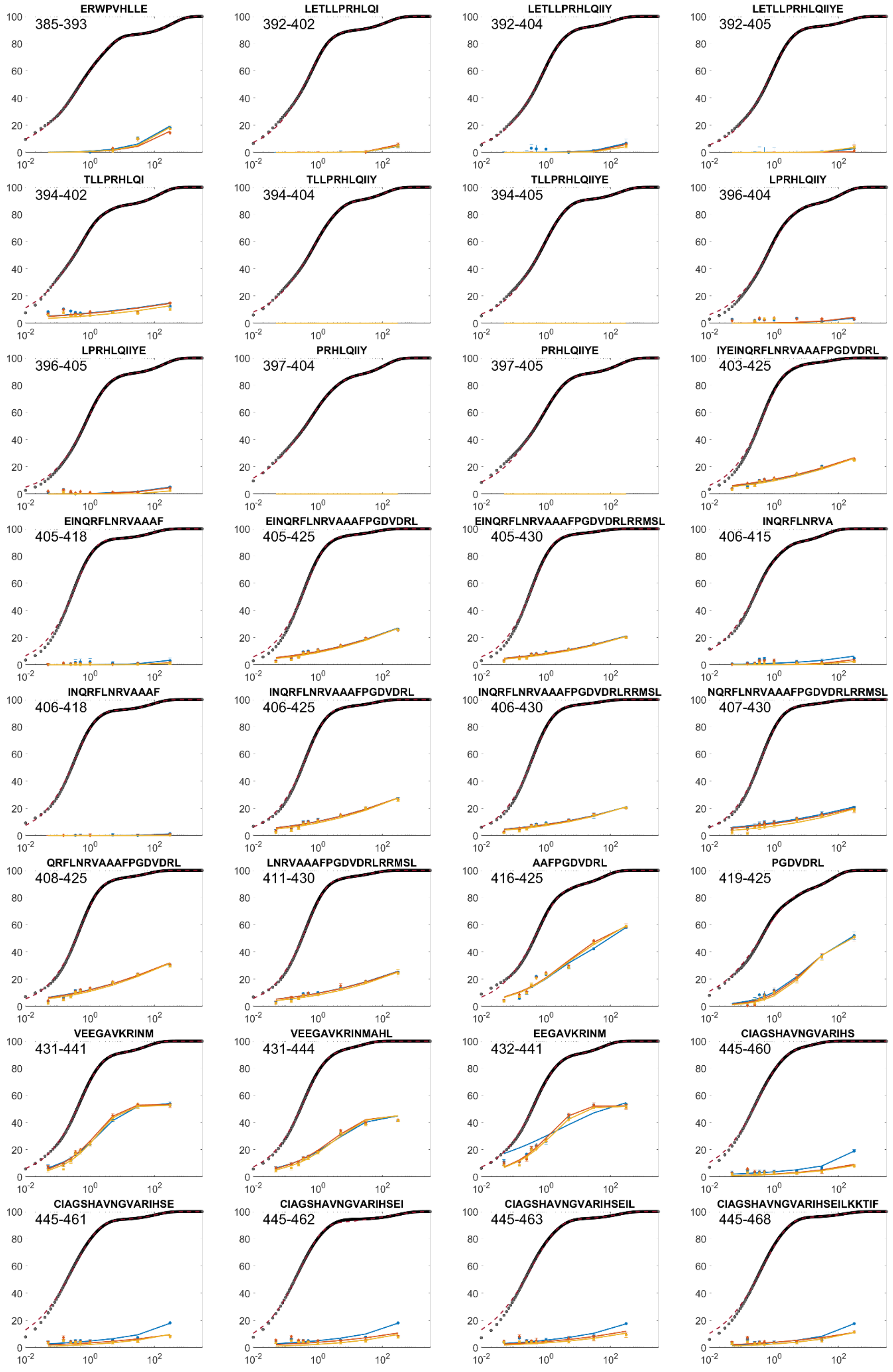
ERWPVHLLE
385-393
LETLLPRHLQI
392-402
LETLLPRHLQIIY
392-404
LETLLPRHLQIIYE
392-405
TLLPRHLQI
394-402
TLLPRHLQIIY
394-404
TLLPRHLQIIYE
394-405
LPRHLQIIY
396-404
LPRHLQIIYE
396-405
PRHLQIIY
397-404
PRHLQIIYE
397-405
IYEINQRFLNRVAAAFPGDVDRL
403-425
EINQRFLNRVAAAF
405-418
EINQRFLNRVAAAFPGDVDRL
405-425
EINQRFLNRVAAAFPGDVDRLRRMSL
405-430
INQRFLNRVA
406-415
INQRFLNRVAAAF
406-418
INQRFLNRVAAAFPGDVDRL
406-425
INQRFLNRVAAAFPGDVDRLRRMSL
406-430
NQRFLNRVAAAFPGDVDRLRRMSL
407-430
QRFLNRVAAAFPGDVDRL
408-425
LNRVAAAFPGDVDRLRRMSL
411-430
AAFPGDVDRL
416-425
PGDVDRL
419-425
VEEGAVKRINM
431-441
VEEGAVKRINMAHL
431-444
EEGAVKRINM
432-441
CIAGSHAVNGVARIHS
445-460
CIAGSHAVNGVARIHSE
445-461
CIAGSHAVNGVARIHSEI
445-462
CIAGSHAVNGVARIHSEIL
445-463
CIAGSHAVNGVARIHSEILKKTIF
445-468

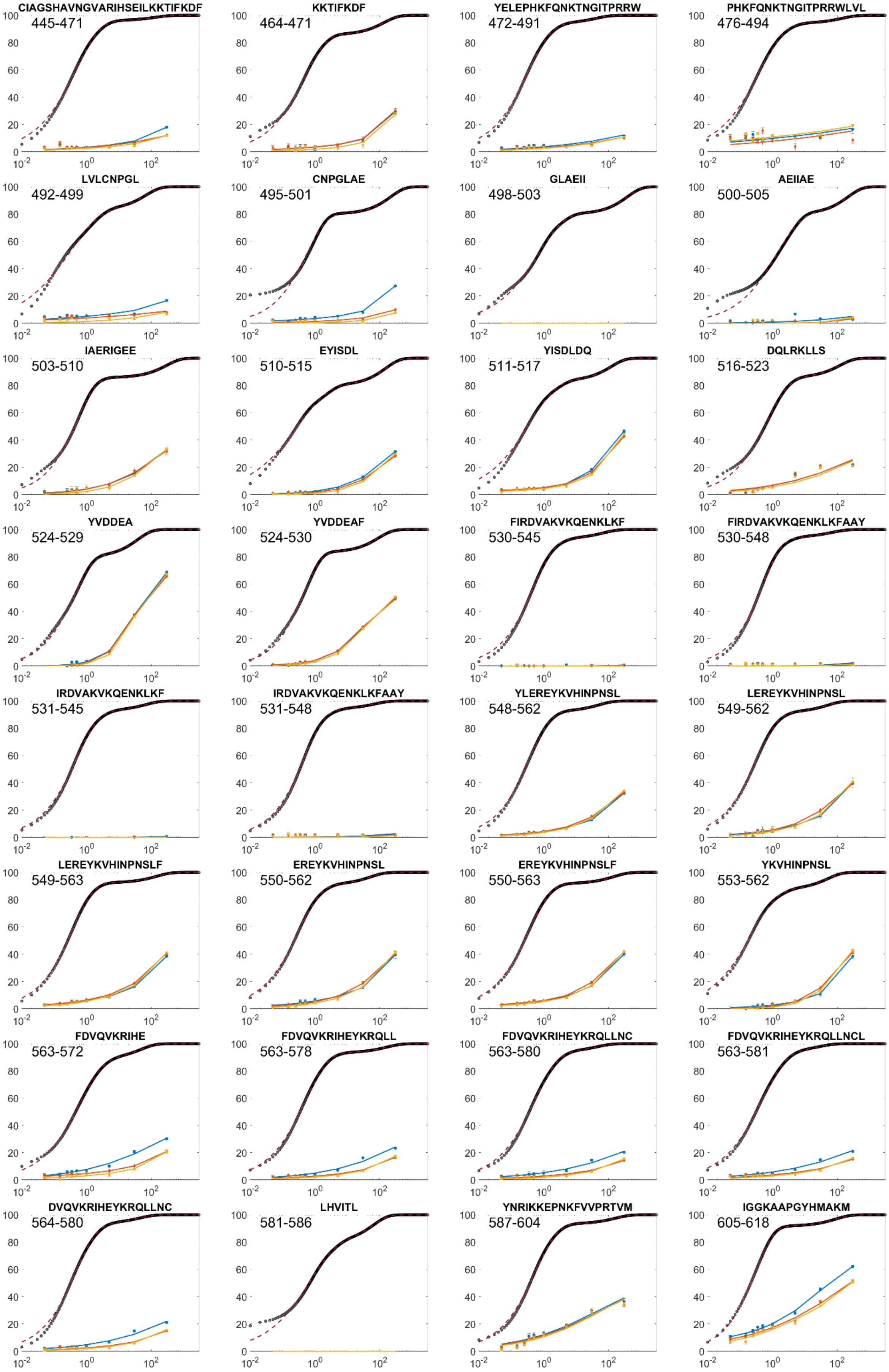
CIAGSHAVNGVARIHSEILKKTIFKDF
445-471
KKTIFKDF
464-471
YELEPHKFQNKTNGITPRRW
472-491
PHKFQNKTNGITPRRWLVL
476-494
LVLCNPGL
492-499
CNPGLAE
495-501
GLAEII
498-503
AEIIAE
500-505
IAERIGEE
503-510
EYISDL
510-515
YISDLDQ
511-517
DQLRKLLS
516-523
YVDDEA
524-529
YVDDEAF
524-530
FIRDVAKVKQENKLKF
530-545
FIRDVAKVKQENKLKFAAY
530-548
IRDVAKVKQENKLKF
531-545
IRDVAKVKQENKLKFAAY
531-548
YLEREYKVHINPNSL
548-562
LEREYKVHINPNSL
549-562
LEREYKVHINPNSLF
549-563
EREYKVHINPNSL
550-562
EREYKVHINPNSLF
550-563
YKVHINPNSL
553-562
FDVQVKRIHE
563-572
FDVQVKRIHEYKRQLL
563-578
FDVQVKRIHEYKRQLLNC
563-580
FDVQVKRIHEYKRQLLNCL
563-581
DVQVKRIHEYKRQLLNC
564-580
LHVITL
581-586
YNRIKKEPNKFVVPRTVM
587-604
IGGKAAPGYHMAKM
605-618

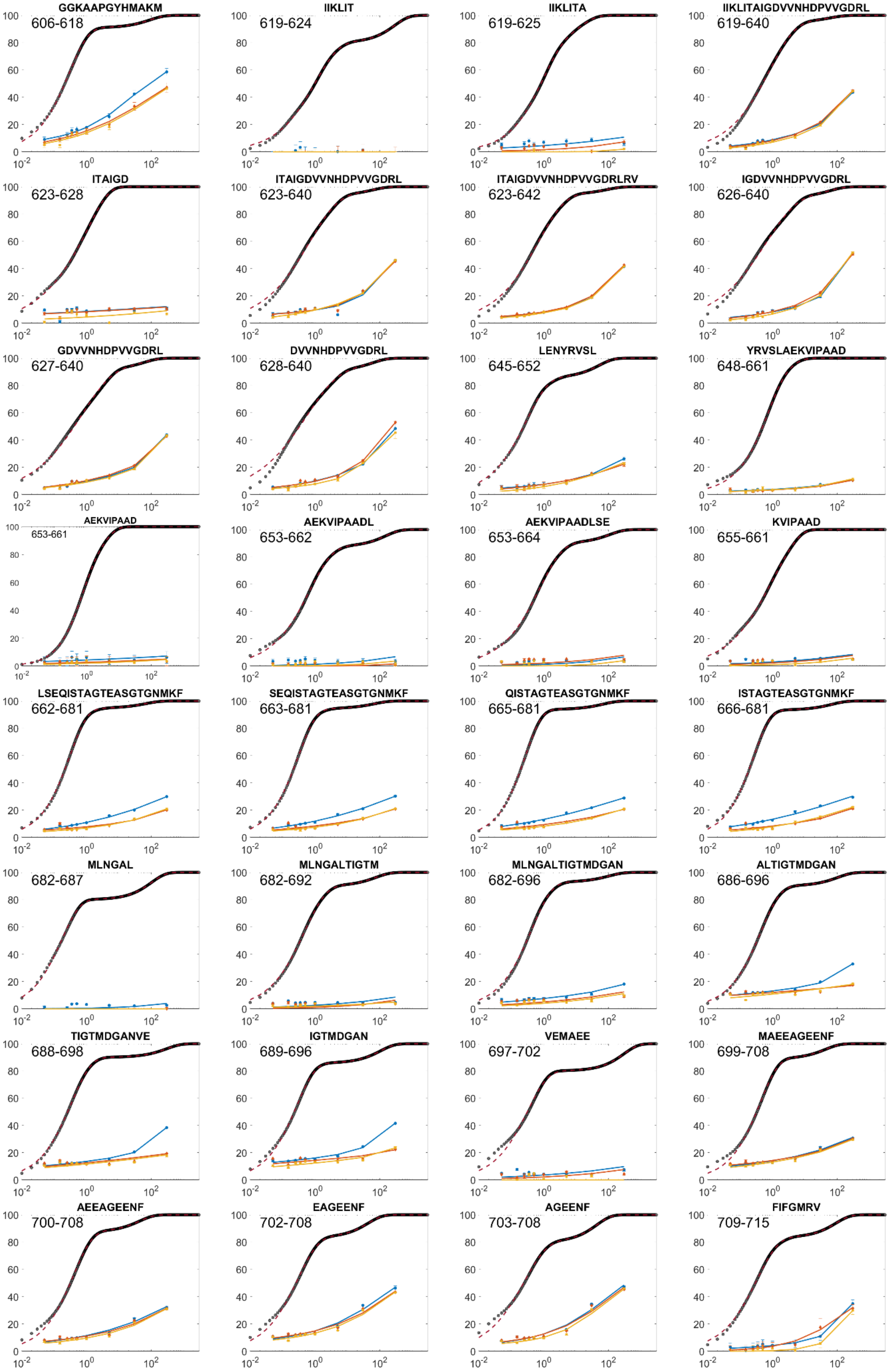
GGKAAPGYHMAKM
606-618
IIKLIT
619-624
IIKLITA
619-625
IIKLITAIGDVVNHDPVVGDRL
619-640
ITAIGD
623-628
ITAIGDVVNHDPVVGDRL
623-640
ITAIGDVVNHDPVVGDRLRV
623-642
IGDVVNHDPVVGDRL
626-640
GDVVNHDPVVGDRL
627-640
DVVNHDPVVGDRL
628-640
LENYRVSL
645-652
YRVSLAEKVIPAAD
648-661
AEKVIPAAD
653-661
AEKVIPAADL
653-662
AEKVIPAADLSE
653-664
KVIPAAD
655-661
LSEQISTAGTEASGTGNMKF
662-681
SEQISTAGTEASGTGNMKF
663-681
QISTAGTEASGTGNMKF
665-681
ISTAGTEASGTGNMKF
666-681
MLNGAL
682-687
MLNGALTIGTM
682-692
MLNGALTIGTMDGAN
682-696
ALTIGTMDGAN
686-696
TIGTMDGANVE
688-698
IGTMDGAN
689-696
VEMAEE
697-702
MAEEAGEENF
699-708
AEEAGEENF
700-708
EAGEENF
702-708
AGEENF
703-708
FIFGMRV
709-715

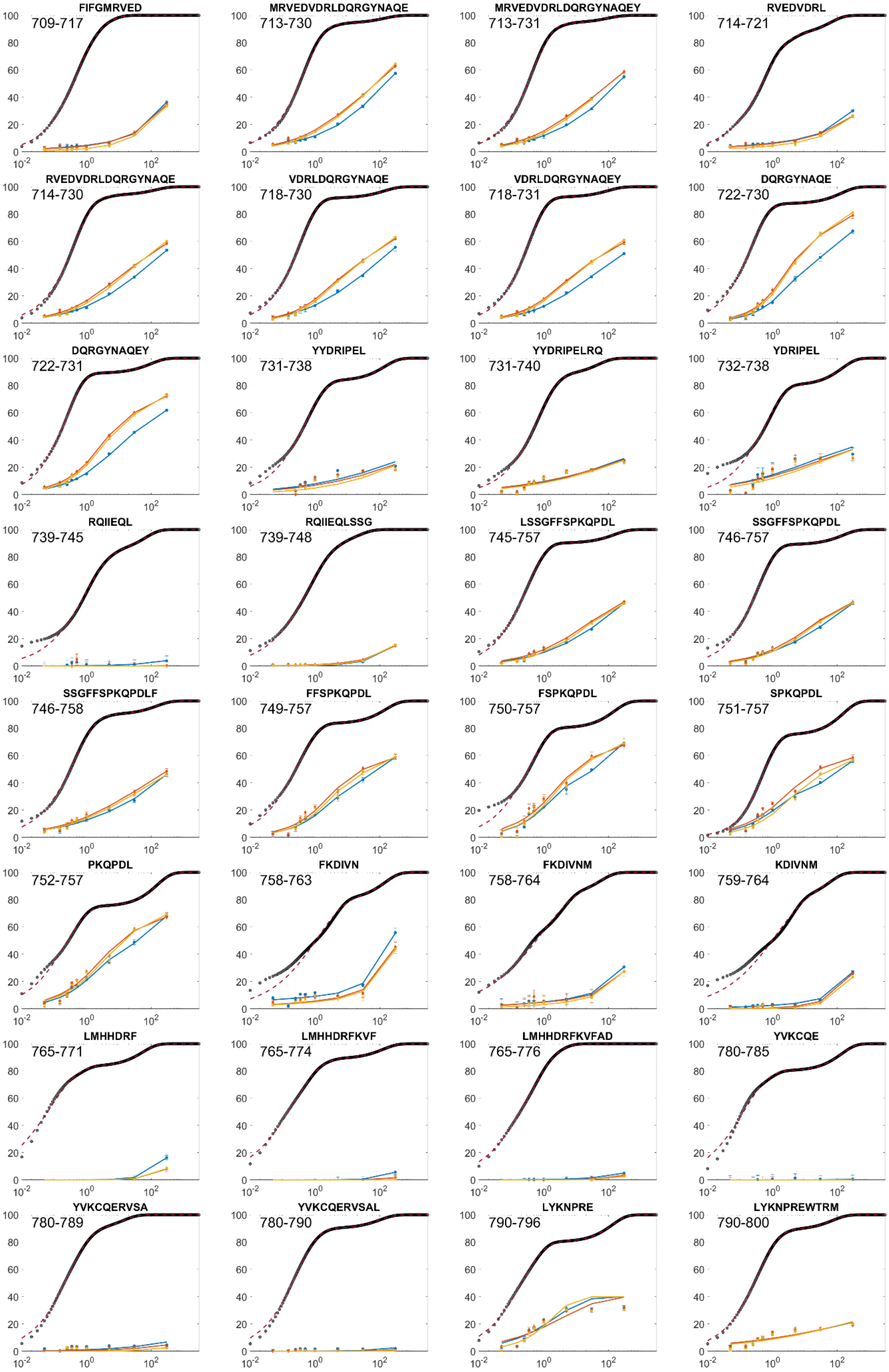
FIFGMRVED
709-717
MRVEDVDRLDQRGYNAQE
713-730
MRVEDVDRLDQRGYNAQEY
713-731
RVEDVDRL
714-721
RVEDVDRLDQRGYNAQE
714-730
VDRLDQRGYNAQE
718-730
VDRLDQRGYNAQEY
718-731
DQRGYNAQE
722-730
DQRGYNAQEY
722-731
YYDRIPEL
731-738
YYDRIPELRQ
731-740
YDRIPEL
732-738
RQIIEQL
739-745
RQIIEQLSSG
739-748
LSSGFFSPKQPDL
745-757
SSGFFSPKQPDL
746-757
SSGFFSPKQPDLF
746-758
FFSPKQPDL
749-757
FSPKQPDL
750-757
SPKQPDL
751-757
PKQPDL
752-757
FKDIVN
758-763
FKDIVNM
758-764
KDIVNM
759-764
LMHHDRF
765-771
LMHHDRFKVF
765-774
LMHHDRFKVFAD
765-776
YVKCQE
780-785
YVKCQERVSA
780-789
YVKCQERVSAL
780-790
LYKNPRE
790-796
LYKNPREWTRM
790-800

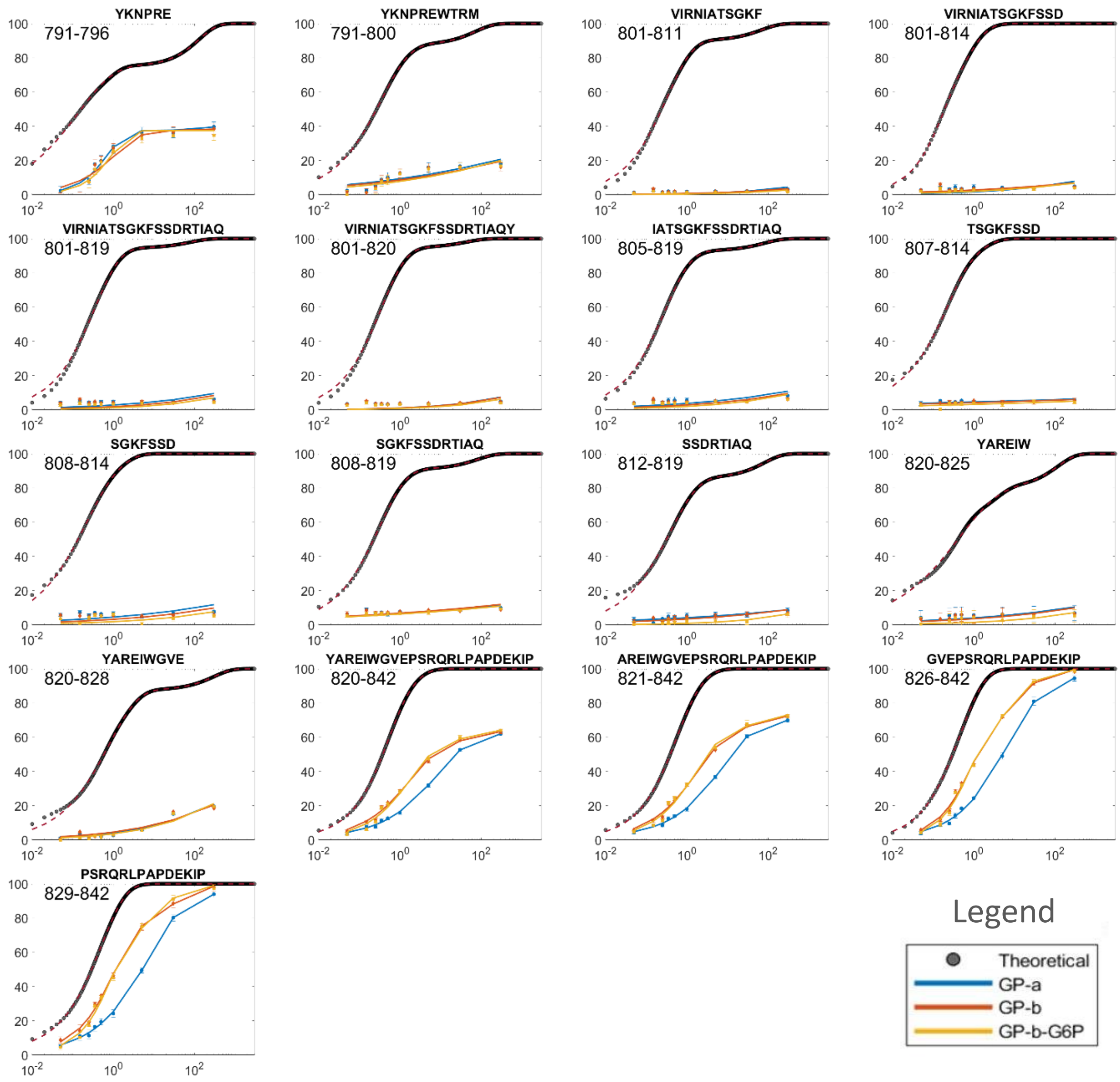


*Figure S 13. HDX analysis of GlyP per peptide. Back exchange corrected data for three protein states shown (red – apo GlyPb; yellow – G6P-bound inhibited GlyPb; blue – pSer14 activated GlyPa) against theoretical intrinsic exchange rates, calculated from [1]. Peptide sequence given above the plot, amino acid number in top-left. Y-axis shows absolute % deuteration at labeling time (s) on X-axis (labels omitted for space).*

*Table S4 Fitting parameters to multi-phase stretched exponential for hydrogen/deuterium-exchange mass spectrometry of GlyP in apo, activated and inactivated enzyme.*

| Sequence | | Intrinsic fits | | | | | | 'GP-a' | | | | | | GP-b' | | | | | | 'GP-b-G6P' | | | | | |
|---|---|---|---|---|---|---|---|---|---|---|---|---|---|---|---|---|---|---|---|---|---|---|---|---|---|
| | | $k_{int1}$ | $\beta_1$ | $k_{int2}$ | $N_1$ | $\beta_2$ | $N_2$ | $k_{exp1}$ | $\beta_1$ | $k_{exp2}$ | $N_1$ | $\beta_2$ | $N_2$ | $k_{exp1}$ | $\beta_1$ | $k_{exp2}$ | $N_1$ | $\beta_2$ | $N_2$ | $k_{exp1}$ | $\beta_1$ | $k_{exp2}$ | $N_1$ | $\beta_2$ | $N_2$ |
| 'NVTELKKNFNRHLHF' | 23 - 37 | 4E+00 | 0.7 | 8E-02 | 12 | 0.7 | 2 | 6E-08 | 0.4 | 2E-04 | 7 | 0.1 | 7 | 4E-08 | 0.4 | 1E-04 | 7 | 0.1 | 7 | 4E-05 | 1.0 | 3E-04 | 6 | 0.2 | 8 |
| 'TELKKNFNRHLHF' | 25 - 37 | 5E+00 | 0.7 | 8E-02 | 10 | 0.6 | 2 | 5E-06 | 1.0 | 5E-10 | 5 | 0.2 | 7 | 3E-06 | 1.0 | 3E-08 | 5 | 0.2 | 7 | 1E-05 | 1.0 | 8E-08 | 5 | 0.3 | 7 |
| 'TELKKNFNRHLHFTL' | 25 - 39 | 4E+00 | 0.9 | 2E-02 | 13 | 0.6 | 1 | 7E-08 | 0.5 | 3E-05 | 6 | 0.1 | 8 | 3E-05 | 1.0 | 1E-10 | 6 | 0.2 | 8 | 2E-05 | 1.0 | 1E-11 | 6 | 0.2 | 8 |
| 'LKKNFNRHLHF' | 27 - 37 | 6E+00 | 1.0 | 5E-02 | 9 | 0.7 | 1 | 1E-05 | 1.0 | 8E-08 | 4 | 0.3 | 6 | 9E-06 | 1.0 | 4E-14 | 4 | 0.3 | 6 | 4E-05 | 1.0 | 5E-07 | 4 | 0.6 | 6 |
| 'LKKNFNRHLHFTL' | 27 - 39 | 5E+00 | 1.0 | 2E-02 | 11 | 0.8 | 1 | 3E-06 | 1.0 | 2E-13 | 5 | 0.2 | 7 | 9E-06 | 0.9 | 6E-14 | 5 | 0.3 | 7 | 9E-06 | 1.0 | 4E-08 | 5 | 0.3 | 7 |
| 'TLVKDRNVAT' | 38 - 47 | 6E-03 | 0.6 | 2E+00 | 1 | 1.0 | 8 | 8E-04 | 0.2 | 6E-05 | 5 | 1.0 | 4 | 1E-03 | 0.3 | 1E-01 | 5 | 1.0 | 4 | 8E-05 | 0.4 | 4E-02 | 5 | 1.0 | 4 |
| 'TLVKDRNVATPRD' | 38 - 50 | 2E+00 | 0.6 | 2E+00 | 3 | 1.0 | 8 | 6E-03 | 0.3 | 6E-08 | 5 | 0.3 | 6 | 3E+00 | 0.4 | 1E-03 | 1 | 0.7 | 10 | 1E+00 | 0.4 | 4E-04 | 1 | 0.7 | 10 |
| 'TLVKDRNVATPRDY' | 38 - 51 | 1E-02 | 0.6 | 2E+00 | 1 | 0.9 | 11 | 4E-14 | 0.3 | 1E-03 | 5 | 1.0 | 7 | 1E-04 | 0.3 | 6E-02 | 7 | 0.4 | 5 | 7E-12 | 0.4 | 5E-02 | 6 | 1.0 | 6 |
| 'TLVKDRNVATPRDYY' | 38 - 52 | 2E+00 | 0.9 | 2E-02 | 12 | 0.7 | 1 | 2E-03 | 1.0 | 5E-11 | 6 | 0.3 | 7 | 3E-02 | 1.0 | 4E-04 | 6 | 0.3 | 7 | 1E-02 | 1.0 | 7E-14 | 6 | 0.4 | 7 |
| 'TLVKDRNVATPRDYYF' | 38 - 53 | 2E+00 | 0.9 | 1E-02 | 13 | 0.7 | 1 | 1E-04 | 1.0 | 3E-07 | 9 | 0.3 | 5 | 5E-03 | 1.0 | 5E-04 | 6 | 0.2 | 8 | 2E-03 | 1.0 | 1E-04 | 6 | 0.3 | 8 |
| 'VKDRNVATPRDYY' | 40 - 52 | 2E-02 | 0.7 | 2E+00 | 1 | 1.0 | 10 | 2E-08 | 0.3 | 3E-03 | 5 | 0.3 | 6 | 7E-04 | 0.3 | 1E-01 | 6 | 0.8 | 5 | 2E-04 | 0.4 | 3E-02 | 6 | 1.0 | 5 |
| 'YYFALAHTVRDHLVGRW' | 51 - 67 | 3E-02 | 0.6 | 2E+00 | 1 | 0.8 | 15 | 8E-05 | 0.2 | 1E-05 | 9 | 1.0 | 7 | 6E-13 | 0.2 | 6E-04 | 9 | 1.0 | 7 | 2E-11 | 0.2 | 7E-04 | 9 | 1.0 | 7 |
| 'YFALAHTVRDHLVGRW' | 52 - 67 | 3E-02 | 0.6 | 3E+00 | 1 | 0.8 | 14 | 3E-06 | 0.2 | 6E-06 | 8 | 1.0 | 7 | 7E-14 | 0.2 | 2E-03 | 8 | 1.0 | 7 | 4E-14 | 0.2 | 3E-03 | 8 | 1.0 | 7 |
| 'FALAHTVRD' | 53 - 61 | 1E+01 | 0.8 | 9E-01 | 3 | 0.6 | 5 | 4E-06 | 1.0 | 1E-12 | 2 | 0.2 | 6 | 4E-06 | 1.0 | 2E-13 | 2 | 0.2 | 6 | 4E-06 | 1.0 | 4E-13 | 2 | 0.2 | 6 |
| 'FALAHTVRDHL' | 53 - 63 | 3E+00 | 1.0 | 4E-02 | 9 | 0.6 | 1 | 8E-06 | 1.0 | 3E-12 | 5 | 0.3 | 5 | 1E-05 | 1.0 | 6E-08 | 5 | 0.3 | 5 | 4E-05 | 1.0 | 4E-07 | 5 | 0.6 | 5 |
| 'FALAHTVRDHLVG' | 53 - 65 | 2E+00 | 1.0 | 2E-02 | 11 | 0.5 | 1 | 1E-05 | 1.0 | 3E-07 | 5 | 0.2 | 7 | 1E-05 | 1.0 | 1E-11 | 5 | 0.1 | 7 | 3E-05 | 1.0 | 7E-14 | 5 | 0.2 | 7 |
| 'FALAHTVRDHLVGRW' | 53 - 67 | 2E+00 | 0.8 | 3E-02 | 13 | 0.6 | 1 | 3E-05 | 1.0 | 6E-05 | 6 | 0.3 | 8 | 6E-04 | 1.0 | 4E-14 | 6 | 0.2 | 8 | 8E-04 | 1.0 | 2E-14 | 6 | 0.3 | 8 |
| 'ALAHTVRD' | 54 - 61 | 1E+01 | 0.7 | 1E+00 | 2 | 1.0 | 5 | 7E-08 | 1.0 | 3E-14 | 1 | 0.2 | 6 | 3E-06 | 1.0 | 3E-05 | 1 | 0.2 | 6 | 1E-06 | 1.0 | 2E-08 | 1 | 0.2 | 6 |
| 'ALAHTVRDHLVGRW' | 54 - 67 | 3E+00 | 0.8 | 4E-02 | 12 | 0.6 | 1 | 3E-05 | 1.0 | 1E-04 | 6 | 0.3 | 7 | 3E-03 | 1.0 | 2E-14 | 6 | 0.2 | 7 | 3E-03 | 1.0 | 2E-14 | 6 | 0.3 | 7 |
| 'AHTVRDHLVGRW' | 56 - 67 | 3E+00 | 0.8 | 3E-02 | 10 | 0.6 | 1 | 8E-06 | 1.0 | 2E-06 | 5 | 0.2 | 6 | 6E-04 | 1.0 | 2E-14 | 5 | 0.2 | 6 | 8E-04 | 1.0 | 5E-14 | 5 | 0.2 | 6 |
| 'HTVRDHLVGRW' | 57 - 67 | 2E+00 | 0.8 | 3E-02 | 9 | 0.6 | 1 | 3E-06 | 0.9 | 1E-04 | 4 | 0.2 | 6 | 4E-04 | 1.0 | 3E-13 | 4 | 0.2 | 6 | 4E-04 | 1.0 | 3E-14 | 4 | 0.2 | 6 |
| 'IRTQQHY' | 68 - 74 | 7E+00 | 1.0 | 6E-02 | 5 | 0.7 | 1 | 2E-06 | 1.0 | 8E-08 | 3 | 0.2 | 3 | 2E-05 | 1.0 | 3E-04 | 3 | 0.3 | 3 | 1E-04 | 1.0 | 2E-06 | 3 | 0.5 | 3 |
| 'IRTQQHYYEKDPKRIYY' | 68 - 84 | 3E+00 | 1.0 | 2E-02 | 14 | 0.7 | 1 | 3E-08 | 0.6 | 9E-04 | 8 | 0.1 | 7 | 4E-03 | 1.0 | 8E-13 | 7 | 0.4 | 8 | 2E-03 | 1.0 | 2E-10 | 7 | 0.6 | 8 |
| 'IRTQQHYYEKDPKRIYYLSL' | 68 - 87 | 2E+00 | 1.0 | 2E-02 | 17 | 0.7 | 1 | 2E-02 | 0.1 | 3E-08 | 11 | 1.0 | 7 | 9E-03 | 1.0 | 4E-14 | 12 | 0.5 | 6 | 1E-02 | 1.0 | 5E-05 | 8 | 0.9 | 10 |
| 'YEKDPKRIYY' | 75 - 84 | 2E-02 | 0.7 | 2E+00 | 1 | 1.0 | 7 | 2E-14 | 0.7 | 9E-03 | 4 | 1.0 | 4 | 2E-11 | 0.7 | 1E-02 | 4 | 1.0 | 4 | 2E-14 | 1.0 | 1E-02 | 4 | 1.0 | 4 |
| 'PKRIYY' | 79 - 84 | 2E-02 | 0.6 | 2E+00 | 1 | 1.0 | 4 | 2E-14 | 0.4 | 4E-03 | 3 | 1.0 | 2 | 2E-14 | 0.5 | 5E-03 | 3 | 1.0 | 2 | 3E-14 | 0.7 | 3E-03 | 3 | 1.0 | 2 |

| | | | | | | | | | | | | | | | | | | | | | | | | | |
|---|---|---|---|---|---|---|---|---|---|---|---|---|---|---|---|---|---|---|---|---|---|---|---|---|---|
| 'EFYMGRTLQNT' | 88 - 98 | 1E-02 | 0.7 | 3E+00 | 1 | 1.0 | 9 | 2E-03 | 0.3 | 6E-06 | 6 | 1.0 | 5 | 2E-04 | 0.8 | 8E-04 | 5 | 0.9 | 5 | 1E-03 | 1.0 | 1E-07 | 5 | 1.0 | 5 |
| 'EFYMGRTLQNTM' | 88 - 99 | 1E-02 | 0.7 | 3E+00 | 1 | 1.0 | 10 | 3E-04 | 0.9 | 2E-03 | 6 | 1.0 | 5 | 6E-04 | 1.0 | 5E-04 | 6 | 1.0 | 5 | 9E-04 | 1.0 | 1E-04 | 6 | 1.0 | 5 |
| 'EFYMGRTLQNTMVN' | 88 - 101 | 7E-03 | 0.7 | 3E+00 | 1 | 1.0 | 12 | 8E-04 | 0.5 | 5E-04 | 7 | 1.0 | 6 | 4E-04 | 0.4 | 1E-04 | 7 | 1.0 | 6 | 3E-04 | 0.6 | 2E-04 | 7 | 1.0 | 6 |
| 'FYMGRTLQNT' | 89 - 98 | 1E-02 | 0.7 | 3E+00 | 1 | 1.0 | 8 | 1E-03 | 0.3 | 1E-05 | 5 | 1.0 | 4 | 5E-04 | 0.3 | 3E-05 | 5 | 1.0 | 4 | 7E-05 | 0.4 | 2E-04 | 5 | 1.0 | 4 |
| 'FYMGRTLQNTM' | 89 - 99 | 1E-02 | 0.7 | 3E+00 | 1 | 1.0 | 9 | 1E-03 | 0.6 | 9E-04 | 6 | 1.0 | 4 | 7E-04 | 0.4 | 9E-05 | 6 | 1.0 | 4 | 5E-07 | 0.7 | 1E-03 | 6 | 1.0 | 4 |
| 'FYMGRTLQNTMVN' | 89 - 101 | 7E-03 | 0.7 | 3E+00 | 1 | 1.0 | 11 | 9E-04 | 0.2 | 5E-06 | 7 | 1.0 | 5 | 3E-04 | 0.4 | 8E-05 | 7 | 1.0 | 5 | 4E-04 | 0.6 | 2E-04 | 7 | 1.0 | 5 |
| 'YMGRTLQNT' | 90 - 98 | 1E-02 | 0.7 | 4E+00 | 1 | 1.0 | 7 | 2E-03 | 0.2 | 2E-06 | 5 | 1.0 | 4 | 1E-03 | 0.2 | 7E-06 | 5 | 1.0 | 4 | 9E-04 | 0.2 | 2E-05 | 5 | 1.0 | 4 |
| 'YMGRTLQNTM' | 90 - 99 | 1E-02 | 0.7 | 4E+00 | 1 | 1.0 | 8 | 5E-04 | 0.9 | 2E-03 | 5 | 1.0 | 4 | 3E-04 | 0.7 | 1E-03 | 5 | 1.0 | 4 | 2E-14 | 0.8 | 2E-03 | 5 | 1.0 | 4 |
| 'YMGRTLQNTMVNL' | 90 - 102 | 3E+00 | 0.9 | 2E-02 | 11 | 0.7 | 1 | 3E-04 | 1.0 | 4E-04 | 5 | 0.5 | 7 | 2E-04 | 1.0 | 1E-04 | 5 | 0.5 | 7 | 9E-04 | 1.0 | 5E-07 | 5 | 0.8 | 7 |
| 'LALENACD' | 102 - 109 | 8E+00 | 0.9 | 9E-01 | 4 | 0.7 | 3 | 8E-02 | 0.6 | 2E-04 | 3 | 0.6 | 4 | 1E-02 | 0.2 | 5E-08 | 3 | 0.9 | 4 | 4E-03 | 1.0 | 5E-05 | 4 | 0.7 | 3 |
| 'ALENACD' | 103 - 109 | 1E+00 | 0.8 | 9E+00 | 2 | 0.9 | 4 | 1E-03 | 1.0 | 1E-01 | 4 | 1.0 | 2 | 8E-09 | 1.0 | 1E-02 | 3 | 1.0 | 3 | 4E-03 | 1.0 | 5E-10 | 4 | 0.9 | 2 |
| 'ALENACDE' | 103 - 110 | 3E-03 | 0.6 | 6E+00 | 1 | 1.0 | 6 | 7E-05 | 0.6 | 6E-02 | 3 | 0.8 | 4 | 1E-02 | 0.2 | 2E-07 | 3 | 0.8 | 4 | 8E-03 | 0.3 | 3E-06 | 3 | 1.0 | 4 |
| 'EATYQLGL' | 110 - 117 | 1E-02 | 0.7 | 2E+00 | 1 | 1.0 | 6 | 9E-08 | 0.4 | 1E-05 | 4 | 1.0 | 3 | 2E-07 | 0.4 | 2E-05 | 4 | 1.0 | 3 | 6E-07 | 0.5 | 2E-05 | 4 | 1.0 | 3 |
| 'ATYQLGL' | 111 - 117 | 1E-02 | 0.7 | 2E+00 | 1 | 1.0 | 5 | 4E-14 | 0.4 | 8E-06 | 3 | 1.0 | 3 | 2E-09 | 0.2 | 3E-07 | 3 | 1.0 | 3 | 3E-05 | 0.5 | 2E-05 | 3 | 1.0 | 3 |
| 'ATYQLGLD' | 111 - 118 | 1E+00 | 1.0 | 1E+00 | 5 | 0.5 | 2 | 2E-12 | 0.5 | 1E-02 | 6 | 1.0 | 1 | 2E-04 | 0.4 | 1E-03 | 6 | 0.9 | 1 | 5E-11 | 0.6 | 1E-02 | 6 | 1.0 | 1 |
| 'ATYQLGLDM' | 111 - 119 | 6E-03 | 0.8 | 2E+00 | 1 | 1.0 | 7 | 3E-14 | 0.4 | 2E-03 | 4 | 1.0 | 4 | 2E-14 | 0.5 | 7E-04 | 4 | 1.0 | 4 | 2E-14 | 0.5 | 5E-04 | 4 | 1.0 | 4 |
| 'YQLGLD' | 113 - 118 | 1E+00 | 1.0 | 9E-01 | 4 | 0.5 | 1 | 1E-03 | 1.0 | 2E-14 | 2 | 0.6 | 3 | 1E-03 | 1.0 | 3E-14 | 2 | 0.6 | 3 | 4E-11 | 1.0 | 2E-02 | 4 | 1.0 | 1 |
| 'YQLGLDM' | 113 - 119 | 6E-03 | 0.7 | 2E+00 | 1 | 1.0 | 5 | 2E-14 | 0.6 | 2E-03 | 3 | 1.0 | 3 | 2E-14 | 0.6 | 2E-03 | 3 | 1.0 | 3 | 7E-13 | 0.7 | 1E-03 | 3 | 1.0 | 3 |
| 'EEIEEDAGLGNGGLGRL' | 123 - 139 | 2E-02 | 0.7 | 2E+00 | 1 | 0.9 | 15 | 2E-03 | 0.4 | 1E-01 | 9 | 1.0 | 7 | 2E-14 | 0.4 | 1E-02 | 9 | 1.0 | 7 | 4E-05 | 0.4 | 1E-02 | 9 | 1.0 | 7 |
| 'IEEDAGLGNGGLGRL' | 125 - 139 | 2E+00 | 1.0 | 2E-02 | 13 | 0.7 | 1 | 1E-01 | 1.0 | 1E-03 | 7 | 0.4 | 7 | 1E-02 | 1.0 | 2E-14 | 7 | 0.4 | 7 | 2E-02 | 1.0 | 2E-12 | 7 | 0.5 | 7 |
| 'IEEDAGLGNGGLGRLAAC' | 125 - 142 | 1E-02 | 0.8 | 2E+00 | 1 | 1.0 | 16 | 1E-03 | 0.3 | 5E-02 | 9 | 1.0 | 8 | 1E-12 | 0.4 | 5E-03 | 9 | 1.0 | 8 | 2E-12 | 0.4 | 5E-03 | 9 | 1.0 | 8 |
| 'DAGLGNGGLGRL' | 128 - 139 | 2E+00 | 1.0 | 2E-02 | 10 | 0.7 | 1 | 2E-01 | 1.0 | 1E-03 | 5 | 0.4 | 6 | 1E-02 | 1.0 | 2E-12 | 5 | 0.4 | 6 | 2E-02 | 1.0 | 2E-14 | 5 | 0.4 | 6 |
| 'DAGLGNGGLGRLAA' | 128 - 141 | 2E+00 | 1.0 | 4E-02 | 12 | 0.7 | 1 | 1E-01 | 1.0 | 7E-04 | 6 | 0.3 | 7 | 9E-03 | 1.0 | 2E-14 | 6 | 0.3 | 7 | 1E-02 | 1.0 | 2E-14 | 6 | 0.3 | 7 |
| 'LDSMAT' | 144 - 149 | 6E-03 | 0.7 | 6E+00 | 1 | 1.0 | 4 | 2E-13 | 1.0 | 5E-09 | 3 | 1.0 | 2 | 2E-12 | 1.0 | 8E-08 | 3 | 1.0 | 2 | 2E-14 | 1.0 | 3E-09 | 3 | 1.0 | 2 |
| 'LDSMATLGL' | 144 - 152 | 1E-02 | 0.7 | 3E+00 | 1 | 1.0 | 7 | 7E-08 | 0.9 | 3E-05 | 4 | 0.4 | 4 | 3E-08 | 1.0 | 1E-08 | 3 | 0.4 | 5 | 2E-14 | 1.0 | 6E-10 | 4 | 1.0 | 4 |
| 'MATLGL' | 147 - 152 | 1E-02 | 0.7 | 2E+00 | 1 | 1.0 | 4 | 2E-14 | 1.0 | 1E-10 | 3 | 1.0 | 2 | 2E-14 | 1.0 | 3E-12 | 3 | 1.0 | 2 | 2E-14 | 1.0 | 1E-13 | 3 | 1.0 | 2 |
| 'MATLGLAA' | 147 - 154 | 2E+00 | 1.0 | 4E-02 | 6 | 0.8 | 1 | 5E-06 | 1.0 | 2E-04 | 3 | 0.3 | 4 | 2E-04 | 1.0 | 3E-04 | 3 | 0.5 | 4 | 2E-04 | 1.0 | 2E-04 | 3 | 0.7 | 4 |
| 'ATLGLAA' | 148 - 154 | 4E-02 | 0.7 | 2E+00 | 1 | 1.0 | 5 | 3E-04 | 1.0 | 2E-04 | 3 | 1.0 | 3 | 4E-04 | 1.0 | 6E-04 | 3 | 1.0 | 3 | 7E-06 | 1.0 | 8E-04 | 3 | 1.0 | 3 |
| 'AAYGYG' | 153 - 158 | 2E-02 | 0.7 | 3E+00 | 1 | 1.0 | 4 | 2E-14 | 1.0 | 4E-10 | 3 | 1.0 | 2 | 9E-14 | 1.0 | 1E-08 | 3 | 1.0 | 2 | 2E-14 | 1.0 | 4E-09 | 3 | 1.0 | 2 |
| 'AAYGYGIRYEF' | 153 - 163 | 8E-03 | 0.8 | 2E+00 | 1 | 1.0 | 9 | 2E-14 | 1.0 | 2E-12 | 6 | 1.0 | 4 | 2E-14 | 1.0 | 1E-12 | 6 | 1.0 | 4 | 2E-14 | 1.0 | 6E-10 | 6 | 1.0 | 4 |
| 'YGYGIRYEF' | 155 - 163 | 8E-03 | 0.7 | 2E+00 | 1 | 1.0 | 7 | 1E-04 | 1.0 | 1E-04 | 5 | 1.0 | 3 | 2E-14 | 1.0 | 4E-14 | 5 | 1.0 | 3 | 1E-13 | 1.0 | 3E-11 | 5 | 1.0 | 3 |
| 'NQKICGGWQM' | 167 - 176 | 1E-02 | 0.6 | 3E+00 | 1 | 1.0 | 8 | 2E-03 | 0.2 | 5E-01 | 5 | 1.0 | 4 | 2E-14 | 0.2 | 6E-01 | 5 | 1.0 | 4 | 6E-13 | 0.2 | 2E-01 | 5 | 1.0 | 4 |

| | | | | | | | | | | | | | | | | | | | | | | | | | |
|---|---|---|---|---|---|---|---|---|---|---|---|---|---|---|---|---|---|---|---|---|---|---|---|---|---|
| 'MEEADD' | 176 - 181 | 3E+00 | 1.0 | 2E-01 | 4 | 0.6 | 1 | 7E-02 | 0.3 | 4E-03 | 2 | 0.7 | 3 | 1E-07 | 0.9 | 6E-03 | 2 | 0.1 | 3 | 3E-08 | 1.0 | 6E-03 | 2 | 0.1 | 3 |
| 'WLRYGNPWEKARPEF' | 182 - 196 | 8E-03 | 0.7 | 2E+00 | 1 | 1.0 | 11 | 4E-05 | 0.4 | 2E-02 | 7 | 1.0 | 5 | 2E-14 | 0.2 | 7E-03 | 7 | 1.0 | 5 | 2E-14 | 0.3 | 4E-03 | 7 | 1.0 | 5 |
| 'WLRYGNPWEKARPEFTL' | 182 - 198 | 2E+00 | 1.0 | 2E-02 | 13 | 0.8 | 1 | 1E-02 | 1.0 | 2E-04 | 6 | 0.4 | 8 | 5E-03 | 1.0 | 7E-10 | 6 | 0.3 | 8 | 2E-03 | 1.0 | 4E-14 | 6 | 0.3 | 8 |
| 'WLRYGNPWEKARPEFTLPVHF' | 182 - 202 | 2E+00 | 0.6 | 1E-01 | 15 | 0.7 | 2 | 9E-03 | 1.0 | 3E-04 | 7 | 0.3 | 10 | 2E-03 | 1.0 | 1E-10 | 7 | 0.2 | 10 | 4E-04 | 1.0 | 2E-10 | 7 | 0.3 | 10 |
| 'LRYGNPWEKARPEFTLPVHF' | 183 - 202 | 1E-01 | 0.7 | 2E+00 | 2 | 0.7 | 14 | 7E-05 | 0.3 | 2E-02 | 9 | 1.0 | 7 | 1E-11 | 0.2 | 6E-03 | 9 | 1.0 | 7 | 2E-14 | 0.3 | 3E-03 | 9 | 1.0 | 7 |
| 'YGRVEHTSQGAKW' | 203 - 215 | 4E+00 | 1.0 | 2E-02 | 11 | 0.7 | 1 | 4E-01 | 0.4 | 4E-03 | 6 | 0.5 | 6 | 4E-01 | 0.4 | 3E-03 | 6 | 0.5 | 6 | 3E-01 | 0.5 | 2E-03 | 6 | 0.5 | 6 |
| 'YGRVEHTSQGAKWVDT' | 203 - 218 | 3E+00 | 1.0 | 4E-03 | 14 | 0.6 | 1 | 7E-02 | 0.4 | 1E-03 | 7 | 0.4 | 8 | 1E-01 | 0.5 | 1E-03 | 8 | 0.4 | 7 | 1E-01 | 0.7 | 1E-03 | 7 | 0.5 | 8 |
| 'YGRVEHTSQGAKWVDTQ' | 203 - 219 | 2E-02 | 0.6 | 3E+00 | 1 | 0.9 | 15 | 9E-04 | 0.4 | 9E-02 | 8 | 0.4 | 8 | 6E-04 | 0.5 | 7E-02 | 9 | 0.3 | 7 | 7E-04 | 0.5 | 1E-01 | 8 | 0.5 | 8 |
| 'YGRVEHTSQGAKWVDTQVV' | 203 - 221 | 5E-03 | 0.6 | 2E+00 | 1 | 1.0 | 17 | 3E-07 | 0.4 | 2E-02 | 6 | 0.2 | 12 | 2E-03 | 0.5 | 8E-02 | 9 | 1.0 | 9 | 6E-04 | 0.6 | 3E-02 | 10 | 0.3 | 8 |
| 'QVVLAM' | 219 - 224 | 8E-03 | 0.6 | 9E-01 | 1 | 1.0 | 4 | 1E-06 | 0.5 | 3E-05 | 3 | 1.0 | 2 | 2E-06 | 0.5 | 3E-05 | 3 | 1.0 | 2 | 4E-08 | 0.9 | 2E-04 | 3 | 1.0 | 2 |
| 'QVVLAMPYDTPVPGYRN' | 219 - 235 | 1E+00 | 0.9 | 2E-02 | 12 | 0.7 | 1 | 5E-06 | 1.0 | 9E-11 | 6 | 0.2 | 7 | 5E-06 | 1.0 | 1E-10 | 6 | 0.2 | 7 | 4E-06 | 1.0 | 3E-08 | 6 | 0.3 | 7 |
| 'QVVLAMPYDTPVPGYRNNVVNT' | 219 - 240 | 1E-02 | 0.6 | 1E+00 | 1 | 1.0 | 17 | 5E-07 | 0.2 | 3E-06 | 9 | 1.0 | 9 | 3E-05 | 0.3 | 3E-05 | 9 | 1.0 | 9 | 3E-05 | 0.3 | 1E-05 | 9 | 1.0 | 9 |
| 'VVLAMPYDTPVPGYRNNVVNT' | 220 - 240 | 1E-02 | 0.6 | 2E+00 | 1 | 0.9 | 16 | 3E-08 | 0.3 | 4E-06 | 8 | 1.0 | 9 | 2E-04 | 0.2 | 1E-05 | 9 | 1.0 | 8 | 2E-06 | 0.3 | 1E-05 | 8 | 1.0 | 9 |
| 'LAMPYDTPVPGYRN' | 222 - 235 | 2E-02 | 0.7 | 2E+00 | 1 | 0.9 | 9 | 4E-09 | 0.3 | 1E-05 | 6 | 1.0 | 4 | 8E-12 | 0.3 | 1E-05 | 6 | 1.0 | 4 | 3E-11 | 0.4 | 1E-05 | 6 | 1.0 | 4 |
| 'LAMPYDTPVPGYRNNVVNT' | 222 - 240 | 1E-02 | 0.6 | 2E+00 | 1 | 0.9 | 14 | 2E-05 | 0.3 | 2E-05 | 8 | 1.0 | 7 | 2E-04 | 0.3 | 2E-05 | 8 | 1.0 | 7 | 1E-05 | 0.3 | 2E-05 | 7 | 1.0 | 8 |
| 'AMPYDTPVPGYRN' | 223 - 235 | 2E-02 | 0.6 | 2E+00 | 1 | 0.9 | 8 | 2E-11 | 0.2 | 4E-06 | 5 | 1.0 | 4 | 2E-11 | 0.2 | 3E-06 | 5 | 1.0 | 4 | 1E-11 | 0.2 | 3E-06 | 5 | 1.0 | 4 |
| 'AMPYDTPVPGYRNNVVNT' | 223 - 240 | 2E+00 | 1.0 | 1E-02 | 13 | 0.5 | 1 | 8E-06 | 1.0 | 2E-09 | 7 | 0.2 | 7 | 5E-05 | 1.0 | 1E-04 | 6 | 0.3 | 8 | 1E-05 | 1.0 | 1E-04 | 6 | 0.3 | 8 |
| 'PYDTPVPGYRNNVVNT' | 225 - 240 | 1E-02 | 0.5 | 2E+00 | 1 | 0.9 | 12 | 1E-05 | 0.3 | 1E-05 | 6 | 1.0 | 7 | 3E-04 | 0.2 | 2E-05 | 7 | 1.0 | 6 | 4E-05 | 0.3 | 7E-05 | 7 | 1.0 | 6 |
| 'WSAKAPNDF' | 244 - 252 | 7E-03 | 0.7 | 5E+00 | 1 | 1.0 | 6 | 3E-04 | 0.6 | 3E-01 | 4 | 1.0 | 3 | 2E-14 | 0.4 | 4E-02 | 4 | 1.0 | 3 | 2E-14 | 0.4 | 2E-02 | 4 | 1.0 | 3 |
| 'WSAKAPNDFNLKD' | 244 - 256 | 3E+00 | 0.6 | 2E+00 | 3 | 1.0 | 8 | 8E-01 | 0.4 | 6E-04 | 7 | 0.9 | 4 | 4E-01 | 0.2 | 6E-07 | 7 | 0.7 | 4 | 2E-01 | 0.4 | 1E-05 | 7 | 0.7 | 4 |
| 'SAKAPNDF' | 245 - 252 | 7E-03 | 0.7 | 5E+00 | 1 | 1.0 | 5 | 1E-03 | 0.7 | 1E+00 | 3 | 0.9 | 3 | 3E-14 | 0.4 | 3E-01 | 3 | 1.0 | 3 | 1E-06 | 0.4 | 2E-01 | 3 | 0.8 | 3 |
| 'NLKDFNVGG' | 253 - 261 | 2E+00 | 1.0 | 3E-02 | 7 | 0.7 | 1 | 1E+00 | 1.0 | 6E-02 | 6 | 1.0 | 2 | 1E+00 | 0.5 | 2E-03 | 7 | 1.0 | 1 | 1E+00 | 0.5 | 6E-03 | 7 | 1.0 | 1 |
| 'NLKDFNVGGYIQ' | 253 - 264 | 7E-03 | 0.7 | 2E+00 | 1 | 1.0 | 10 | 4E-03 | 0.7 | 5E-01 | 4 | 0.5 | 7 | 2E-03 | 1.0 | 1E+00 | 4 | 0.6 | 7 | 3E-03 | 1.0 | 1E+00 | 4 | 0.5 | 7 |
| 'NLKDFNVGGYIQA' | 253 - 265 | 6E-02 | 0.7 | 2E+00 | 1 | 1.0 | 11 | 5E-03 | 0.7 | 6E-01 | 5 | 0.5 | 7 | 7E-04 | 1.0 | 2E+00 | 5 | 0.4 | 7 | 1E-03 | 1.0 | 2E+00 | 5 | 0.5 | 7 |
| 'FNVGGYIQ' | 257 - 264 | 7E-03 | 0.7 | 2E+00 | 1 | 1.0 | 6 | 4E-03 | 0.7 | 4E-01 | 3 | 0.6 | 4 | 2E-03 | 1.0 | 2E+00 | 3 | 0.8 | 4 | 2E-03 | 1.0 | 2E+00 | 3 | 1.0 | 4 |
| 'FNVGGYIQA' | 257 - 265 | 2E+00 | 1.0 | 6E-02 | 7 | 0.7 | 1 | 5E-01 | 0.5 | 6E-03 | 4 | 0.6 | 4 | 2E+00 | 0.7 | 2E-03 | 4 | 1.0 | 4 | 2E+00 | 0.9 | 2E-03 | 4 | 1.0 | 4 |
| 'YIQAVL' | 262 - 267 | 7E-03 | 0.6 | 2E+00 | 1 | 1.0 | 4 | 1E-03 | 0.6 | 3E-02 | 3 | 1.0 | 2 | 9E-07 | 0.7 | 2E-03 | 3 | 1.0 | 2 | 7E-04 | 1.0 | 9E-04 | 3 | 1.0 | 2 |
| 'VLDRNLAE' | 266 - 273 | 5E-03 | 0.7 | 3E+00 | 1 | 1.0 | 6 | 2E-03 | 0.4 | 2E-02 | 4 | 1.0 | 3 | 7E-03 | 0.2 | 2E-05 | 3 | 1.0 | 4 | 8E-03 | 0.2 | 1E-07 | 4 | 1.0 | 3 |
| 'VLDRNLAEN' | 266 - 274 | 7E-03 | 0.7 | 2E+00 | 1 | 1.0 | 7 | 1E-03 | 0.5 | 5E-02 | 4 | 1.0 | 4 | 7E-03 | 0.4 | 3E-04 | 2 | 1.0 | 6 | 2E-03 | 0.6 | 5E-03 | 4 | 0.9 | 4 |
| 'VLDRNLAENI' | 266 - 275 | 9E-03 | 0.7 | 3E+00 | 1 | 1.0 | 8 | 2E-03 | 0.2 | 5E-06 | 5 | 1.0 | 4 | 5E-03 | 0.2 | 4E-07 | 5 | 1.0 | 4 | 2E-03 | 0.7 | 5E-03 | 5 | 0.9 | 4 |
| 'DRNLAE' | 268 - 273 | 5E-03 | 0.5 | 4E+00 | 1 | 1.0 | 4 | 2E-03 | 0.4 | 2E-02 | 3 | 1.0 | 2 | 8E-03 | 0.1 | 5E-08 | 2 | 0.9 | 3 | 7E-03 | 0.2 | 1E-07 | 2 | 0.9 | 3 |
| 'DRNLAEN' | 268 - 274 | 7E-03 | 0.6 | 3E+00 | 1 | 1.0 | 5 | 2E-03 | 0.5 | 3E-02 | 3 | 1.0 | 3 | 6E-03 | 0.3 | 1E-04 | 2 | 1.0 | 4 | 7E-03 | 0.3 | 3E-05 | 3 | 1.0 | 3 |

| | | | | | | | | | | | | | | | | | | | | | | | | | |
|---|---|---|---|---|---|---|---|---|---|---|---|---|---|---|---|---|---|---|---|---|---|---|---|---|---|
| 'DRNLAENIS' | 268 - 276 | 8E-03 | 0.6 | 3E+00 | 1 | 1.0 | 7 | 4E-03 | 0.2 | 1E-04 | 4 | 1.0 | 4 | 3E-03 | 0.2 | 2E-05 | 4 | 1.0 | 4 | 5E-03 | 0.2 | 2E-07 | 4 | 0.9 | 4 |
| 'DRNLAENISRVLYPNDNF' | 268 - 285 | 2E-02 | 0.6 | 2E+00 | 1 | 1.0 | 15 | 1E-03 | 0.4 | 9E-02 | 9 | 1.0 | 7 | 2E-05 | 0.5 | 2E-02 | 9 | 0.9 | 7 | 4E-07 | 0.5 | 1E-02 | 8 | 1.0 | 8 |
| 'LAENISRVLYPNDNF' | 271 - 285 | 2E+00 | 1.0 | 2E-02 | 12 | 0.6 | 1 | 4E-01 | 1.0 | 1E-03 | 6 | 0.4 | 7 | 3E-02 | 1.0 | 3E-14 | 6 | 0.4 | 7 | 3E-02 | 1.0 | 1E-13 | 6 | 0.4 | 7 |
| 'NISRVLYPNDN' | 274 - 284 | 6E-03 | 0.6 | 2E+00 | 1 | 1.0 | 8 | 4E-04 | 0.4 | 3E-01 | 5 | 1.0 | 4 | 2E-14 | 0.3 | 1E-02 | 5 | 1.0 | 4 | 2E-14 | 0.3 | 1E-02 | 5 | 1.0 | 4 |
| 'NISRVLYPNDNF' | 274 - 285 | 2E+00 | 1.0 | 2E-02 | 9 | 0.6 | 1 | 9E-01 | 1.0 | 8E-04 | 4 | 0.6 | 6 | 4E-02 | 1.0 | 8E-14 | 4 | 0.5 | 6 | 6E-02 | 1.0 | 7E-14 | 4 | 0.5 | 6 |
| 'ISRVLYPNDN' | 275 - 284 | 6E-03 | 0.5 | 2E+00 | 1 | 1.0 | 7 | 9E-04 | 0.8 | 1E+00 | 4 | 1.0 | 4 | 7E-14 | 0.5 | 6E-02 | 4 | 1.0 | 4 | 4E-14 | 0.5 | 7E-02 | 4 | 1.0 | 4 |
| 'ISRVLYPNDNF' | 275 - 285 | 2E-02 | 0.5 | 2E+00 | 1 | 1.0 | 8 | 6E-04 | 1.0 | 2E+00 | 5 | 0.6 | 4 | 8E-07 | 0.5 | 1E-01 | 5 | 1.0 | 4 | 1E-12 | 0.6 | 1E-01 | 5 | 1.0 | 4 |
| 'ISRVLYPNDNFF' | 275 - 286 | 1E-02 | 0.6 | 2E+00 | 1 | 1.0 | 9 | 6E-04 | 1.0 | 2E+00 | 5 | 0.3 | 5 | 1E-06 | 0.5 | 1E-01 | 5 | 0.6 | 5 | 2E-07 | 0.5 | 1E-01 | 4 | 0.5 | 6 |
| 'SRVLYPNDNF' | 276 - 285 | 2E-02 | 0.5 | 2E+00 | 1 | 1.0 | 7 | 7E-04 | 0.5 | 2E+00 | 3 | 1.0 | 5 | 6E-07 | 0.4 | 1E-01 | 4 | 0.7 | 4 | 8E-05 | 0.5 | 2E-01 | 4 | 1.0 | 4 |
| 'RVLYPNDNF' | 277 - 285 | 2E-02 | 0.5 | 3E+00 | 1 | 1.0 | 6 | 6E-03 | 1.0 | 2E+00 | 3 | 0.5 | 4 | 5E-09 | 0.6 | 9E-02 | 3 | 0.4 | 4 | 1E-07 | 0.7 | 1E-01 | 3 | 0.4 | 4 |
| 'FFEGKE' | 285 - 290 | 6E-03 | 0.7 | 3E+00 | 1 | 1.0 | 4 | 2E+00 | 0.7 | 4E-01 | 3 | 1.0 | 2 | 2E-01 | 1.0 | 4E+00 | 3 | 0.3 | 2 | 2E-01 | 1.0 | 5E+00 | 3 | 0.4 | 2 |
| 'FEGKELRL' | 286 - 293 | 2E-02 | 0.6 | 2E+00 | 1 | 1.0 | 6 | 1E-03 | 0.7 | 1E+00 | 4 | 0.4 | 3 | 5E-07 | 0.3 | 4E-01 | 3 | 0.8 | 4 | 5E-07 | 0.4 | 4E-01 | 3 | 0.8 | 4 |
| 'EGKELRLKQEYF' | 287 - 298 | 1E-02 | 0.7 | 2E+00 | 1 | 1.0 | 10 | 7E-05 | 0.4 | 2E-01 | 6 | 1.0 | 5 | 2E-14 | 0.3 | 1E-01 | 6 | 1.0 | 5 | 2E-14 | 0.3 | 8E-02 | 6 | 1.0 | 5 |
| 'LRLKQEY' | 291 - 297 | 1E-02 | 0.7 | 3E+00 | 1 | 1.0 | 5 | 9E-08 | 1.0 | 3E-05 | 3 | 1.0 | 3 | 2E-14 | 1.0 | 1E-09 | 3 | 1.0 | 3 | 2E-14 | 1.0 | 3E-10 | 3 | 1.0 | 3 |
| 'QDIIRRFKSSKFGCRDPVRTNF' | 305 - 326 | 4E-01 | 0.7 | 4E+00 | 6 | 0.4 | 14 | 2E-05 | 0.7 | 7E-01 | 10 | 0.4 | 10 | 1E+00 | 0.3 | 2E-06 | 9 | 0.8 | 11 | 2E-05 | 0.7 | 3E-01 | 10 | 0.5 | 10 |
| 'IIRRFKSSKFGCRDPVRTNF' | 307 - 326 | 5E-02 | 0.7 | 4E+00 | 2 | 0.7 | 16 | 2E-04 | 0.7 | 7E-01 | 6 | 0.4 | 12 | 9E-05 | 0.8 | 1E+00 | 6 | 0.3 | 12 | 1E-04 | 0.7 | 3E-01 | 5 | 0.5 | 13 |
| 'DAFPDKVAIQL' | 327 - 337 | 2E-02 | 0.7 | 1E+00 | 1 | 1.0 | 8 | 4E-11 | 0.2 | 3E-06 | 5 | 1.0 | 4 | 3E-12 | 0.2 | 4E-06 | 5 | 1.0 | 4 | 9E-12 | 0.5 | 2E-05 | 5 | 1.0 | 4 |
| 'PDKVAIQL' | 330 - 337 | 2E-02 | 0.7 | 1E+00 | 1 | 1.0 | 6 | 1E-11 | 0.6 | 3E-05 | 4 | 1.0 | 3 | 4E-08 | 0.6 | 3E-05 | 4 | 1.0 | 3 | 3E-07 | 0.7 | 2E-05 | 4 | 1.0 | 3 |
| 'LNDTHPSLAIPE' | 337 - 348 | 3E-03 | 0.6 | 2E+00 | 1 | 1.0 | 8 | 2E-12 | 0.4 | 1E-04 | 5 | 1.0 | 4 | 6E-05 | 0.3 | 4E-06 | 5 | 1.0 | 4 | 6E-06 | 0.5 | 1E-04 | 5 | 1.0 | 4 |
| 'NDTHPSL' | 338 - 344 | 2E-02 | 0.5 | 6E+00 | 1 | 1.0 | 4 | 4E-04 | 0.1 | 7E-05 | 3 | 0.5 | 2 | 1E-04 | 0.1 | 1E-05 | 3 | 1.0 | 2 | 7E-04 | 0.2 | 3E-05 | 3 | 1.0 | 2 |
| 'NDTHPSLAIPEL' | 338 - 349 | 7E-03 | 0.6 | 2E+00 | 1 | 1.0 | 8 | 3E-07 | 0.3 | 4E-05 | 5 | 1.0 | 4 | 2E-06 | 0.6 | 1E-04 | 5 | 1.0 | 4 | 2E-04 | 1.0 | 6E-05 | 5 | 1.0 | 4 |
| 'NDTHPSLAIPELM' | 338 - 350 | 5E-03 | 0.6 | 2E+00 | 1 | 1.0 | 9 | 5E-07 | 0.5 | 1E-04 | 5 | 1.0 | 5 | 1E-06 | 1.0 | 1E-04 | 5 | 1.0 | 5 | 1E-04 | 1.0 | 1E-05 | 5 | 1.0 | 5 |
| 'MRVLVD' | 350 - 355 | 6E+04 | 0.8 | 4E-01 | 1 | 0.1 | 4 | 5E+04 | 1.0 | 7E-12 | 0 | 0.9 | 5 | 5E+04 | 1.0 | 3E-11 | 0 | 0.9 | 5 | 5E+04 | 1.0 | 1E-11 | 0 | 0.9 | 5 |
| 'MRVLVDL' | 350 - 356 | 6E-03 | 0.6 | 9E-01 | 1 | 1.0 | 5 | 2E-14 | 1.0 | 3E-11 | 3 | 1.0 | 3 | 4E-14 | 1.0 | 1E-09 | 3 | 1.0 | 3 | 4E-14 | 1.0 | 7E-10 | 3 | 1.0 | 3 |
| 'RVLVDL' | 351 - 356 | 6E-03 | 0.6 | 1E+00 | 1 | 1.0 | 4 | 4E-14 | 1.0 | 1E-09 | 3 | 1.0 | 2 | 4E-14 | 1.0 | 4E-10 | 3 | 1.0 | 2 | 4E-14 | 1.0 | 4E-10 | 3 | 1.0 | 2 |
| 'LERLDWDKA' | 356 - 364 | 2E+00 | 1.0 | 5E-02 | 7 | 0.7 | 1 | 3E-02 | 1.0 | 7E-04 | 3 | 0.5 | 5 | 4E-02 | 1.0 | 6E-04 | 3 | 0.5 | 5 | 3E-02 | 1.0 | 5E-04 | 3 | 0.6 | 5 |
| 'LERLDWDKAWE' | 356 - 366 | 4E-03 | 0.8 | 2E+00 | 1 | 1.0 | 9 | 2E-04 | 0.4 | 6E-03 | 6 | 1.0 | 4 | 4E-05 | 0.4 | 8E-03 | 6 | 1.0 | 4 | 2E-14 | 0.5 | 7E-03 | 6 | 1.0 | 4 |
| 'ERLDWDKA' | 357 - 364 | 2E+00 | 1.0 | 5E-02 | 6 | 0.7 | 1 | 8E-02 | 1.0 | 2E-03 | 3 | 0.5 | 4 | 9E-02 | 1.0 | 1E-03 | 3 | 0.6 | 4 | 6E-02 | 1.0 | 1E-03 | 3 | 0.7 | 4 |
| 'ERLDWDKAW' | 357 - 365 | 1E-02 | 0.7 | 2E+00 | 1 | 1.0 | 7 | 1E-03 | 0.5 | 1E-01 | 5 | 1.0 | 3 | 1E-03 | 0.5 | 1E-01 | 5 | 1.0 | 3 | 1E-03 | 0.5 | 8E-02 | 5 | 1.0 | 3 |
| 'ERLDWDKAWE' | 357 - 366 | 4E-03 | 0.7 | 2E+00 | 1 | 1.0 | 8 | 2E-04 | 0.4 | 2E-02 | 5 | 1.0 | 4 | 8E-12 | 0.5 | 2E-02 | 5 | 1.0 | 4 | 2E-12 | 0.5 | 1E-02 | 5 | 1.0 | 4 |
| 'VTVKTCA' | 367 - 373 | 3E+00 | 1.0 | 1E-01 | 5 | 0.5 | 1 | 3E-11 | 1.0 | 2E-14 | 3 | 1.0 | 3 | 3E-09 | 1.0 | 7E-14 | 3 | 1.0 | 3 | 1E-08 | 1.0 | 3E-13 | 3 | 1.0 | 3 |
| 'VKTCAYTNHTVLPEAL' | 369 - 384 | 2E+00 | 1.0 | 1E-02 | 13 | 0.6 | 1 | 1E-01 | 1.0 | 1E-03 | 6 | 0.5 | 8 | 1E-02 | 1.0 | 9E-06 | 6 | 0.5 | 8 | 1E-02 | 1.0 | 3E-07 | 7 | 0.5 | 7 |

| | | | | | | | | | | | | | | | | | | | | | | | | | |
|---|---|---|---|---|---|---|---|---|---|---|---|---|---|---|---|---|---|---|---|---|---|---|---|---|---|
| 'YTNHTVL' | 374 - 380 | 7E-03 | 0.4 | 1E+01 | 1 | 1.0 | 5 | 4E-03 | 0.9 | 2E-01 | 3 | 0.9 | 3 | 8E-09 | 1.0 | 1E-02 | 3 | 1.0 | 3 | 2E-08 | 1.0 | 1E-02 | 2 | 1.0 | 4 |
| 'YTNHTVLPEA' | 374 - 383 | 2E-02 | 0.4 | 3E+00 | 1 | 1.0 | 7 | 1E-02 | 0.7 | 3E-01 | 4 | 0.5 | 4 | 1E-07 | 0.7 | 2E-02 | 3 | 0.1 | 5 | 1E-03 | 0.7 | 2E-02 | 4 | 0.4 | 4 |
| 'YTNHTVLPEAL' | 374 - 384 | 1E-02 | 0.5 | 3E+00 | 1 | 1.0 | 8 | 3E-03 | 0.7 | 2E-01 | 5 | 0.8 | 4 | 6E-04 | 0.7 | 3E-02 | 5 | 1.0 | 4 | 8E-04 | 0.9 | 3E-02 | 5 | 1.0 | 4 |
| 'TNHTVL' | 375 - 380 | 7E-03 | 0.4 | 8E+00 | 1 | 1.0 | 4 | 4E-03 | 0.6 | 3E-01 | 3 | 0.6 | 2 | 6E-03 | 0.1 | 3E-07 | 3 | 0.9 | 2 | 5E-10 | 1.0 | 2E-02 | 3 | 1.0 | 2 |
| 'LERWPVHL' | 384 - 391 | 7E-02 | 0.5 | 3E+00 | 1 | 0.8 | 5 | 3E-04 | 0.5 | 2E-01 | 4 | 0.5 | 2 | 6E-04 | 0.5 | 1E-01 | 4 | 1.0 | 2 | 5E-04 | 0.5 | 9E-02 | 4 | 1.0 | 2 |
| 'ERWPVHL' | 385 - 391 | 2E+00 | 1.0 | 4E-02 | 4 | 0.4 | 1 | 1E-01 | 1.0 | 5E-04 | 2 | 0.5 | 3 | 1E-01 | 1.0 | 4E-04 | 2 | 0.6 | 3 | 9E-02 | 1.0 | 3E-04 | 2 | 0.7 | 3 |
| 'ERWPVHLLE' | 385 - 393 | 3E-03 | 0.5 | 1E+00 | 1 | 1.0 | 6 | 2E-14 | 0.6 | 1E-03 | 4 | 1.0 | 3 | 4E-14 | 0.6 | 9E-04 | 4 | 1.0 | 3 | 5E-13 | 0.6 | 1E-03 | 4 | 1.0 | 3 |
| 'LETLLPRHLQI' | 392 - 402 | 7E-03 | 0.6 | 2E+00 | 1 | 1.0 | 8 | 2E-04 | 1.0 | 7E-05 | 5 | 1.0 | 4 | 3E-04 | 1.0 | 8E-05 | 5 | 1.0 | 4 | 2E-04 | 1.0 | 1E-04 | 5 | 1.0 | 4 |
| 'LETLLPRHLQIIY' | 392 - 404 | 8E-03 | 0.6 | 1E+00 | 1 | 1.0 | 10 | 6E-07 | 0.7 | 3E-04 | 6 | 1.0 | 5 | 6E-07 | 0.9 | 4E-04 | 6 | 1.0 | 5 | 4E-06 | 1.0 | 3E-04 | 6 | 1.0 | 5 |
| 'LETLLPRHLQIIYE' | 392 - 405 | 5E-03 | 0.6 | 1E+00 | 1 | 1.0 | 11 | 1E-04 | 1.0 | 5E-05 | 7 | 1.0 | 5 | 2E-05 | 1.0 | 1E-05 | 7 | 1.0 | 5 | 1E-04 | 1.0 | 1E-04 | 6 | 1.0 | 6 |
| 'TLLPRHLQI' | 394 - 402 | 7E-03 | 0.5 | 2E+00 | 1 | 1.0 | 6 | 4E-11 | 0.1 | 7E-06 | 4 | 1.0 | 3 | 2E-08 | 0.1 | 6E-06 | 4 | 1.0 | 3 | 1E-11 | 0.2 | 6E-06 | 4 | 1.0 | 3 |
| 'TLLPRHLQIIY' | 394 - 404 | 8E-03 | 0.5 | 1E+00 | 1 | 1.0 | 8 | 2E-14 | 1.0 | 4E-11 | 5 | 1.0 | 4 | 3E-14 | 1.0 | 2E-10 | 5 | 1.0 | 4 | 2E-14 | 1.0 | 5E-11 | 5 | 1.0 | 4 |
| 'TLLPRHLQIIYE' | 394 - 405 | 5E-03 | 0.6 | 1E+00 | 1 | 1.0 | 9 | 2E-14 | 1.0 | 6E-12 | 5 | 1.0 | 5 | 2E-14 | 1.0 | 6E-12 | 5 | 1.0 | 5 | 3E-14 | 1.0 | 1E-09 | 5 | 1.0 | 5 |
| 'LPRHLQIIY' | 396 - 404 | 9E-03 | 0.6 | 1E+00 | 1 | 1.0 | 6 | 5E-12 | 0.5 | 2E-05 | 4 | 1.0 | 3 | 3E-12 | 0.5 | 3E-05 | 4 | 1.0 | 3 | 3E-14 | 1.0 | 1E-11 | 4 | 1.0 | 3 |
| 'LPRHLQIIYE' | 396 - 405 | 5E-03 | 0.6 | 1E+00 | 1 | 1.0 | 7 | 3E-05 | 0.4 | 1E-05 | 4 | 1.0 | 4 | 2E-06 | 0.5 | 3E-05 | 5 | 1.0 | 3 | 1E-04 | 1.0 | 2E-05 | 5 | 1.0 | 3 |
| 'PRHLQIIY' | 397 - 404 | 8E-03 | 0.5 | 2E+00 | 1 | 1.0 | 6 | 5E-13 | 1.0 | 9E-10 | 4 | 1.0 | 3 | 2E-14 | 1.0 | 3E-11 | 4 | 1.0 | 3 | 2E-14 | 1.0 | 6E-13 | 4 | 1.0 | 3 |
| 'PRHLQIIYE' | 397 - 405 | 5E-03 | 0.5 | 2E+00 | 1 | 1.0 | 7 | 2E-14 | 1.0 | 3E-14 | 4 | 1.0 | 4 | 2E-14 | 1.0 | 1E-12 | 3 | 1.0 | 4 | 2E-14 | 1.0 | 5E-14 | 3 | 1.0 | 4 |
| 'IYEINQRFLNRVAAAFPGDVDRL' | 403 - 425 | 2E+00 | 0.7 | 3E-02 | 20 | 0.7 | 1 | 2E-03 | 1.0 | 9E-13 | 10 | 0.2 | 11 | 2E-03 | 1.0 | 9E-13 | 10 | 0.2 | 11 | 1E-03 | 1.0 | 3E-14 | 10 | 0.2 | 11 |
| 'EINQRFLNRVAAAF' | 405 - 418 | 3E+00 | 1.0 | 1E-02 | 12 | 0.7 | 1 | 6E-05 | 1.0 | 2E-06 | 6 | 0.6 | 7 | 7E-05 | 1.0 | 3E-07 | 6 | 0.9 | 7 | 8E-05 | 1.0 | 3E-11 | 6 | 1.0 | 7 |
| 'EINQRFLNRVAAAFPGDVDRL' | 405 - 425 | 2E-02 | 0.7 | 2E+00 | 1 | 0.8 | 18 | 7E-13 | 0.2 | 2E-03 | 10 | 1.0 | 9 | 9E-13 | 0.2 | 2E-03 | 10 | 1.0 | 9 | 3E-14 | 0.2 | 2E-03 | 10 | 1.0 | 9 |
| 'EINQRFLNRVAAAFPGDVDRLRRMSL' | 405 - 430 | 2E+00 | 0.7 | 3E-02 | 23 | 0.7 | 1 | 2E-04 | 1.0 | 2E-13 | 11 | 0.2 | 13 | 2E-04 | 1.0 | 4E-12 | 11 | 0.2 | 13 | 3E-04 | 1.0 | 1E-13 | 11 | 0.2 | 13 |
| 'INQRFLNRVA' | 406 - 415 | 3E+00 | 1.0 | 3E-02 | 8 | 0.6 | 1 | 8E-06 | 1.0 | 7E-07 | 4 | 0.3 | 5 | 6E-05 | 1.0 | 6E-12 | 4 | 0.6 | 5 | 5E-05 | 1.0 | 2E-06 | 4 | 0.7 | 5 |
| 'INQRFLNRVAAAF' | 406 - 418 | 1E-02 | 0.7 | 3E+00 | 1 | 1.0 | 11 | 1E-06 | 0.9 | 6E-05 | 7 | 1.0 | 5 | 3E-05 | 1.0 | 9E-07 | 7 | 1.0 | 5 | 7E-09 | 1.0 | 1E-06 | 7 | 1.0 | 5 |
| 'INQRFLNRVAAAFPGDVDRL' | 406 - 425 | 2E+00 | 0.8 | 2E-02 | 17 | 0.7 | 1 | 2E-03 | 1.0 | 1E-13 | 8 | 0.2 | 10 | 2E-03 | 1.0 | 9E-14 | 8 | 0.2 | 10 | 2E-03 | 1.0 | 3E-14 | 8 | 0.2 | 10 |
| 'INQRFLNRVAAAFPGDVDRLRRMSL' | 406 - 430 | 2E+00 | 0.8 | 3E-02 | 22 | 0.7 | 1 | 2E-04 | 1.0 | 3E-13 | 11 | 0.2 | 12 | 2E-04 | 1.0 | 2E-12 | 11 | 0.2 | 12 | 3E-04 | 1.0 | 4E-13 | 11 | 0.2 | 12 |
| 'NQRFLNRVAAAFPGDVDRLRRMSL' | 407 - 430 | 2E+00 | 0.8 | 3E-02 | 21 | 0.7 | 1 | 2E-04 | 1.0 | 5E-13 | 10 | 0.2 | 12 | 1E-04 | 1.0 | 8E-14 | 10 | 0.2 | 12 | 2E-04 | 1.0 | 3E-14 | 10 | 0.2 | 12 |
| 'QRFLNRVAAAFPGDVDRL' | 408 - 425 | 2E+00 | 0.9 | 2E-02 | 15 | 0.7 | 1 | 6E-03 | 1.0 | 5E-14 | 7 | 0.2 | 9 | 7E-03 | 1.0 | 4E-14 | 7 | 0.2 | 9 | 6E-03 | 1.0 | 1E-13 | 7 | 0.3 | 9 |
| 'LNRVAAAFPGDVDRLRRMSL' | 411 - 430 | 2E+00 | 0.8 | 3E-02 | 17 | 0.7 | 1 | 1E-03 | 1.0 | 1E-12 | 8 | 0.2 | 10 | 1E-03 | 1.0 | 4E-14 | 8 | 0.2 | 10 | 1E-03 | 1.0 | 2E-13 | 8 | 0.2 | 10 |
| 'AAFPGDVDRL' | 416 - 425 | 2E-02 | 0.6 | 2E+00 | 1 | 0.9 | 7 | 1E-03 | 0.5 | 4E-01 | 5 | 1.0 | 3 | 2E-04 | 0.4 | 3E-01 | 4 | 0.4 | 4 | 5E-04 | 0.5 | 3E-01 | 5 | 0.6 | 3 |
| 'PGDVDRL' | 419 - 425 | 2E-02 | 0.5 | 2E+00 | 1 | 1.0 | 5 | 2E-04 | 0.5 | 6E-02 | 3 | 1.0 | 2 | 1E-04 | 0.6 | 6E-02 | 3 | 1.0 | 2 | 1E-04 | 0.6 | 6E-02 | 3 | 1.0 | 2 |
| 'VEEGAVKRINM' | 431 - 441 | 2E-02 | 0.7 | 2E+00 | 1 | 1.0 | 9 | 8E-09 | 0.6 | 4E-01 | 5 | 0.2 | 5 | 1E-13 | 0.6 | 5E-01 | 5 | 1.0 | 5 | 1E-06 | 0.6 | 5E-01 | 5 | 0.5 | 5 |
| 'VEEGAVKRINMAHL' | 431 - 444 | 3E+00 | 0.8 | 6E-02 | 12 | 0.7 | 1 | 2E-01 | 1.0 | 2E-14 | 6 | 0.4 | 7 | 3E-01 | 1.0 | 2E-14 | 6 | 0.5 | 7 | 2E-01 | 1.0 | 2E-14 | 6 | 0.5 | 7 |

| | | | | | | | | | | | | | | | | | | | | | | | | | |
|---|---|---|---|---|---|---|---|---|---|---|---|---|---|---|---|---|---|---|---|---|---|---|---|---|---|
| 'EEGAVKRINM' | 432 - 441 | 2E-02 | 0.6 | 2E+00 | 1 | 1.0 | 8 | 8E-07 | 0.2 | 3E-01 | 3 | 0.6 | 5 | 2E-14 | 0.6 | 7E-01 | 4 | 1.0 | 5 | 2E-07 | 0.5 | 6E-01 | 4 | 0.4 | 5 |
| 'CIAGSHAVNGVARIHS' | 445 - 460 | 3E+00 | 0.8 | 9E-02 | 14 | 0.6 | 1 | 7E-06 | 1.0 | 5E-04 | 7 | 0.2 | 8 | 2E-05 | 1.0 | 2E-09 | 7 | 0.3 | 8 | 1E-05 | 1.0 | 2E-06 | 7 | 0.3 | 8 |
| 'CIAGSHAVNGVARIHSE' | 445 - 461 | 9E-03 | 0.5 | 3E+00 | 1 | 1.0 | 15 | 3E-04 | 0.2 | 1E-05 | 9 | 1.0 | 7 | 2E-14 | 0.2 | 4E-07 | 7 | 1.0 | 9 | 1E-12 | 0.3 | 1E-05 | 9 | 1.0 | 7 |
| 'CIAGSHAVNGVARIHSEI' | 445 - 462 | 7E-03 | 0.6 | 3E+00 | 2 | 0.5 | 15 | 2E-04 | 0.2 | 3E-05 | 9 | 0.9 | 8 | 2E-12 | 0.2 | 4E-06 | 9 | 1.0 | 8 | 8E-08 | 0.3 | 1E-05 | 9 | 1.0 | 8 |
| 'CIAGSHAVNGVARIHSEIL' | 445 - 463 | 6E-03 | 0.5 | 3E+00 | 1 | 1.0 | 17 | 1E-04 | 0.2 | 4E-05 | 10 | 1.0 | 8 | 5E-12 | 0.2 | 5E-06 | 10 | 1.0 | 8 | 2E-07 | 0.2 | 5E-06 | 10 | 1.0 | 8 |
| 'CIAGSHAVNGVARIHSEILKKTIF' | 445 - 468 | 7E-03 | 0.6 | 2E+00 | 1 | 1.0 | 22 | 3E-04 | 0.2 | 2E-05 | 12 | 1.0 | 11 | 5E-08 | 0.2 | 3E-06 | 12 | 1.0 | 11 | 1E-05 | 0.3 | 2E-05 | 12 | 1.0 | 11 |
| 'CIAGSHAVNGVARIHSEILKKTIFKDF' | 445 - 471 | 2E+00 | 1.0 | 8E-03 | 25 | 0.6 | 1 | 2E-05 | 1.0 | 4E-04 | 12 | 0.2 | 14 | 2E-05 | 1.0 | 7E-06 | 13 | 0.2 | 13 | 5E-06 | 1.0 | 2E-04 | 13 | 0.2 | 13 |
| 'KKTIFKDF' | 464 - 471 | 7E-03 | 0.6 | 2E+00 | 1 | 1.0 | 6 | 1E-03 | 0.2 | 3E-05 | 4 | 1.0 | 3 | 1E-03 | 0.2 | 3E-05 | 4 | 1.0 | 3 | 1E-03 | 0.3 | 3E-05 | 4 | 1.0 | 3 |
| 'YELEPHKFQNKTNGITPRRW' | 472 - 491 | 3E+00 | 0.9 | 3E-02 | 16 | 0.6 | 1 | 2E-05 | 1.0 | 9E-07 | 8 | 0.2 | 9 | 1E-05 | 1.0 | 8E-12 | 8 | 0.2 | 9 | 1E-05 | 1.0 | 2E-06 | 10 | 0.3 | 7 |
| 'PHKFQNKTNGITPRRWLVL' | 476 - 494 | 7E-03 | 0.6 | 2E+00 | 1 | 1.0 | 16 | 1E-10 | 0.1 | 4E-06 | 8 | 1.0 | 8 | 6E-12 | 0.1 | 4E-06 | 8 | 1.0 | 8 | 2E-08 | 0.1 | 6E-06 | 8 | 1.0 | 8 |
| 'LVLCNPGL' | 492 - 499 | 1E-02 | 0.5 | 3E+00 | 1 | 1.0 | 5 | 2E-04 | 0.2 | 2E-05 | 3 | 1.0 | 3 | 3E-14 | 0.2 | 3E-07 | 3 | 1.0 | 3 | 3E-14 | 0.3 | 3E-05 | 3 | 1.0 | 3 |
| 'CNPGLAE' | 495 - 501 | 5E-03 | 0.7 | 2E+00 | 1 | 1.0 | 4 | 1E-03 | 0.2 | 3E-05 | 3 | 1.0 | 2 | 2E-04 | 0.3 | 1E-05 | 3 | 1.0 | 2 | 1E-04 | 0.5 | 5E-05 | 3 | 1.0 | 2 |
| 'GLAEII' | 498 - 503 | 3E-03 | 0.5 | 1E+00 | 1 | 1.0 | 4 | 2E-14 | 1.0 | 5E-12 | 3 | 1.0 | 2 | 2E-14 | 1.0 | 4E-13 | 3 | 1.0 | 2 | 2E-14 | 1.0 | 4E-10 | 3 | 1.0 | 2 |
| 'AEIIAE' | 500 - 505 | 5E-03 | 0.6 | 7E-01 | 1 | 1.0 | 4 | 1E-10 | 0.3 | 7E-06 | 3 | 1.0 | 2 | 1E-09 | 0.8 | 1E-04 | 3 | 1.0 | 2 | 2E-05 | 1.0 | 3E-04 | 3 | 1.0 | 2 |
| 'IAERIGEE' | 503 - 510 | 3E-03 | 0.7 | 2E+00 | 1 | 1.0 | 6 | 3E-05 | 0.4 | 2E-03 | 3 | 0.6 | 4 | 2E-14 | 0.5 | 7E-03 | 4 | 1.0 | 3 | 2E-14 | 0.6 | 7E-03 | 4 | 1.0 | 3 |
| 'EYISDL' | 510 - 515 | 7E-03 | 0.5 | 3E+00 | 1 | 1.0 | 4 | 2E-06 | 0.5 | 2E-03 | 2 | 0.7 | 3 | 6E-07 | 0.6 | 5E-03 | 3 | 1.0 | 2 | 2E-14 | 0.7 | 5E-03 | 3 | 1.0 | 2 |
| 'YISDLDQ' | 511 - 517 | 8E-03 | 0.5 | 3E+00 | 1 | 1.0 | 5 | 8E-03 | 0.1 | 1E-08 | 3 | 0.8 | 3 | 5E-03 | 0.1 | 6E-09 | 3 | 0.8 | 3 | 4E-03 | 0.2 | 8E-07 | 3 | 0.9 | 3 |
| 'DQLRKLLS' | 516 - 523 | 8E-03 | 0.6 | 1E+00 | 1 | 1.0 | 6 | 2E-14 | 0.3 | 2E-03 | 4 | 1.0 | 3 | 2E-14 | 0.3 | 2E-03 | 4 | 1.0 | 3 | 2E-14 | 0.3 | 2E-03 | 4 | 1.0 | 3 |
| 'YVDDEA' | 524 - 529 | 3E-02 | 0.7 | 3E+00 | 1 | 1.0 | 4 | 6E-06 | 0.9 | 3E-02 | 2 | 0.5 | 3 | 2E-03 | 1.0 | 5E-02 | 3 | 1.0 | 2 | 6E-04 | 1.0 | 3E-02 | 2 | 1.0 | 3 |
| 'YVDDEAF' | 524 - 530 | 1E-02 | 0.8 | 2E+00 | 1 | 1.0 | 5 | 4E-05 | 0.7 | 2E-02 | 3 | 0.8 | 3 | 6E-07 | 0.7 | 3E-02 | 3 | 0.3 | 3 | 1E-06 | 0.7 | 2E-02 | 3 | 0.9 | 3 |
| 'FIRDVAKVKQENKLKF' | 530 - 545 | 2E-02 | 0.7 | 2E+00 | 1 | 0.9 | 14 | 3E-07 | 0.9 | 3E-05 | 8 | 1.0 | 7 | 1E-06 | 0.9 | 2E-05 | 8 | 1.0 | 7 | 3E-14 | 1.0 | 8E-09 | 8 | 1.0 | 7 |
| 'FIRDVAKVKQENKLKFAAY' | 530 - 548 | 2E-02 | 0.7 | 2E+00 | 1 | 0.8 | 17 | 7E-07 | 0.7 | 4E-05 | 10 | 1.0 | 8 | 4E-11 | 0.8 | 5E-05 | 10 | 1.0 | 8 | 8E-07 | 0.8 | 2E-05 | 10 | 1.0 | 8 |
| 'IRDVAKVKQENKLKF' | 531 - 545 | 2E+00 | 0.9 | 2E-02 | 13 | 0.7 | 1 | 3E-05 | 1.0 | 2E-07 | 6 | 0.9 | 8 | 4E-09 | 1.0 | 2E-12 | 6 | 1.0 | 8 | 5E-09 | 1.0 | 2E-14 | 6 | 1.0 | 8 |
| 'IRDVAKVKQENKLKFAAY' | 531 - 548 | 2E-02 | 0.7 | 2E+00 | 1 | 0.8 | 16 | 1E-06 | 0.4 | 5E-06 | 9 | 1.0 | 8 | 6E-07 | 0.7 | 4E-05 | 9 | 1.0 | 8 | 3E-07 | 0.7 | 3E-05 | 9 | 1.0 | 8 |
| 'YLEREYKVHINPNSL' | 548 - 562 | 3E+00 | 1.0 | 2E-02 | 12 | 0.7 | 1 | 5E-04 | 1.0 | 8E-04 | 6 | 0.3 | 7 | 9E-07 | 0.5 | 2E-03 | 7 | 0.2 | 6 | 5E-05 | 0.6 | 2E-03 | 7 | 0.3 | 6 |
| 'LEREYKVHINPNSL' | 549 - 562 | 3E+00 | 1.0 | 2E-02 | 11 | 0.7 | 1 | 1E-03 | 1.0 | 1E-03 | 5 | 0.3 | 7 | 3E-03 | 0.6 | 1E-07 | 8 | 0.4 | 4 | 4E-03 | 1.0 | 8E-04 | 5 | 0.4 | 7 |
| 'LEREYKVHINPNSLF' | 549 - 563 | 7E-03 | 0.7 | 3E+00 | 1 | 1.0 | 12 | 4E-03 | 0.2 | 9E-06 | 4 | 0.8 | 9 | 4E-03 | 0.2 | 3E-05 | 4 | 0.7 | 9 | 1E-03 | 0.3 | 2E-03 | 7 | 0.9 | 6 |
| 'EREYKVHINPNSL' | 550 - 562 | 3E+00 | 1.0 | 2E-02 | 10 | 0.6 | 1 | 1E-03 | 1.0 | 1E-03 | 5 | 0.3 | 6 | 8E-07 | 0.6 | 5E-03 | 6 | 0.2 | 5 | 3E-03 | 0.9 | 8E-04 | 5 | 0.4 | 6 |
| 'EREYKVHINPNSLF' | 550 - 563 | 7E-03 | 0.6 | 3E+00 | 1 | 1.0 | 11 | 8E-03 | 0.2 | 8E-06 | 3 | 1.0 | 9 | 1E-02 | 0.2 | 3E-07 | 4 | 0.8 | 8 | 9E-03 | 0.3 | 3E-05 | 3 | 1.0 | 9 |
| 'YKVHINPNSL' | 553 - 562 | 2E-02 | 0.6 | 4E+00 | 1 | 1.0 | 7 | 2E-03 | 0.4 | 6E-04 | 5 | 1.0 | 3 | 2E-04 | 0.7 | 9E-03 | 5 | 1.0 | 3 | 3E-04 | 0.8 | 8E-03 | 5 | 1.0 | 3 |
| 'FDVQVKRIHE' | 563 - 572 | 2E+00 | 1.0 | 2E-02 | 8 | 0.6 | 1 | 6E-03 | 1.0 | 4E-14 | 4 | 0.3 | 5 | 1E-06 | 0.6 | 2E-04 | 4 | 0.2 | 5 | 7E-05 | 1.0 | 5E-04 | 4 | 0.3 | 5 |
| 'FDVQVKRIHEYKRQLL' | 563 - 578 | 2E+00 | 1.0 | 6E-03 | 14 | 0.6 | 1 | 1E-03 | 1.0 | 2E-13 | 7 | 0.3 | 8 | 7E-05 | 1.0 | 2E-04 | 7 | 0.3 | 8 | 1E-04 | 1.0 | 3E-04 | 7 | 0.3 | 8 |

| | | | | | | | | | | | | | | | | | | | | | | | | | |
|---|---|---|---|---|---|---|---|---|---|---|---|---|---|---|---|---|---|---|---|---|---|---|---|---|---|
| 'FDVQVKRIHEYKRQLLNC' | 563 - 580 | 2E+00 | 0.9 | 3E-02 | 16 | 0.6 | 1 | 5E-04 | 1.0 | 2E-12 | 8 | 0.3 | 9 | 4E-05 | 1.0 | 2E-04 | 8 | 0.3 | 9 | 1E-04 | 1.0 | 2E-04 | 8 | 0.3 | 9 |
| 'FDVQVKRIHEYKRQLLNCL' | 563 - 581 | 2E+00 | 0.8 | 5E-02 | 17 | 0.6 | 1 | 6E-04 | 1.0 | 2E-12 | 8 | 0.3 | 10 | 4E-05 | 1.0 | 1E-04 | 8 | 0.2 | 10 | 9E-05 | 1.0 | 2E-04 | 8 | 0.3 | 10 |
| 'DVQVKRIHEYKRQLLNC' | 564 - 580 | 3E-02 | 0.7 | 2E+00 | 1 | 0.9 | 15 | 8E-13 | 0.3 | 8E-04 | 9 | 1.0 | 7 | 2E-04 | 0.3 | 7E-05 | 9 | 1.0 | 7 | 2E-04 | 0.4 | 1E-04 | 9 | 1.0 | 7 |
| 'LHVITL' | 581 - 586 | 2E-02 | 0.5 | 1E+00 | 1 | 1.0 | 4 | 4E-14 | 1.0 | 2E-08 | 3 | 1.0 | 2 | 4E-12 | 1.0 | 6E-09 | 3 | 1.0 | 2 | 1E-13 | 1.0 | 3E-10 | 3 | 1.0 | 2 |
| 'YNRIKKEPNKFVVPRTVM' | 587 - 604 | 6E-03 | 0.6 | 2E+00 | 1 | 1.0 | 14 | 2E-14 | 0.3 | 2E-02 | 8 | 1.0 | 7 | 2E-14 | 0.3 | 2E-02 | 8 | 1.0 | 7 | 3E-14 | 0.3 | 2E-02 | 8 | 1.0 | 7 |
| 'IGGKAAPGYHMAKM' | 605 - 618 | 1E-02 | 0.8 | 3E+00 | 1 | 1.0 | 11 | 1E-03 | 0.5 | 4E-01 | 7 | 1.0 | 5 | 3E-04 | 0.5 | 2E-01 | 7 | 1.0 | 5 | 3E-04 | 0.5 | 1E-01 | 7 | 1.0 | 5 |
| 'GGKAAPGYHMAKM' | 606 - 618 | 1E-02 | 0.8 | 3E+00 | 1 | 1.0 | 10 | 9E-04 | 0.5 | 3E-01 | 6 | 1.0 | 5 | 7E-05 | 0.5 | 1E-01 | 6 | 1.0 | 5 | 4E-05 | 0.5 | 9E-02 | 6 | 1.0 | 5 |
| 'IIKLIT' | 619 - 624 | 3E-03 | 0.6 | 1E+00 | 1 | 1.0 | 4 | 3E-14 | 1.0 | 3E-08 | 3 | 1.0 | 2 | 2E-14 | 1.0 | 2E-10 | 3 | 1.0 | 2 | 2E-14 | 1.0 | 1E-11 | 3 | 1.0 | 2 |
| 'IIKLITA' | 619 - 625 | 1E+00 | 0.9 | 7E-02 | 5 | 0.7 | 1 | 4E-06 | 1.0 | 4E-09 | 2 | 0.2 | 4 | 1E-05 | 1.0 | 7E-06 | 2 | 0.3 | 4 | 2E-04 | 1.0 | 7E-07 | 2 | 1.0 | 4 |
| 'IIKLITAIGDVVNHDPVVGDRL' | 619 - 640 | 1E+00 | 1.0 | 2E-02 | 19 | 0.6 | 1 | 2E-03 | 1.0 | 1E-03 | 10 | 0.3 | 10 | 4E-04 | 0.7 | 2E-03 | 14 | 0.3 | 6 | 3E-03 | 1.0 | 1E-03 | 10 | 0.3 | 10 |
| 'ITAIGD' | 623 - 628 | 4E+00 | 0.9 | 5E-01 | 3 | 0.5 | 2 | 6E-06 | 1.0 | 3E-14 | 1 | 0.1 | 4 | 3E-06 | 1.0 | 3E-14 | 1 | 0.1 | 4 | 7E-06 | 1.0 | 2E-08 | 1 | 0.1 | 4 |
| 'ITAIGDVVNHDPVVGDRL' | 623 - 640 | 1E+00 | 1.0 | 2E-02 | 15 | 0.5 | 1 | 3E-08 | 0.8 | 6E-03 | 10 | 0.1 | 6 | 4E-08 | 0.7 | 6E-03 | 10 | 0.1 | 6 | 4E-03 | 1.0 | 1E-03 | 8 | 0.3 | 8 |
| 'ITAIGDVVNHDPVVGDRLRV' | 623 - 642 | 1E+00 | 1.0 | 1E-02 | 17 | 0.5 | 1 | 8E-06 | 0.8 | 4E-03 | 12 | 0.2 | 6 | 1E-06 | 0.7 | 3E-03 | 10 | 0.1 | 8 | 2E-05 | 0.9 | 6E-03 | 14 | 0.2 | 4 |
| 'IGDVVNHDPVVGDRL' | 626 - 640 | 2E+00 | 1.0 | 1E-02 | 12 | 0.5 | 1 | 7E-04 | 1.0 | 2E-03 | 6 | 0.2 | 7 | 3E-03 | 0.9 | 2E-03 | 6 | 0.3 | 7 | 4E-03 | 1.0 | 2E-03 | 6 | 0.3 | 7 |
| 'GDVVNHDPVVGDRL' | 627 - 640 | 1E+00 | 1.0 | 1E-02 | 11 | 0.5 | 1 | 1E-03 | 1.0 | 1E-03 | 6 | 0.2 | 6 | 2E-03 | 1.0 | 1E-03 | 6 | 0.2 | 6 | 8E-04 | 1.0 | 1E-03 | 6 | 0.2 | 6 |
| 'DVVNHDPVVGDRL' | 628 - 640 | 2E-02 | 0.5 | 2E+00 | 1 | 1.0 | 10 | 1E-03 | 0.2 | 2E-03 | 6 | 0.9 | 5 | 1E-02 | 0.2 | 7E-05 | 3 | 1.0 | 8 | 2E-02 | 0.2 | 4E-06 | 3 | 1.0 | 8 |
| 'LENYRVSL' | 645 - 652 | 2E-02 | 0.7 | 3E+00 | 1 | 1.0 | 6 | 3E-04 | 0.1 | 4E-08 | 4 | 0.4 | 3 | 5E-06 | 0.2 | 7E-06 | 4 | 0.3 | 3 | 3E-14 | 0.3 | 2E-03 | 4 | 1.0 | 3 |
| 'YRVSLAEKVIPAAD' | 648 - 661 | 1E+00 | 1.0 | 1E+00 | 9 | 0.6 | 3 | 3E-06 | 0.1 | 1E-07 | 10 | 0.4 | 2 | 2E-05 | 0.1 | 5E-05 | 10 | 0.5 | 2 | 3E-05 | 0.1 | 2E-04 | 10 | 0.6 | 2 |
| 'AEKVIPAAD' | 653 - 661 | 1E+00 | 0.0 | 6E-01 | 0 | 7.0 | 0 | 3E-14 | 0.1 | 2E-08 | 5 | 1.0 | 2 | 3E-14 | 0.1 | 7E-09 | 5 | 1.0 | 2 | 3E-14 | 0.2 | 1E-07 | 5 | 1.0 | 2 |
| 'AEKVIPAADL' | 653 - 662 | 7E-03 | 0.6 | 1E+00 | 1 | 1.0 | 7 | 8E-13 | 0.3 | 1E-05 | 5 | 1.0 | 3 | 2E-12 | 0.7 | 3E-05 | 5 | 1.0 | 3 | 2E-12 | 0.4 | 9E-06 | 5 | 1.0 | 3 |
| 'AEKVIPAADLSE' | 653 - 664 | 9E-03 | 0.6 | 1E+00 | 1 | 1.0 | 9 | 2E-08 | 0.3 | 9E-06 | 6 | 1.0 | 4 | 1E-12 | 0.2 | 4E-06 | 6 | 1.0 | 4 | 7E-12 | 0.6 | 5E-05 | 6 | 1.0 | 4 |
| 'KVIPAAD' | 655 - 661 | 3E-01 | 0.6 | 2E+00 | 2 | 1.0 | 3 | 2E-08 | 0.2 | 6E-06 | 3 | 1.0 | 2 | 2E-08 | 0.2 | 8E-06 | 3 | 1.0 | 2 | 4E-08 | 0.3 | 2E-05 | 3 | 1.0 | 2 |
| 'LSEQISTAGTEASGTGNMKF' | 662 - 681 | 3E+00 | 0.9 | 2E-02 | 18 | 0.8 | 1 | 3E-03 | 1.0 | 5E-05 | 9 | 0.2 | 10 | 1E-06 | 0.6 | 1E-04 | 9 | 0.1 | 10 | 6E-08 | 0.4 | 1E-04 | 11 | 0.1 | 8 |
| 'SEQISTAGTEASGTGNMKF' | 663 - 681 | 3E+00 | 0.9 | 2E-02 | 17 | 0.9 | 1 | 2E-02 | 1.0 | 2E-14 | 8 | 0.5 | 10 | 2E-03 | 1.0 | 2E-14 | 8 | 0.5 | 10 | 2E-03 | 1.0 | 2E-14 | 8 | 0.5 | 10 |
| 'QISTAGTEASGTGNMKF' | 665 - 681 | 3E+00 | 0.9 | 2E-02 | 15 | 0.8 | 1 | 2E-02 | 1.0 | 2E-14 | 7 | 0.5 | 9 | 3E-03 | 1.0 | 2E-14 | 7 | 0.5 | 9 | 2E-03 | 1.0 | 2E-14 | 7 | 0.5 | 9 |
| 'ISTAGTEASGTGNMKF' | 666 - 681 | 3E+00 | 1.0 | 1E-02 | 14 | 0.9 | 1 | 3E-02 | 1.0 | 2E-14 | 7 | 0.5 | 8 | 2E-03 | 1.0 | 4E-14 | 7 | 0.5 | 8 | 3E-03 | 1.0 | 2E-14 | 7 | 0.5 | 8 |
| 'MLNGAL' | 682 - 687 | 1E-02 | 0.8 | 6E+00 | 1 | 1.0 | 4 | 7E-13 | 0.4 | 8E-06 | 3 | 1.0 | 2 | 1E-12 | 1.0 | 5E-07 | 3 | 1.0 | 2 | 2E-14 | 1.0 | 1E-08 | 3 | 1.0 | 2 |
| 'MLNGALTIGTM' | 682 - 692 | 1E-02 | 0.7 | 2E+00 | 1 | 1.0 | 9 | 5E-13 | 0.2 | 3E-06 | 6 | 1.0 | 4 | 8E-08 | 0.3 | 1E-05 | 6 | 1.0 | 4 | 4E-14 | 0.2 | 1E-08 | 5 | 1.0 | 5 |
| 'MLNGALTIGTMDGAN' | 682 - 696 | 2E+00 | 1.0 | 1E-02 | 13 | 0.7 | 1 | 2E-05 | 1.0 | 1E-04 | 6 | 0.2 | 8 | 4E-06 | 1.0 | 6E-11 | 6 | 0.2 | 8 | 4E-06 | 1.0 | 6E-08 | 6 | 0.2 | 8 |
| 'ALTIGTMDGAN' | 686 - 696 | 1E-02 | 0.7 | 2E+00 | 1 | 1.0 | 9 | 7E-04 | 0.1 | 4E-05 | 6 | 0.9 | 4 | 1E-13 | 0.1 | 2E-08 | 5 | 0.8 | 5 | 5E-06 | 0.1 | 7E-06 | 6 | 1.0 | 4 |
| 'TIGTMDGANVE' | 688 - 698 | 3E-03 | 0.7 | 3E+00 | 1 | 1.0 | 9 | 1E-03 | 0.1 | 6E-05 | 6 | 1.0 | 4 | 1E-06 | 0.1 | 6E-06 | 6 | 1.0 | 4 | 9E-11 | 0.1 | 4E-06 | 6 | 1.0 | 4 |
| 'IGTMDGAN' | 689 - 696 | 1E-02 | 0.8 | 4E+00 | 1 | 1.0 | 6 | 1E-03 | 0.1 | 7E-05 | 4 | 0.7 | 3 | 1E-04 | 0.1 | 7E-06 | 4 | 1.0 | 3 | 3E-04 | 0.1 | 1E-05 | 4 | 1.0 | 3 |

| | | | | | | | | | | | | | | | | | | | | | | | | | |
|---|---|---|---|---|---|---|---|---|---|---|---|---|---|---|---|---|---|---|---|---|---|---|---|---|---|
| 'VEMAEE' | 697 - 702 | 3E-03 | 0.7 | 3E+00 | 1 | 1.0 | 4 | 1E-12 | 0.2 | 4E-06 | 3 | 1.0 | 2 | 2E-13 | 0.2 | 4E-06 | 3 | 1.0 | 2 | 4E-14 | 1.0 | 2E-10 | 3 | 1.0 | 2 |
| 'MAEEAGEENF' | 699 - 708 | 2E+00 | 1.0 | 2E-02 | 8 | 0.8 | 1 | 7E-08 | 0.3 | 8E-05 | 4 | 0.1 | 5 | 2E-06 | 0.4 | 9E-05 | 4 | 0.1 | 5 | 4E-05 | 0.4 | 5E-05 | 4 | 0.1 | 5 |
| 'AEEAGEENF' | 700 - 708 | 2E-02 | 0.7 | 3E+00 | 1 | 1.0 | 7 | 3E-04 | 0.1 | 1E-06 | 3 | 0.3 | 5 | 3E-04 | 0.1 | 5E-08 | 4 | 0.4 | 4 | 5E-04 | 0.1 | 4E-08 | 4 | 0.4 | 4 |
| 'EAGEENF' | 702 - 708 | 2E-02 | 0.7 | 3E+00 | 1 | 1.0 | 5 | 1E-02 | 0.1 | 3E-08 | 2 | 0.4 | 4 | 2E-03 | 0.1 | 2E-08 | 3 | 0.4 | 3 | 4E-03 | 0.1 | 5E-09 | 3 | 0.4 | 3 |
| 'AGEENF' | 703 - 708 | 2E-02 | 0.6 | 4E+00 | 1 | 1.0 | 4 | 3E-07 | 0.5 | 1E-02 | 3 | 0.1 | 2 | 1E-07 | 0.4 | 1E-02 | 3 | 0.1 | 2 | 2E-02 | 0.1 | 2E-08 | 2 | 0.6 | 3 |
| 'FIFGMRV' | 709 - 715 | 1E-02 | 0.7 | 3E+00 | 1 | 1.0 | 5 | 2E-03 | 0.2 | 4E-05 | 4 | 1.0 | 2 | 4E-14 | 0.5 | 7E-03 | 4 | 1.0 | 2 | 6E-04 | 0.8 | 2E-03 | 4 | 1.0 | 2 |
| 'FIFGMRVED' | 709 - 717 | 2E+00 | 0.7 | 4E-01 | 6 | 0.7 | 2 | 1E-04 | 0.8 | 1E-03 | 3 | 0.2 | 5 | 1E-07 | 0.6 | 2E-03 | 3 | 0.2 | 5 | 1E-08 | 0.7 | 5E-03 | 5 | 0.2 | 3 |
| 'MRVEDVDRLDQRGYNAQE' | 713 - 730 | 8E-03 | 0.7 | 2E+00 | 1 | 0.9 | 16 | 1E-06 | 0.5 | 1E-02 | 7 | 0.2 | 10 | 1E-03 | 0.4 | 1E-01 | 9 | 1.0 | 8 | 1E-03 | 0.5 | 1E-01 | 9 | 1.0 | 8 |
| 'MRVEDVDRLDQRGYNAQEY' | 713 - 731 | 1E-02 | 0.7 | 2E+00 | 1 | 0.9 | 17 | 6E-04 | 0.3 | 1E-02 | 7 | 1.0 | 11 | 9E-04 | 0.4 | 1E-01 | 10 | 1.0 | 8 | 5E-04 | 0.5 | 1E-01 | 10 | 1.0 | 8 |
| 'RVEDVDRL' | 714 - 721 | 2E-02 | 0.6 | 2E+00 | 1 | 0.9 | 6 | 7E-04 | 0.2 | 3E-05 | 4 | 0.8 | 3 | 5E-04 | 0.1 | 2E-08 | 4 | 0.5 | 3 | 8E-04 | 0.1 | 4E-08 | 4 | 0.6 | 3 |
| 'RVEDVDRLDQRGYNAQE' | 714 - 730 | 8E-03 | 0.7 | 2E+00 | 1 | 0.9 | 15 | 3E-04 | 0.4 | 3E-02 | 7 | 1.0 | 9 | 8E-04 | 0.5 | 2E-01 | 8 | 1.0 | 8 | 6E-04 | 0.5 | 1E-01 | 9 | 0.7 | 7 |
| 'VDRLDQRGYNAQE' | 718 - 730 | 7E-03 | 0.8 | 3E+00 | 1 | 1.0 | 11 | 8E-04 | 0.4 | 7E-02 | 6 | 1.0 | 5 | 1E-03 | 0.6 | 3E-01 | 6 | 0.9 | 6 | 1E-03 | 0.6 | 2E-01 | 7 | 1.0 | 5 |
| 'VDRLDQRGYNAQEY' | 718 - 731 | 3E+00 | 1.0 | 1E-02 | 12 | 0.8 | 1 | 6E-02 | 1.0 | 4E-04 | 6 | 0.4 | 7 | 2E-01 | 1.0 | 9E-04 | 6 | 0.5 | 7 | 2E-01 | 1.0 | 1E-03 | 6 | 0.5 | 7 |
| 'DQRGYNAQE' | 722 - 730 | 7E-03 | 0.8 | 5E+00 | 1 | 1.0 | 7 | 2E-03 | 0.6 | 3E-01 | 4 | 0.9 | 4 | 2E-03 | 0.7 | 3E-01 | 3 | 0.6 | 5 | 2E-03 | 0.7 | 3E-01 | 3 | 0.6 | 5 |
| 'DQRGYNAQEY' | 722 - 731 | 1E-02 | 0.8 | 4E+00 | 1 | 1.0 | 8 | 6E-04 | 0.5 | 2E-01 | 5 | 0.6 | 4 | 1E-03 | 0.7 | 3E-01 | 5 | 0.3 | 4 | 1E-03 | 0.6 | 3E-01 | 4 | 0.5 | 5 |
| 'YYDRIPEL' | 731 - 738 | 7E-03 | 0.7 | 2E+00 | 1 | 1.0 | 5 | 2E-13 | 0.2 | 2E-03 | 3 | 1.0 | 3 | 2E-14 | 0.3 | 1E-03 | 3 | 1.0 | 3 | 2E-14 | 0.3 | 1E-03 | 3 | 1.0 | 3 |
| 'YYDRIPELRQ' | 731 - 740 | 2E-02 | 0.7 | 1E+00 | 1 | 1.0 | 7 | 1E-12 | 0.2 | 2E-03 | 4 | 1.0 | 4 | 2E-12 | 0.2 | 2E-03 | 4 | 1.0 | 4 | 4E-14 | 0.2 | 2E-03 | 4 | 1.0 | 4 |
| 'YDRIPEL' | 732 - 738 | 7E-03 | 0.7 | 2E+00 | 1 | 1.0 | 4 | 2E-14 | 0.3 | 5E-02 | 3 | 1.0 | 2 | 2E-14 | 0.3 | 3E-02 | 3 | 1.0 | 2 | 2E-14 | 0.3 | 2E-02 | 3 | 1.0 | 2 |
| 'RQIIEQL' | 739 - 745 | 2E-02 | 0.6 | 1E+00 | 1 | 1.0 | 5 | 3E-06 | 0.5 | 3E-05 | 3 | 1.0 | 3 | 2E-10 | 1.0 | 2E-05 | 3 | 1.0 | 3 | 9E-14 | 1.0 | 7E-09 | 3 | 1.0 | 3 |
| 'RQIIEQLSSG' | 739 - 748 | 2E+00 | 1.0 | 5E-02 | 8 | 0.6 | 1 | 1E-03 | 1.0 | 4E-07 | 4 | 0.7 | 5 | 2E-04 | 1.0 | 2E-04 | 4 | 0.5 | 5 | 3E-04 | 1.0 | 3E-04 | 4 | 0.5 | 5 |
| 'LSSGFFSPKQPDL' | 745 - 757 | 7E-03 | 0.7 | 3E+00 | 1 | 1.0 | 9 | 4E-04 | 0.4 | 3E-02 | 5 | 1.0 | 5 | 2E-04 | 0.4 | 6E-02 | 5 | 1.0 | 5 | 1E-04 | 0.5 | 5E-02 | 5 | 1.0 | 5 |
| 'SSGFFSPKQPDL' | 746 - 757 | 7E-03 | 0.7 | 3E+00 | 1 | 1.0 | 8 | 3E-04 | 0.4 | 3E-02 | 5 | 1.0 | 4 | 2E-04 | 0.4 | 7E-02 | 5 | 1.0 | 4 | 2E-04 | 0.5 | 6E-02 | 5 | 1.0 | 4 |
| 'SSGFFSPKQPDLF' | 746 - 758 | 7E-03 | 0.6 | 2E+00 | 1 | 1.0 | 9 | 5E-04 | 0.3 | 3E-02 | 6 | 1.0 | 4 | 4E-04 | 0.4 | 8E-02 | 6 | 1.0 | 4 | 3E-04 | 0.3 | 6E-02 | 6 | 1.0 | 4 |
| 'FFSPKQPDL' | 749 - 757 | 7E-03 | 0.7 | 3E+00 | 1 | 1.0 | 5 | 1E-03 | 0.6 | 3E-01 | 3 | 1.0 | 3 | 2E-04 | 0.6 | 3E-01 | 3 | 0.5 | 3 | 5E-04 | 0.6 | 3E-01 | 3 | 0.5 | 3 |
| 'FSPKQPDL' | 750 - 757 | 7E-03 | 0.7 | 3E+00 | 1 | 1.0 | 4 | 2E-03 | 0.6 | 6E-01 | 3 | 0.6 | 2 | 4E-04 | 0.6 | 4E-01 | 2 | 0.5 | 3 | 7E-04 | 0.6 | 4E-01 | 2 | 0.5 | 3 |
| 'SPKQPDL' | 751 - 757 | 7E-03 | 0.9 | 2E+00 | 1 | 1.0 | 3 | 1E-03 | 0.5 | 5E-01 | 3 | 0.9 | 1 | 9E-05 | 0.5 | 3E-01 | 2 | 0.5 | 2 | 3E-04 | 0.6 | 3E-01 | 2 | 0.5 | 2 |
| 'PKQPDL' | 752 - 757 | 7E-03 | 0.6 | 5E+00 | 1 | 1.0 | 3 | 1E-03 | 0.5 | 3E-01 | 1 | 1.0 | 2 | 3E-04 | 0.5 | 3E-01 | 1 | 0.5 | 2 | 5E-04 | 0.5 | 3E-01 | 1 | 0.5 | 2 |
| 'FKDIVN' | 758 - 763 | 7E-03 | 0.5 | 9E-01 | 1 | 1.0 | 4 | 4E-03 | 0.1 | 4E-06 | 3 | 1.0 | 2 | 2E-03 | 0.2 | 4E-05 | 3 | 1.0 | 2 | 2E-03 | 0.2 | 1E-05 | 3 | 1.0 | 2 |
| 'FKDIVNM' | 758 - 764 | 2E-02 | 0.4 | 1E+00 | 1 | 1.0 | 5 | 1E-03 | 0.2 | 5E-05 | 3 | 1.0 | 3 | 1E-03 | 0.2 | 1E-05 | 3 | 1.0 | 3 | 1E-03 | 0.3 | 5E-05 | 3 | 1.0 | 3 |
| 'KDIVNM' | 759 - 764 | 2E-02 | 0.5 | 8E-01 | 1 | 1.0 | 4 | 1E-03 | 0.2 | 7E-06 | 3 | 1.0 | 2 | 9E-04 | 0.5 | 5E-04 | 3 | 1.0 | 2 | 5E-04 | 0.8 | 1E-03 | 3 | 1.0 | 2 |
| 'LMHHDRF' | 765 - 771 | 2E-02 | 0.5 | 1E+01 | 1 | 1.0 | 5 | 5E-04 | 1.0 | 8E-04 | 4 | 1.0 | 2 | 3E-04 | 1.0 | 3E-04 | 4 | 1.0 | 2 | 3E-04 | 1.0 | 3E-04 | 4 | 1.0 | 2 |
| 'LMHHDRFKVF' | 765 - 774 | 8E-03 | 0.5 | 5E+00 | 1 | 1.0 | 8 | 3E-04 | 1.0 | 1E-04 | 5 | 1.0 | 4 | 3E-05 | 1.0 | 7E-05 | 5 | 1.0 | 4 | 2E-14 | 1.0 | 1E-05 | 5 | 1.0 | 4 |

| | | | | | | | | | | | | | | | | | | | | | | | | | |
|---|---|---|---|---|---|---|---|---|---|---|---|---|---|---|---|---|---|---|---|---|---|---|---|---|---|
| 'LMHHDRFKVFAD' | 765 - 776 | 3E+00 | 1.0 | 3E-01 | 10 | 0.5 | 1 | 5E-05 | 1.0 | 2E-08 | 5 | 0.5 | 6 | 1E-04 | 1.0 | 3E-07 | 5 | 0.8 | 6 | 2E-04 | 1.0 | 8E-06 | 5 | 1.0 | 6 |
| 'YVKCQE' | 780 - 785 | 7E-03 | 0.6 | 8E+00 | 1 | 1.0 | 4 | 6E-08 | 0.9 | 2E-05 | 3 | 1.0 | 2 | 3E-14 | 1.0 | 1E-10 | 3 | 1.0 | 2 | 3E-14 | 1.0 | 7E-11 | 3 | 1.0 | 2 |
| 'YVKCQERVSA' | 780 - 789 | 4E+00 | 0.9 | 8E-02 | 8 | 0.6 | 1 | 1E-05 | 1.0 | 2E-10 | 4 | 0.3 | 5 | 1E-05 | 1.0 | 4E-10 | 4 | 0.4 | 5 | 4E-05 | 1.0 | 9E-07 | 4 | 0.7 | 5 |
| 'YVKCQERVSAL' | 780 - 790 | 1E-02 | 0.7 | 4E+00 | 1 | 1.0 | 9 | 5E-12 | 0.6 | 3E-05 | 6 | 1.0 | 4 | 6E-07 | 0.8 | 2E-05 | 6 | 1.0 | 4 | 1E-07 | 0.9 | 2E-05 | 6 | 1.0 | 4 |
| 'LYKNPRE' | 790 - 796 | 8E-03 | 0.6 | 5E+00 | 1 | 1.0 | 4 | 2E-14 | 0.5 | 4E-01 | 3 | 1.0 | 2 | 3E-14 | 0.4 | 2E-01 | 3 | 1.0 | 2 | 2E-14 | 0.7 | 5E-01 | 3 | 1.0 | 2 |
| 'LYKNPREWTRM' | 790 - 800 | 1E-02 | 0.7 | 3E+00 | 1 | 1.0 | 8 | 1E-12 | 0.2 | 3E-04 | 5 | 1.0 | 4 | 2E-12 | 0.2 | 3E-04 | 5 | 1.0 | 4 | 9E-14 | 0.2 | 4E-04 | 5 | 1.0 | 4 |
| 'YKNPRE' | 791 - 796 | 8E-03 | 0.5 | 7E+00 | 1 | 1.0 | 3 | 1E-04 | 1.0 | 1E+00 | 2 | 1.0 | 2 | 4E-05 | 0.7 | 9E-01 | 2 | 1.0 | 2 | 3E-14 | 1.0 | 1E+00 | 2 | 1.0 | 2 |
| 'YKNPREWTRM' | 791 - 800 | 1E-02 | 0.6 | 3E+00 | 1 | 1.0 | 7 | 4E-13 | 0.2 | 2E-04 | 5 | 1.0 | 3 | 5E-14 | 0.2 | 2E-04 | 5 | 1.0 | 3 | 2E-13 | 0.2 | 3E-04 | 5 | 1.0 | 3 |
| 'VIRNIATSGKF' | 801 - 811 | 1E-02 | 0.7 | 3E+00 | 1 | 1.0 | 9 | 4E-08 | 0.4 | 5E-06 | 5 | 1.0 | 4 | 1E-10 | 0.3 | 3E-07 | 5 | 1.0 | 5 | 2E-07 | 0.3 | 7E-07 | 6 | 1.0 | 4 |
| 'VIRNIATSGKFSSD' | 801 - 814 | 2E+00 | 1.0 | 7E+00 | 9 | 0.7 | 4 | 3E-05 | 0.9 | 3E-06 | 4 | 0.3 | 9 | 2E-14 | 0.2 | 2E-04 | 11 | 1.0 | 2 | 2E-14 | 0.2 | 3E-04 | 11 | 1.0 | 2 |
| 'VIRNIATSGKFSSDRTIAQ' | 801 - 819 | 3E+00 | 1.0 | 1E-02 | 17 | 0.7 | 1 | 5E-06 | 1.0 | 1E-12 | 8 | 0.2 | 10 | 9E-06 | 1.0 | 9E-13 | 8 | 0.3 | 10 | 1E-05 | 1.0 | 1E-07 | 8 | 0.3 | 10 |
| 'VIRNIATSGKFSSDRTIAQY' | 801 - 820 | 2E-02 | 0.7 | 3E+00 | 1 | 0.9 | 18 | 3E-06 | 0.3 | 2E-05 | 10 | 1.0 | 9 | 2E-06 | 0.3 | 2E-05 | 10 | 1.0 | 9 | 1E-06 | 0.4 | 1E-05 | 10 | 1.0 | 9 |
| 'IATSGKFSSDRTIAQ' | 805 - 819 | 3E+00 | 1.0 | 1E-02 | 13 | 0.7 | 1 | 5E-06 | 1.0 | 4E-11 | 6 | 0.2 | 8 | 5E-06 | 1.0 | 2E-11 | 6 | 0.2 | 8 | 1E-05 | 1.0 | 2E-11 | 6 | 0.3 | 8 |
| 'TSGKFSSD' | 807 - 814 | 4E+00 | 1.0 | 5E+00 | 5 | 0.5 | 2 | 8E-07 | 0.1 | 3E-04 | 6 | 1.0 | 1 | 2E-14 | 0.1 | 2E-04 | 6 | 1.0 | 1 | 9E-08 | 0.1 | 3E-05 | 6 | 1.0 | 1 |
| 'SGKFSSD' | 808 - 814 | 5E-01 | 0.6 | 5E+00 | 1 | 1.0 | 5 | 6E-08 | 0.2 | 6E-06 | 3 | 1.0 | 3 | 2E-08 | 0.2 | 5E-06 | 3 | 1.0 | 3 | 6E-08 | 0.3 | 6E-06 | 3 | 1.0 | 3 |
| 'SGKFSSDRTIAQ' | 808 - 819 | 1E-02 | 0.7 | 3E+00 | 1 | 1.0 | 10 | 4E-14 | 0.1 | 3E-08 | 6 | 1.0 | 5 | 4E-14 | 0.1 | 4E-08 | 6 | 1.0 | 5 | 2E-14 | 0.1 | 2E-08 | 6 | 1.0 | 5 |
| 'SSDRTIAQ' | 812 - 819 | 1E-02 | 0.6 | 3E+00 | 1 | 1.0 | 6 | 1E-09 | 0.1 | 2E-08 | 3 | 1.0 | 4 | 4E-14 | 0.2 | 1E-06 | 4 | 1.0 | 3 | 2E-06 | 0.4 | 1E-05 | 4 | 1.0 | 3 |
| 'YAREIW' | 820 - 825 | 8E-03 | 0.4 | 2E+00 | 1 | 1.0 | 4 | 5E-12 | 0.2 | 5E-06 | 3 | 1.0 | 2 | 4E-12 | 0.2 | 5E-06 | 3 | 1.0 | 2 | 5E-12 | 0.3 | 1E-05 | 3 | 1.0 | 2 |
| 'YAREIWGVE' | 820 - 828 | 3E-03 | 0.6 | 1E+00 | 1 | 1.0 | 7 | 2E-14 | 0.3 | 1E-03 | 4 | 1.0 | 4 | 2E-14 | 0.3 | 7E-04 | 4 | 1.0 | 4 | 2E-14 | 0.4 | 1E-03 | 4 | 1.0 | 4 |
| 'YAREIWGVEPSRQRLPAPDEKIP' | 820 - 842 | 2E+00 | 0.7 | 1E+00 | 5 | 1.0 | 13 | 1E-01 | 0.2 | 6E-08 | 10 | 0.6 | 8 | 5E-01 | 0.5 | 1E-04 | 10 | 0.6 | 8 | 5E-01 | 0.5 | 1E-04 | 10 | 0.7 | 8 |
| 'AREIWGVEPSRQRLPAPDEKIP' | 821 - 842 | 2E+00 | 0.7 | 1E+00 | 4 | 1.0 | 13 | 1E-01 | 0.2 | 1E-06 | 11 | 0.6 | 6 | 5E-01 | 0.4 | 2E-04 | 11 | 0.6 | 6 | 5E-01 | 0.4 | 2E-04 | 11 | 0.7 | 6 |
| 'GVEPSRQRLPAPDEKIP' | 826 - 842 | 3E+00 | 0.8 | 1E+00 | 5 | 0.8 | 7 | 2E-01 | 0.4 | 1E-02 | 8 | 0.6 | 4 | 1E+00 | 0.7 | 8E-02 | 6 | 0.9 | 5 | 1E+00 | 0.7 | 8E-02 | 6 | 1.0 | 6 |
| 'PSRQRLPAPDEKIP' | 829 - 842 | 1E+02 | 0.7 | 2E+00 | 0 | 0.2 | 10 | 9E-05 | 0.5 | 1E-01 | 1 | 0.5 | 9 | 6E-02 | 0.8 | 1E+00 | 4 | 0.4 | 6 | 1E+00 | 0.6 | 1E-01 | 5 | 1.0 | 5 |

1. Bai Y, Milne JS, Mayne L, Englander SW (1993) Primary structure effects on peptide group hydrogen exchange. *Proteins* 17(1):75–86.
2. Gutfreund, H. (1969), Resolution of optical and sampling methods. Methods in Enzymology, 16, 229-249.
3. Kemmer, G. and Keller, S. 2010. Nonlinear least-squares data fitting in Excel spreadsheets. Nature protocols. 5(2),pp.267–281.
